# Beyond species means – the intraspecific contribution to global wood density variation

**DOI:** 10.1101/2025.08.25.671896

**Authors:** Fabian Jörg Fischer, Jérôme Chave, Amy Zanne, Tommaso Jucker, Alex Fajardo, Adeline Fayolle, Renato Augusto Ferreira de Lima, Ghislain Vieilledent, Hans Beeckman, Wannes Hubau, Tom De Mil, Daniel Wallenus, Ana María Aldana, Esteban Alvarez-Dávila, Luciana F. Alves, Deborah M. G. Apgaua, Fátima Arcanjo, Jean-François Bastin, Andrii Bilous, Philippe Birnbaum, Volodymyr Blyshchyk, Joli Borah, Vanessa Boukili, J. Julio Camarero, Luisa Casas, Roberto Cazzolla Gatti, Jeffrey Q. Chambers, Ezequiel Chimbioputo Fabiano, Brendan Choat, Georgina Conti, Will Cornwell, Javid Ahmad Dar, Ashesh Kumar Das, Magnus Dobler, Dao Dougabka, David P. Edwards, Robert Evans, Daniel Falster, Philip Fearnside, Olivier Flores, Nikolaos Fyllas, Jean Gérard, Rosa C. Goodman, Daniel Guibal, L. Francisco Henao-Diaz, Vincent Hervé, Peter Hietz, Jürgen Homeier, Thomas Ibanez, Jugo Ilic, Steven Jansen, Rinku Moni Kalita, Tanaka Kenzo, Liana Kindermann, Subashree Kothandaraman, Martyna Kotowska, Yasuhiro Kubota, Patrick Langbour, James Lawson, André Luiz Alves de Lima, Roman Mathias Link, Anja Linstädter, Rosana López, Cate Macinnis-Ng, Luiz Fernando S. Magnago, Adam R. Martin, Ashley M. Matheny, James K. McCarthy, Regis B. Miller, Arun Jyoti Nath, Bruce Walker Nelson, Marco Njana, Euler Melo Nogueira, Alexandre Oliveira, Rafael Oliveira, Mark Olson, Yusuke Onoda, Keryn Paul, Daniel Piotto, Phil Radtke, Onja Razafindratsima, Tahiana Ramananantoandro, Jennifer Read, Sarah Richardson, Enrique G. de la Riva, Oris Rodríguez-Reyes, Samir G. Rolim, Victor Rolo, Julieta A. Rosell, Roberto Salguero-Gómez, Nadia S. Santini, Bernhard Schuldt, Luitgard Schwendenmann, Arne Sellin, Timothy Staples, Pablo R Stevenson, Somaiah Sundarapandian, Masha T van der Sande, Bernard Thibaut, David Yue Phin Tng, José Marcelo Domingues Torezan, Boris Villanueva, Aaron Weiskittel, Jessie Wells, S. Joseph Wright, Kasia Zieminska

## Abstract

Wood density is central for estimating vegetation carbon storage and a plant functional trait of great ecological and evolutionary importance. However, the global extent of wood density variation is unclear, especially at the intraspecific level.

We assembled the most comprehensive wood density collection to date (GWDD v.2), including 109,626 records from 16,829 plant species across woody life forms and biomes. Using the GWDD v.2, we explored the sources of variation in wood density within individuals, within species, and across environmental gradients.

Intraspecific variation accounted for up to 15% of overall wood density variation (sd = 0.068 g cm^-3^). Sapwood densities varied 50% less than heartwood densities, and branchwood densities varied 30% less than trunkwood densities. Individuals in extreme environments (dry, hot, acidic soils) had higher wood density than conspecifics elsewhere (+0.02 g cm^-3^, ∼4% of the mean). Intraspecific environmental effects strongly tracked interspecific patterns (r = 0.83) but were only 20–30% as large and varied considerably among taxa.

Individual plant wood density was difficult to predict (RMSE > 0.08 g cm^-3^; single-measurement R^2^ = 0.59). We recommend (i) systematic within-species sampling for local applications, and (ii) expanded taxonomic coverage combined with integrative models for robust estimates across ecological scales.

## 1. Introduction

Wood density, the ovendry mass of wood (g) over its fresh volume (cm^-3^), is an important plant functional trait in ecology and global change studies. Accurate species-level averages of wood density are needed for unbiased estimation of aboveground carbon in vegetation (Phillips et al., 2019). Moreover, wood density defines one of the main axes of global plant trait variation (Díaz et al., 2016). Generally, high-wood density species are less susceptible to mechanical, hydraulic or biotic stress (Chave et al., 2009), experience low mortality at the expense of growth (King et al., 2006; Kraft et al., 2010), and decompose more slowly (Hérault et al., 2010). Wood density therefore displays distinct patterns across successional and environmental gradients (Poorter et al., 2019; Šímová et al., 2018) and is a key factor in the prediction of the global carbon cycle and terrestrial ecosystem dynamics (Sakschewski et al., 2015).

Wood density varies widely among woody plants, from species with incredibly low-density wood (∼0.10 g cm^-3^ in *Jacaratia spinosa* (Aubl.) A.DC) to species such as lignum vitae (*Guaiacum sanctum* L.) whose wood is denser than water (∼1.05 g cm^-3^) and therefore sinks at any moisture level. However, wood density also varies within and among individuals of the same species (Anderegg et al., 2021; Fajardo et al., 2022; Yang et al., 2023). Intraspecific variation in traits provides an imprint of how organisms react to changes in their environment through adaptation and morphological plasticity (Bolnick et al., 2011; Girard-Tercieux et al., 2023; Moran et al., 2016) and can play an important role in ecosystem functioning (Des Roches et al., 2018). Better knowledge of shifts with ontogeny and along environmental gradients would improve vegetation models (Berzaghi et al., 2020), predict how species ranges shift in response to climatic change and disturbance (Anderegg & HilleRisLambers, 2016), and create more robust wood density maps for assessment of functional diversity and vegetation carbon stocks (Boonman et al., 2020; Sæbø et al., 2022).

Radial changes within tree trunks and branches are a common source of intraspecific variation in wood density, usually interpreted as a reflection of hydraulic and mechanical changes during ontogeny (Wiemann & Williamson, 1988; Woodcock & Shier, 2002). In plant species with low wood density near the pith, wood density often increases towards the outer trunk layers, while the opposite may occur in plants with high near-pith wood density (González-Melo et al., 2022; Hietz et al., 2013; Longuetaud et al., 2017; Plourde et al., 2015; Woodcock & Shier, 2002), although there are many exceptions to this pattern (Bastin et al., 2015; Osazuwa-Peters et al., 2014). A link between radial variation in wood density and plant ecological strategies has also been suggested: pioneer plants have low density wood and grow fast early on, but invest in denser tissues as individuals mature. In contrast, shade-tolerants build dense tissues initially, but may invest more in diameter growth than tissue density when reaching the canopy (Bastin et al., 2015; Woodcock & Shier, 2002). However, we do not know how consistent and important these patterns are at global scales. Radial changes in wood density do not always map onto ecological strategies (Hietz et al., 2013), may be influenced by deposition of chemical compounds in heartwood (e.g., non-structural, secondary metabolites known as “extractives”, Lehnebach et al., 2019), and vary among conspecific individuals or even within individuals (Osazuwa-Peters et al., 2014).

Wood density also varies along the hydraulic pathway and across plant organs within an individual, another source of intraspecific variation (Longuetaud et al., 2017; Momo et al., 2020; Schuldt et al., 2013). In trees, for example, wood density has been hypothesized to increase from trunks to branches, because high density should provide more benefits to mechanical stability in horizontal branch than vertical trunk wood (Anten & Schieving, 2010; van Casteren et al., 2012). However, while branch and trunk wood densities are generally tightly correlated with one another, there is little agreement on whether branches are more (Billard et al., 2020; Dibdiakova & Vadla, 2012; Fajardo, 2018; Fegel, 1941) or less dense than trunks (He & Deane, 2016; Sarmiento et al., 2011; Swenson & Enquist, 2008), and there is also variation within trunks and branches (Schuldt et al., 2013; Terrasse et al., 2021). A confounding factor may be that wood density varies less in branches than in trunks: data from temperate ecosystems show that wood density increases from trunk to branches in species with low-density trunk wood and shows the opposite pattern in species with high-density trunk wood (MacFarlane, 2020). It is unclear whether this pattern generalizes across biomes and also whether it reflects different functional requirements of branches and trunks. However, it has been hypothesized that trunk-branch gradients simply reflect radial ontogenetic patterns between juvenile and mature wood, since more distal organs are younger and contain higher fractions of sapwood (Gartner, 1995).

Wood density also varies across individuals of the same species due to differences in genetics and environments (Zobel & van Buijtenen, 1989). As environmental conditions become more extreme (drier, less fertile, more shaded, more windy), species are expected to grow more slowly and invest more resources in dense and stress-resistant tissues (Chave et al., 2009). Similar effects are expected within species (Anderegg et al., 2021). For example, individuals that build tissues with narrow conduits and thick fibre and conduit walls should be more resistant to embolism (Hacke et al., 2001; Olson et al., 2020). In contrast, in warm, fertile and frequently disturbed environments with high plant turn§over, fast-growing individuals with low wood density are expected to be more competitive and successful (Muller-Landau, 2004; Yang et al., 2023). However, genetic control over wood density is high, suggesting that the variation among individuals of the same species is limited (Zobel & Jett, 1995). Wood density variation may also be limited by covariation with other traits, and trade-offs between different wood functions (Anderegg & HilleRisLambers, 2016; Ziemińska et al., 2013). For example, low-density wood can sometimes be beneficial even in harsh conditions, as low-density species tend to have greater water storage and capacitance (Ziemińska et al., 2020), a potential advantage in dry environments.

Overall, theory, empirical observations, and common sense predict that wood density varies predictably within species. However, while many studies find that intraspecific variation is predictable (Anderegg et al., 2021; Anderegg & HilleRisLambers, 2016; Farias et al., 2023), just as many do not (Fajardo, 2018; Richardson et al., 2013; Rosas et al., 2019; Umaña & Swenson, 2019). Intraspecific variation is generally smaller than interspecific variation (Osazuwa-Peters et al., 2014), so it is easily confounded with measurement errors and methodological differences in how wood density is determined (Barbosa & Fearnside, 2004; Jati et al., 2014; Vieilledent et al., 2018; Williamson & Wiemann, 2010). To date, the largest global wood density collections, including the GWDD v.1 (Zanne et al., 2009), do not systematically record the tissue types and plant organs where measurements were taken. As a result, there is a fundamental lack of knowledge about the extent of intraspecific variation in wood density, its determinants within and among individuals, and its implications for ecological models and carbon estimates in woody ecosystems. In particular, we lack practical guidelines as to when to exhaustively measure it versus when it can be safely ignored.

Here, we introduce a substantially updated and improved version of the Global Wood Density Database (GWDD v.2), which more than doubles the taxonomic coverage of the original database (from 7,555 to 16,829 taxonomically resolved species), increases the number of records from 16,468 to 109,626, and, as available, includes a detailed description of where and how measurements were taken within and across individuals. Using these data, we addressed the following questions: 1. How large is intraspecific variation in wood density? 2. How much of this intraspecific variation can be explained by differences in wood density among plant organs? 3. How much of intraspecific variation can be explained by environmental factors, such as temperature, water deficit, wind speeds and soil fertility? Based on our results, we also 4. make a number of recommendations to effectively incorporate variation among and within individual plants into models to improve predictions of wood density.

## 2. Methods

### 2.1 Assembling the Global Wood Density Database v.2

The GWDD v.2 is a substantial update of the GWDD v.1, an open-access database that consisted of a list of 16,468 tissue density values from ca. 7,500 species. It was assembled from more than 200 sources and included a taxonomic identifier, a region where the measurement was taken, and a literature reference (Chave et al., 2009; Zanne et al., 2009). Since its publication, many new wood density values and large trait databases have been published, and improved methodologies have been developed to improve consistency across studies (Cuny et al., 2025; Farias et al., 2020; Langbour et al., 2019; Radtke et al., 2023; Vieilledent et al., 2018; Williamson & Wiemann, 2010). We used this as an opportunity to expand the database and create a new, improved version.

First, we critically re-examined the original database and updated 42% of entries (n = 6,968, details in Supplementary S1). Second, we included additional wood density measurements from published and unpublished sources (cf. Supplementary S1). We put particular emphasis on previously undersampled biomes, such as dry forests, savannas, and the species-rich tropics. We also included individual measurements instead of aggregated values and created a comprehensive documentation. The new database contains 45 attributes that report the original values, sampling techniques, and data transformation methods (Table S1). The database is available online on Zenodo (doi: 10.5281/zenodo.16919509, embargoed until acceptance for publication).

#### Wood density definition and conversion factors

In the GWDD v.2 and throughout this study, wood density is defined as ‘basic’ wood density, the mass of an oven dried wood sample divided by its fresh volume (g cm^-3^). Wood density

thus measures the dry mass contained in the wood volume of live plants and is an indicator of a plant’s investment in woody tissues. When normalized by the density of water (1 g cm^-3^), it is also referred to as “wood specific gravity” (g g-1; Williamson & Wiemann, 2010), but throughout our analyses, we use the term “wood density”. When assembling the GWDD v.2 and in all following analyses, we also included the tissue densities of tree-like monocots without secondary growth, as this was consistent with common inventory protocols (Condit, 1998; The SEOSAW Partnership, 2021) and global tree databases (e.g., Beech et al., 2017). In total, monocots contributed 186 records from 91 species most of which were either Arecaceae (65) or Asparagaceae (18). Since these monocots amounted to less than 0.2% of the total records, their inclusion had a negligible effect on results.

Different wood density definitions are available in the literature. Green density, the fresh mass of wood divided by fresh volume, reflects actual plant growing conditions (Niklas & Spatz, 2010). In the timber industry, a relevant quantity is airdry wood density, the mass of wood divided by its volume, with both measured at ambient air moisture (∼10-15%), reflecting properties of wood in the conditions in which it is used (Détienne & Chanson, 1996; FPL, 1999). Ovendry density, i.e., dry mass over dry volume, has also been reported in the literature (Deklerck et al., 2019), and dendrochronological studies often derive correlates of wood density variation within and between tree rings with X-ray techniques (Jacquin et al., 2017). Fortunately, air- and ovendry densities can each be converted into basic density through physical conversion factors (Brown, 1997; Ilic et al., 2000; Sallenave, 1971), and recent research has shown that this can be done with as little error as 0.015 g cm^-3^ (ca. 2.5% of typical mean wood density, Vieilledent et al., 2018). These factors were also applied in the construction of the GWDD v.2 to maximize the taxonomic and geographic coverage of wood density. Converted values and the source quantity were recorded in the columns *value_reference* and *quantity_reference*, the conversion factor in *wsg_conversion*, and the derived basic wood density value as *wsg* (“wood specific gravity”).

#### Aggregation levels

The GWDD v.2 provides extensive information on where and how records were obtained, which was not available in the GWDD v.1. These new variables include *site*, an informal description of the measurement site, *longitude* and *latitude* in the decimal system, and *country*. The attribution to a *region* has been revised since GWDD v.1 to better reflect geographical variation (Table S1). The database also contains information on sample type (*type_sample* for “core” or “disk”), measurement location within plants (*location_sample* for “trunk”, “branch” or “root”, for example, and *type_tissue* for “heartwood”, “sapwood” or “bark”), whether a particular wood density value is the mean value of multiple individual plants (*plant_agg*), and how many individuals were aggregated (*plants_sampled*). If several measurements were available for the same individual, they were recorded with the same *id_plant*. Direct estimates of variation around mean values were not included in the GWDD v.2, as they were not consistently reported in the literature. Where detailed measurement information was not available, attributes were left empty (“NA”). These samples were excluded from analyses of intraspecific wood density variation in this study.

#### Taxonomic name resolution

Taxonomic names were newly standardized via the *WorldFlora* R package (Kindt, 2020) and the June 2023 version of the *World Flora Online* (WFO) database (The World Flora Online Consortium et al., 2023). Taxon names were converted in the field *species_reference_canonical*, including infraspecific assignations (e.g., variety, subspecies, hybridization) and resolved via the default fuzzy matching in the *WorldFlora* package. Taxonomic authorities were not included as inputs for the matching, as they were inconsistently reported in the source data. For unmatched taxa and anything beyond a missing, added or switched letter, the matching was repeated without infraspecific assignations. In the case of multiple matches, we chose the default value provided by *World Flora Online* (called “smallest id”). Any remaining taxa were manually corrected. We also extracted information on plant families from the WFO database and reported it in the GWDD v.2. Overall, only 34 entries (22 genera) could not be matched to any family.

Taxonomies are constantly updated to resolve ambiguities in species definitions, and sometimes, taxonomic resolution leads to a reclassification of species as subspecies and varieties (or vice versa). For simplicity, we recorded the entire scientific name provided by *WorldFlora* in the GWDD v.2’s *species* column, including infraspecific epithets. We also used this species definition for analyses in our study to provide the most conservative estimates of intraspecific variation. However, preliminary tests revealed that only a negligible portion of the analysed species had infraspecific epithets (∼1% of all species with individual level wood density records, <1% among high-quality records), so this choice had no discernable effect on results.

### 2.2. Intraspecific wood density variation

#### Coverage of taxonomic and geographic variation

Overall, we assembled more than 100,000 records from ca. 17,000 plant species in the GWDD v.2. We assessed the representativeness of the species included in the GWDD v.2 with regard to the number of woody taxa worldwide estimated by assuming that 45% of the flowering plants are woody (FitzJohn et al., 2014), and the total number of flowering plants is ca.

400,000 (Enquist et al., 2019). We also used a verified list of known tree species (GlobalTreeSearch 1.7; Beech et al., 2017), resolved via *World Flora Online* for consistency, to assess which percentage of tree species in each country had a corresponding wood density estimate in the GWDD v.2. Intraspecific coverage was assessed through the number of records per species.

#### Statistical analysis of intraspecific wood density variation

To assess the extent and drivers of intraspecific wood density variation, we examined measurements from multiple sites per species, multiple individuals per site and multiple measurement locations per individual. For some species, the database contains multiple samples of individuals, but only from a single site, while for others, the database contains samples from multiple sites, but each with a single individual. To address this issue, we created subsets of the GWDD v.2 for each question, and alternative modeling strategies to ensure robustness of results (see Table S3 for an overview of models and subset sizes). Throughout, ‘intraspecific variation’ refers to cases where biological variation can be confidently separated from measurement errors. For example, tissue type (heartwood/sapwood) and environmental factors should affect biological variation, not measurement error. By contrast, when variation cannot be attributed to a specific factor, intraspecific variation plus measurement error are referred to as the ‘(residual) wood density distribution’.

Across all datasets, we removed records not identified to species level (“genus” in the *rank_taxonomic* column of the database), samples consisting of bark (“bark” in the *type_tissue* column of the database), and records from experiments (identified by the words “fertilizer” or “treatment” in *experiment_design*) or from plantations (recorded as “plantation” in *type_forest)*. We also excluded root samples (“root” in the *location_sample* column) due to small sample sizes.

Unless otherwise stated, all datasets were analyzed using mixed effects models with varying random slopes and intercept terms and qualitative explicative variables (e.g., “sapwood” vs. “heartwood”) coded as numerical variables (0, 1). Models were fitted in the R environment (R Core Team, 2023), both with a Bayesian approach – using the *brms* package (Bürkner, 2018) and the STAN software (Carpenter et al., 2017) – and the maximum likelihood framework using the *lme4* package (Bates et al., 2015, ‘bobyqa’ optimizer). The Bayesian approach is the default, due to flexibility in model construction, regularization through priors and full propagation of uncertainty. All models were assessed for convergence using standard diagnostics (R-hat <= 1.02) and posterior predictive checks (further details in S3, Table S3 and Fig. S1). Throughout, we also report *lme4* estimates, as they are less expensive computationally and thus more readily used in practice, especially when relying on large databases.

#### Quantification of intraspecific variation in wood density

To assess the overall extent of intraspecific variation in wood density, we first partitioned total wood density variance and its components. We fitted a model with random effects for species nested within genera, genera nested within families, and a crossed random effect for methodological bias (the bibliographic reference or “source”, model M1, Table S3). We computed variance as the sum of variances across levels plus residual variance. To assess robustness, we also fitted a separate model without the “source” effect (Table S3, M2) and restricted the analysis to species with ≥ 3 individuals per species, ≥ 3 species per genus, ≥ 3 genera per family (n = 49,991, n_species_ = 2,735; Tables M3-4).

Second, to assess the contribution of different sources of intraspecific variation to its overall extent, we partitioned the intraspecific variance of wood density. We did so by fitting models with random effects for individuals nested within sites, and sites nested within species, each time with and without a crossed effect for measurement source (formulas in Table S3, M5-8). We restricted the analysis to subsets of species present in at least k sites, one site with at least k individuals, and one individual with at least k measurements (first setting k = 2 and then repeating for k = 3). Sites were defined as the collection of data within the same 1 km^2^ grid pixel. This approach mirrored the resolution of the climate input data and accounted for geolocation uncertainty (e.g., rounding of longitude and latitude values to 2 decimals). Data subsets comprised n = 19,246 (n_individual_ = 14,373, n_site_ = 1,270, n_species_ = 147) records for k=2, and n = 2,494 (n_individual_ = 1,052, n_site_ = 233, n_species_ = 35) for k=3.

The shape of intraspecific wood density distributions was assessed through the direct modeling of the residual variance parameter σ (Table S3, model M1, cf. Supplementary S3) and by comparing how well normal and lognormal distributions fitted each species’ wood density distribution (Shapiro-Wilkes test for untransformed vs. log-transformed wood densities with p < 0.05). To account for methodological differences and report biological variation, we always subtracted the random “source” effect before reporting wood density variation.

#### Estimating variation in wood density within individuals

Variation in wood density within individuals was assessed by subsetting to species with measurements for both sapwood and heartwood (n = 679, n_species_ = 150), or for both trunkwood and branchwood (n = 48,494, n_species_ = 2,018). The fitted models (Table S3, M9-10) had fixed effects for either sapwood (0 for heartwood, 1 for sapwood) or branchwood (also 0/1), random intercepts and slopes at species level, and a crossed random effect for measurement source. To assess whether wood density followed predictable gradients from the pith outwards to the bark and from the trunk upward to the branches, we calculated species means for each woody component and checked whether the slopes of major axis regressions (lmodel2 package; Legendre, 2018) of sapwood vs. heartwood and branchwood vs. trunkwood densities were < 1 (indicating a decrease) or > 1 (indicating an increase, Table S3, M11-12).

To assess robustness and potential confounding effects, such as higher sapwood fraction in branches or variation in sample sizes among taxa, we repeated the analysis with a subset of branchwood and trunkwood densities taken entirely from sapwood (n = 1,193, n_species_ = 523, M13), with a higher-quality dataset (species with >= 5 measurements both for branches and trunks, selecting 5 random measurements from each, n_species_ = 189, M14), and a subset of records where both branch and trunk samples were taken from the same individuals (n = 3,527, n_species_ = 145, M15, fitted at individual plant level, taking a random sample from both trunk and branch).

#### Environmental predictors of intraspecific wood density variation

To examine wood density variation across environmental gradients, we used species sampled from at least two sites, and paired them with the following bioclimatic layers that represent multiple axes of plant environmental gradients including extremes: annual mean temperature (°C), site water balance (kg m^-2^ yr^-1^, but with the sign reversed to indicate water deficit) and mean wind speed (m s^-1^) at 1 km resolution. We used the data from the CHELSA/BIOCLIM+ climatology 1981-2010 (Brun et al., 2022; Karger et al., 2017), but also repeated the analysis with products from the TerraClimate climatology 1981-2010, relying on climatic water deficit (mm) instead of site water balance (Abatzoglou et al., 2018). While site water balance is a general measure of the availability of water to plants (Brun et al., 2022), climatic water deficit directly measures drought stress as the difference between potential and actual evapotranspiration. We also included the following soil layers: sand fraction (g kg^-1^), pH (unitless) and cation exchange capacity (in mmol(c) kg^-1^), based on the *soilgrids* product (Hengl et al., 2017). By default, we report only effects that remained qualitatively consistent across analyses methods (no shift in sign). For simplicity, effect sizes are always taken from the full dataset using CHELSA/BIOCLIM+ predictors.

We matched predictors to wood density observations and assessed environmental effects both among and within species with the following model (cf. Table S3, M16-19): 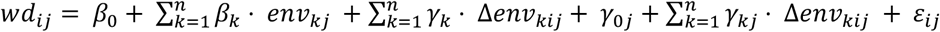 with *ε*_*ij*_ ∼ *N*(0, σ^2^). Here, *n* is the number of individuals, *wd*_*ij*_ is the wood density of individual *i* belonging to species *j, env*_*kj*_ the environmental variable *k* averaged across all individuals from species *j* and Δ*env*_*kij*_ the same environmental predictor *k*, but group-centred around the species mean value, and ε_*ij*_ is the error. The model thus partitions environmental effects into interspecific effects, where *env*_*kj*_ is the typical environment of species *j*, and intraspecific effects, where Δ*env*_*kij*_ represents how much the individual *i* deviates from the species mean environment. The model is equivalent to a standard linear-mixed effects model with random slopes and intercepts for species, but with added group-level predictors *env*_*kj*_ that control for predictable variation among species (Table S3, models M16-17, Bafumi & Gelman, 2011). *β*_0_ is the overall intercept, *β*_*k*_ and γ_*k*_ are fixed-effect parameters, and γ_0*j*_ and γ_*kj*_ are random intercept and slope parameters for species *j*, respectively. We chose this model as it allows for partitioning of intra- and interspecific effects and is more robust when predictors vary systematically with grouping factors (Bafumi & Gelman, 2011).

Since some species may cover only a narrow environmental range, we repeated the analysis with a subset of species for which at least one environmental factor covered a large range (Table S3, n = 30,128, n_species_ = 692, M18-19), defined via TerraClimate and 5 km *soilgrids* as the 90^th^ percentiles of all species’ ranges (Δ_Temperature_ >= 8°C, Δ_Water Deficit_ >= 450 mm, Δ_Wind speed_ >= 1.8 m s^-1^, Δ_Sand content_ >= 300 g kg^-1^, Δ_pH_ >= 1.5 or Δ_Cat. exch. cap._ >= 165 mmol(c) kg^-1^). To assess the consistency of predictors across biomes, we fitted separate models for tropical species (≥ 3 measurements in the tropics, n = 8,783, n_species_ = 700, M20-21) and extratropical species (≥ 3 measurements outside the tropics, n = 26,437, n_species_ = 247, M22-23).

In all models, intra- and interspecific effects were examined on the same standardized scale. However, as environmental gradients among species were larger than among individuals of the same species, we also tested the rescaling of effect sizes to realized environmental ranges for the tropical and extratropical subsets, i.e. standardizing *after* separation of intra- and interspecific effects by their respective standard deviations. We fitted additional models to gymnosperms only (Table S3, n = 12,089, n_species_ = 59, M24-25) to assess the stability of global effects when sampling is reduced to a small number of anatomically diverging taxa.

#### The effect of intraspecific variation on wood density predictions

Since wood density measurements are destructive, samples are usually only taken from a subset of plants, from nearby conspecifics or from global databases. The samples are then used to predict the wood density of the remaining individuals, for example, by using species mean values, or, if those are not available, the average of wood densities from the same genus or plot (Flores & Coomes, 2011; Réjou-Méchain et al., 2017). If many traits have been measured, more complex imputation methods are available (Schrodt et al., 2015). However, most methods risk confounding intraspecific and interspecific variation and it is unclear what to do in edge cases, e.g., if 1-2 measurements from the same species are available, is it better to estimate an individual’s wood density by (i) directly using these values and averaging them, (ii) attributing the genus mean value, or (iii) combining both types of information, e.g., through taxonomic or phylogenetic hierarchical modelling? It is also (iv) unclear if local measurements should be weighted more strongly to account for environmental gradients in wood density.

To answer these questions, we first assessed the influence of intraspecific variation on species-level wood density estimates. We computed average wood densities for all species with more than five measurements (n_species_=1,667) and assessed how accurately these reference values could be predicted if species were not well sampled. To do so, we cycled through all 1,667 species, in turn removed either all species-specific measurements or all species-specific measurements except one or two randomly chosen ones, and then estimated the species’ mean wood density from the remaining data. The estimation was carried out with three models: i) the default approach of estimating wood density means from a genus average (when no species-specific measurements exist) or a species average (when one or two species-specific measurements exist), ii) a hierarchical model of wood density that included a nested taxonomic structure (*family / genus / species*) and nested random effects for sites within studies, and iii) the same model as in ii), but with an optional fixed effect for trunkwood vs. branchwood (Table S3, models M26-27). Model performance was estimated via root mean square error (RMSE, g cm^-3^) and R^2^ (Table S16).

Second, we tested how accurately we could predict the wood density of an individual plant depending on how well the species was sampled locally. To do so, we selected species with measurements from at least 3 sites, and where at least 4 of its individuals were measured at each of the 3 sites (n_species_ = 318). For each of the individuals, we then reduced the set of locally measured conspecifics to 0, 1, 2, or 3 (selecting random individuals where possible), and for each case tested how well the individual’s wood density could be inferred from the remaining data. We tested five models: i) a species average across the entire dataset, ii) a species average, but using only values measured as part of the same study, iii) a species average, but using only values measured locally, and iv) and v) the same hierarchical models as described above (Table S3, M26-27). Model performance was estimated via root mean square error (RMSE, g cm^-3^) and R^2^ (Table S17).

## 3. Results

### The Global Wood Density Database

To create the GWDD v.2, we assembled 109,626 wood density records from 166 countries, 617 primary sources and 17,262 taxa across all woody biomes and biogeographic realms. Of these taxa, we resolved 16,829 to species level. We also included information on aggregation levels, conversion factors and precise geographic location, as well as an additional 15,093 bark density records from 57 studies. The fully assembled GWDD v.2 contained 6.7x as many wood density records and 2.3x as many species as the GWDD v.1 (16,468 records across 7,453 accepted species). We estimated that the GWDD v.2 covered 10% of all woody species, 24% of all known tree species and 49% of gymnosperm species. Of the families with the most known tree species, Sapotaceae and Fabaceae were best represented in the database (38% and 34% coverage). In contrast, wood density estimates were rare in Arecaceae (6%, not true wood), Araliaceae (10%) and Melastomataceae (12%).

Out of all the GWDD v.2’s records, 83,502 (76%) were provided at individual plant level, and 58,675 entries (54%) were precisely geolocated. For 4,508 species, at least five wood density measurements were available, and combinations of branch-trunk wood measurements were available for 2,018 species (146 families). For 78 species, mostly in Pinaceae and Fagaceae, there were more than 100 wood density measurements. The majority (65%) of the wood density values were directly measured as basic wood density (71,746). The remaining values were converted from airdry (∼29%), and ovendry wood densities (∼6%). Geographic coverage varied widely, from a near-complete coverage of recorded tree species in high latitudes to < 50% in tropical regions (95% range: 27.3%; 98.5%, Fig. 1). An exception was West Africa where coverage of known tree species was ≥70%, albeit without precise geolocation for most records (cf. limited number of red dots in Fig. 1).

**Fig. 1:**
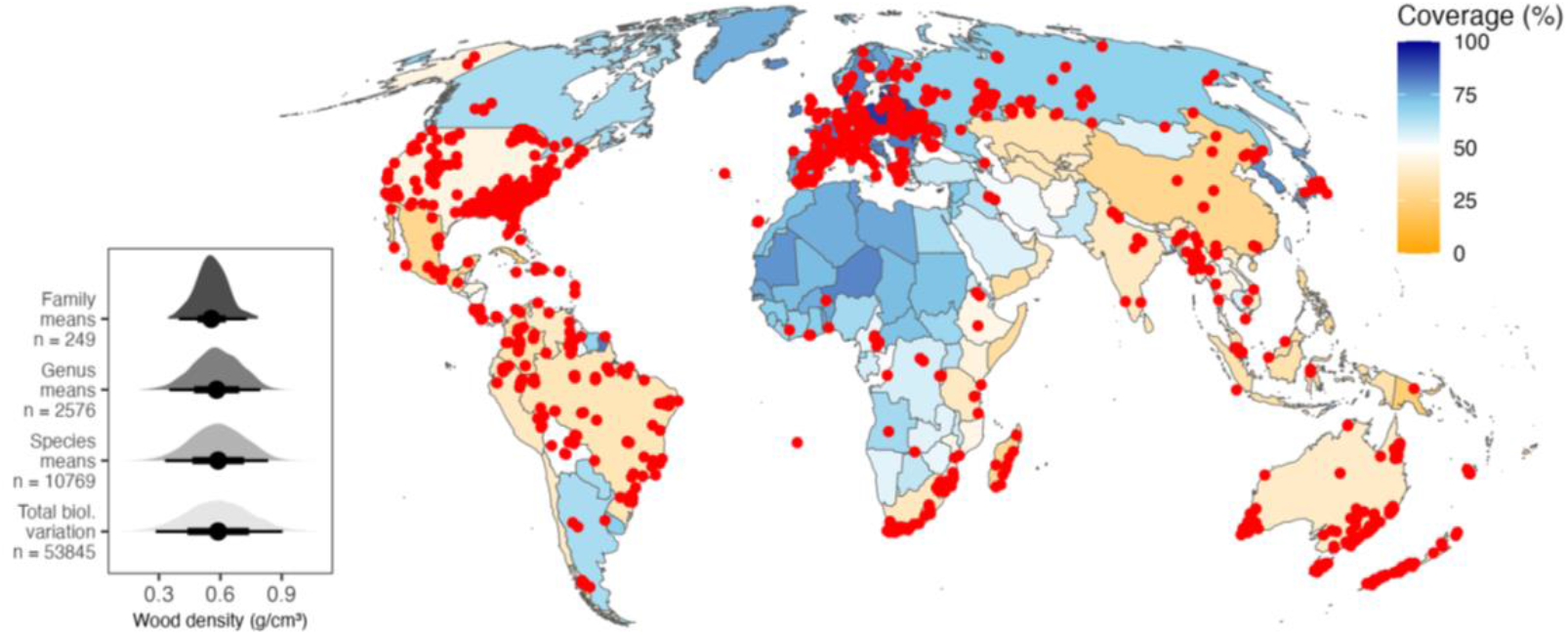
Geographic coverage of the Global Wood Density Database (GWDD) v.2. This figure separately maps 1) the percentage of each country’s species that could be matched to species in the GWDD v.2 (background colour gradient), and 2) all locations with explicitly geolocated wood density measurements in the GWDD v.2 (red dots). For the former, measurements did not have to be explicitly geolocated, meaning some countries had high species coverage, despite a lack of country-specific wood density measurements (e.g., West African countries). Conversely, some large, predominantly temperate countries (e.g., the U.S.) had many explicitly geolocated records, but also covered species-rich subtropical and tropical zones, which lowered their overall species coverage. Country boundaries follow *Natural Earth* (https://www.naturalearthdata.com/, last accessed 14 Dec 2023). The inset in the lower left corner shows the distributions of wood density at various levels of aggregation: distributions of family, genus and species means, as well as the full biological variation in wood density. To attenuate sampling biases, the distributions were based on five random draws from each species (with replacement). Biological variation was estimated by subtracting inferred methodological biases (“source” effect in model M1, Table S3) from the raw values.

### The extent and shape of intraspecific variation in wood density

Globally, wood density displayed a normal distribution with a mean of 0.56 g cm^-3^ (sd = 0.178 g cm^-3^, Fig. 1 inset). The majority of this variation (77%, sd = 0.156 g cm^-3^) was accounted for by variation at the taxonomic family (30%), genus (30%) or species levels (17%), with the rest attributed to study methodology (8%) or intraspecific variation and unknown errors (15%; 0.068 g cm^-3^, Fig. 1, inset, Fig. S2-3, Table S4). The intraspecific contribution was robust to model details (Table S4, Table S5). For well-sampled taxa, we further partitioned intraspecific variance, and found that wood density variation among sites exceeded variation among individuals within sites (sd = 0.025-0.042 g cm^-3^ vs. sd = 0.017-0.028 g cm^-3^, Table S6), but was small overall (ca. 20-30% of total intraspecific variation), and smaller than residual variation (variation within individuals + unknown measurement error, sd = 0.040-0.045 g cm^-3^, Table S5-6).

Intraspecific variation differed in extent and shape between species, with a few species varying much more than the others (Fig. S3, e.g., sd = 0.094 g cm^-3^ for *Quercus ilex*, compared to sd = 0.038 g cm^-3^ for *Quercus alba*). The distribution of wood density values of conspecifics was generally heavy-tailed, but there was no clear signal of skew, with the lognormal distribution more often rejected than the normal distribution (13.7% vs. 12.6%, Shapiro-Wilkes p < 0.05 for log-transformed and untransformed values).

### Wood density variation within individuals

Across plant tissue types, there were strong correlations between heartwood and sapwood densities (Pearson’s r = 0.78, Fig. 2a) and between trunkwood and branchwood densities (r = 0.67, Fig. 2b, Table S9). However, slopes were < 1 in both cases. There was a 50% reduction in the variance of sapwood compared to heartwood (0.144 vs. 0.197 g cm^-3^) and a ∼30% reduction in the variance of branchwood compared to trunkwood (0.120 g cm^-3^ vs. 0.145 g cm^-3^). When comparing only sapwood samples from trunks and branches, this effect was weaker (0.112 vs. 0.124 g cm^-3^, less than a 20% reduction in variance) and the correlation was stronger (r = 0.76, n = 523, Fig. 2c; for more analyses see Table S9, Figs. S5-S6). There was a weak average decrease from heartwood to sapwood of -0.001 g cm^-3^ and a stronger, but still small decrease of -0.023 g cm^-3^ from trunkwood to branchwood (Table S8). However, both effects varied strongly among species (slope sd = 0.060 g cm^-3^ for heartwood vs. sapwood and sd = 0.075 g cm^-3^ for trunkwood vs. sapwood).

**Fig. 2:**
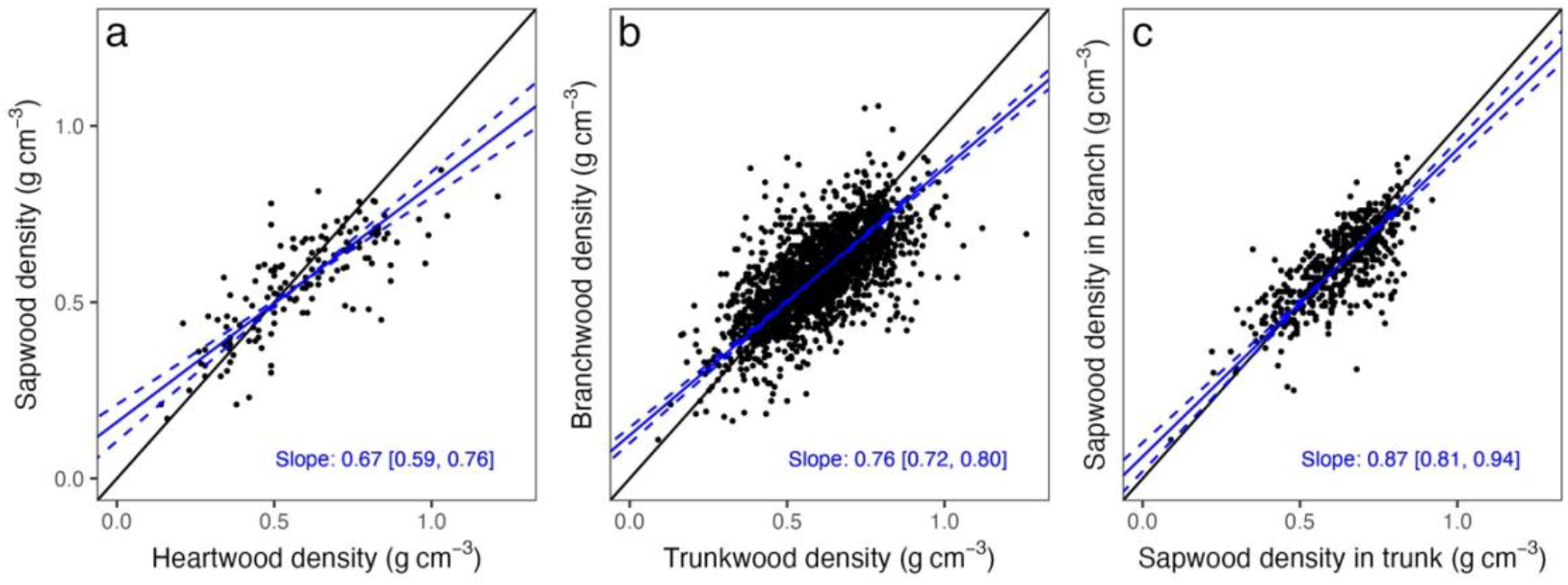
Variation in wood density across plant tissue types. Shown are plots of species means for a) heartwood vs. sapwood densities, b) trunkwood vs. branchwood densities and c) trunkwood vs. branchwood densities when measured on sapwood. Every dot represents a species, blue lines are Major Axis regression lines, *dashed* lines their 95% CI. Solid black lines are the 1:1 lines.

### Wood density variation among individuals

Among individuals within species, environmental factors had small, but consistent, effects on wood density variation (Fig. 3, Tables S10-13, Fig. S7-10). Wood density increased with temperature by 0.012 g cm^-3^ (standardized effect size) and with water deficit by 0.010 g cm^-3^; it decreased weakly with wind speed and soil pH (by -0.003 for both, Tables S10-11). These effects were strongly correlated with interspecific effects (r = 0.83, Fig. S12), but were smaller by 70-80% (Fig. 3, Figs. S7-10, Tables S10-13) and varied from one species to another. For example, a typical intraspecific increase in wood density by 0.010 g cm^-3^ amounted to ∼25% of the respective interspecific effect (0.038 g cm^-3^), and species varied widely around this mean (sd = 0.038 g cm^-3^, Table S10). Patterns were comparable when modelling tropical and extratropical species separately, but in the tropics, increases with water deficit (0.012 g cm^-3^ within and 0.059 g cm^-3^ among species) and decreases with pH (-0.014 within and -0.033 g cm^-3^ among species) were stronger. In a separate analysis for gymnosperms, there was a clear intraspecific decrease of -0.010 g cm^-3^ with soil pH and a strong intraspecific increase with water deficit (0.034 g cm^-3^, Table S15, Fig. S11). Since some species covered a narrow environmental gradient, effects were weaker when rescaling effect sizes to realized environmental ranges (Table S14, Figs. S13-14). For example, within the actual environmental limits of species observed in our data, tropical wood density increased by 0.012 g cm^-3^ with temperature instead of the 0.022 g cm^-3^ predicted by a more generic model.

**Fig. 3:**
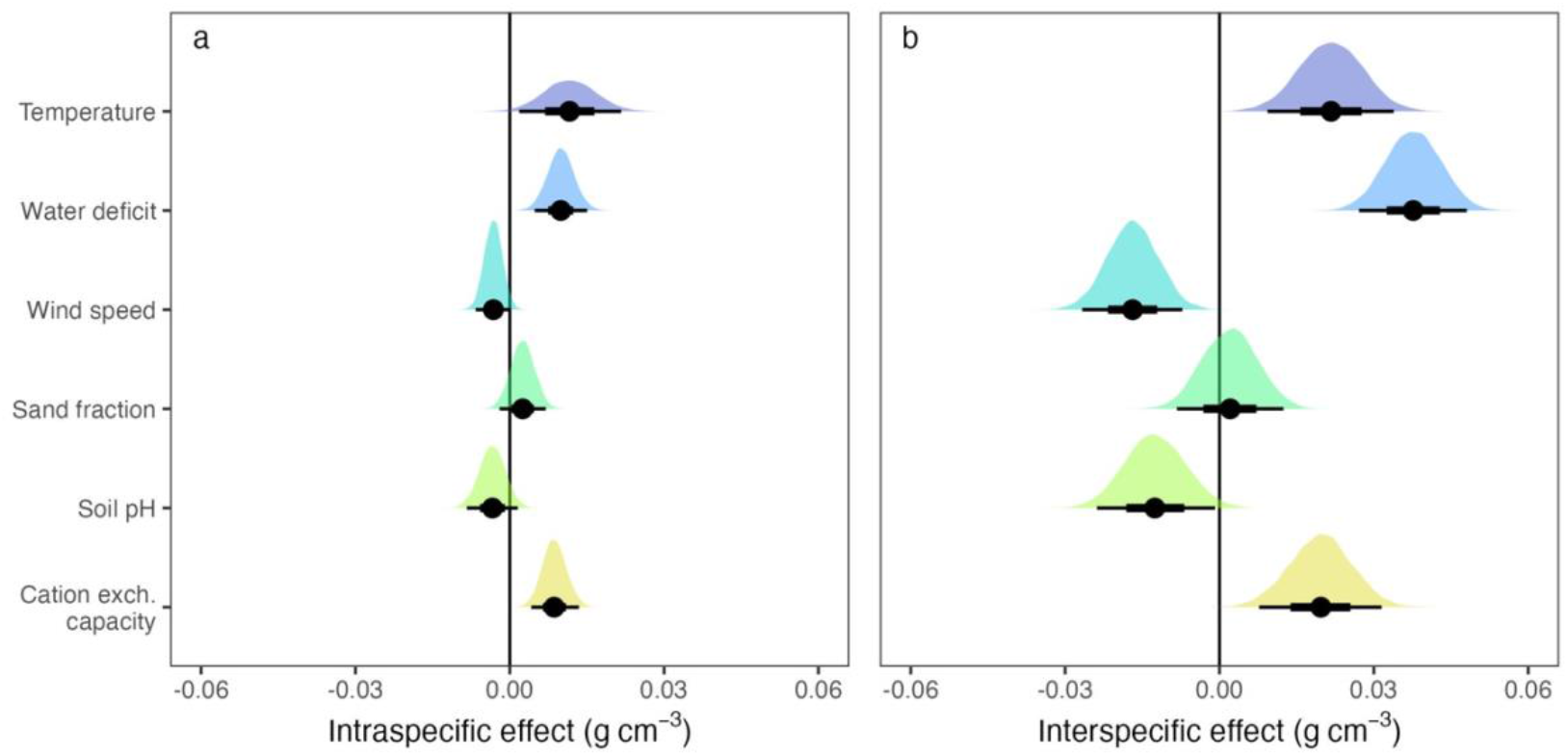
Environmental effects on wood density. Shown are the global effects of climatic and edaphic predictors on wood density, separated into intraspecific (a) and interspecific effects (b). Estimates of effect sizes were derived from a Bayesian hierarchical model, with all predictors scaled by one standard deviation (model M9, cf. Table S3 and Table S10). Climatic variables were from CHELSA/BIOCIM+ (Brun et al., 2022; Karger et al., 2017), edaphic variables from *soilgrids* (Hengl et al., 2017). Black dots indicate the median effect size and black intervals indicate quantile ranges (66% and 95%, partially covered by the black dots). The effects of cation exchange capacity were highly dependent on grid cell resolution and models, meaning effect sizes should be interpreted with caution (Fig. S7).

### The effect of intraspecific variation on wood density estimation

Intraspecific variation strongly reduced the accuracy of wood density estimates in undersampled species. A single wood density measurement was a poor approximation of the species mean (RMSE = 0.084 g cm^-3^); it was in fact comparable to a genus mean that did not involve any sampling of the target species (RMSE = 0.083 g cm^-3^, Table S16). However, the accuracy of species-level wood density estimates improved quickly with better sampling. From the average of two measurements, species-level wood density could already be predicted with RMSE = 0.056 g cm^-3^ (Table S16). Accuracy improved further when applying hierarchical models that combined individual measurements with taxonomic information from the remainder of the GWDD v.2 (RMSE = 0.038 g cm^-3^ for two measurements, Tables S16-17).

The wood density of individual plants was much harder to predict from conspecifics, with an RMSE of 0.107 g cm^-3^ and R^2^ = 0.59 when estimated from a single local wood density measurement (Fig. 4a, Tables S16-17). Errors were lower when pooling information with the GWDD v.2 via hierarchical modelling, but the improvements were moderate (RMSE = 0.086 g cm^-3^ or ca. 20% of the raw estimate, and R^2^ = 0.71, Fig. 4b). Errors were similar when three measures from local conspecifics were included, both as simple average (RMSE = 0.082 g cm^-3^) or based on a hierarchical model (RMSE = 0.083 g cm^-3^, Table S17).

**Fig. 4:**
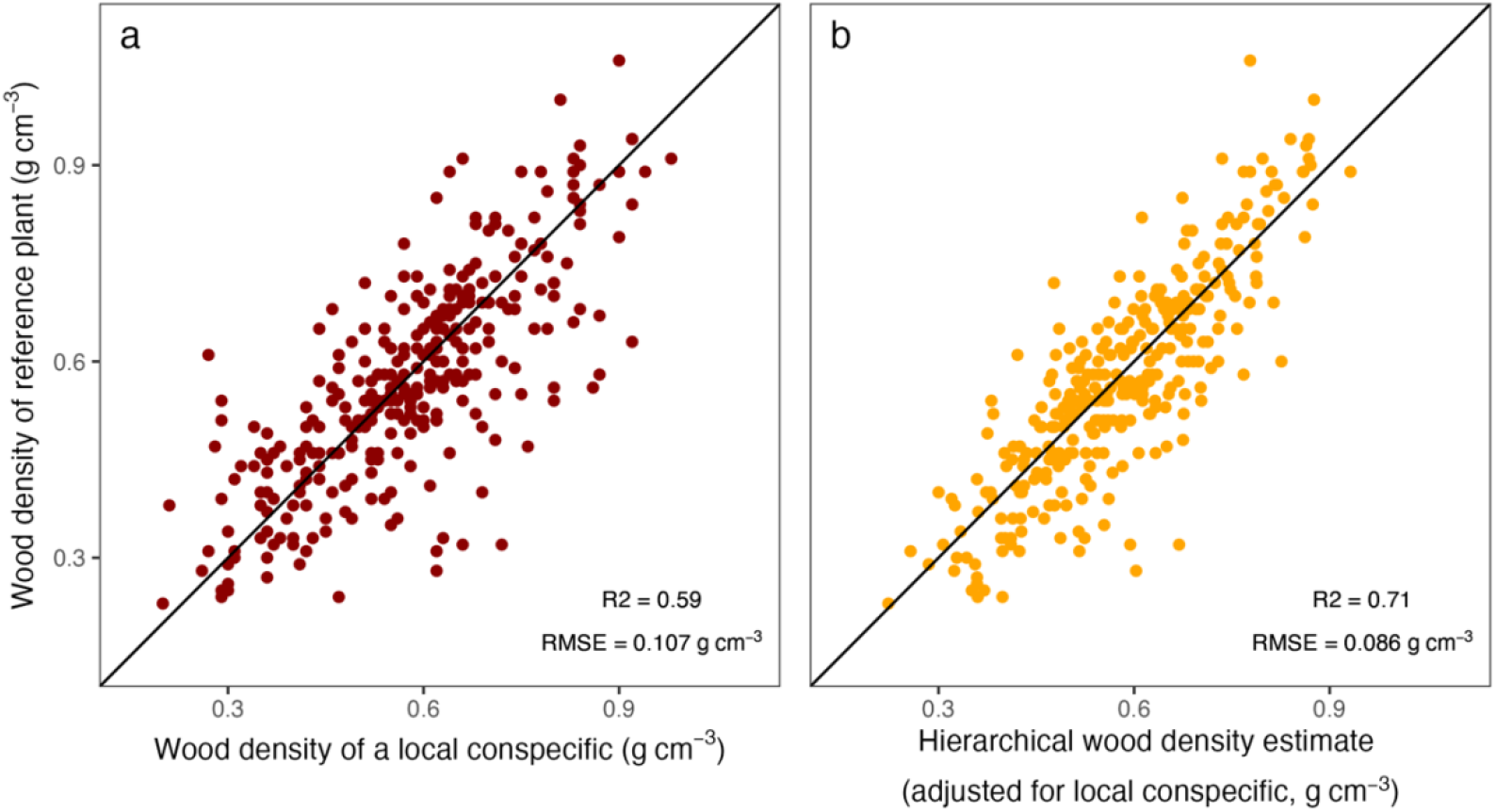
Prediction of an individual plant’s wood density from a local conspecific. Shown are two approaches to estimate an individual plant’s wood density when a measurement from a local conspecific is available: a) using the conspecific’s raw wood density measurement as an estimate, and b) using species or genus means adjusted by the local conspecific’s value via hierarchical models (model M27, Table S3).

## 4. Discussion

Wood density is a key trait in plant ecology, acting as an indicator of species’ competitive abilities (Kunstler et al., 2016), demographic rates (Adler et al., 2014), and ecological strategies (Chave et al., 2009; King et al., 2006; Kraft et al., 2010). The importance of species-level differences in wood density is well-known (Phillips et al., 2019). However, intraspecific variation is commonly considered less relevant due to the large amounts of wood density variation explained by taxonomic and phylogenetic relationships (Chave et al., 2009), the strong genetic control over wood density (Cornelius, 1994; Zobel & Jett, 1995) and a limited contribution to community-level variation in comparison with other plant traits (Siefert et al., 2015). Given that intraspecific variation in traits plays a crucial role for ecological processes (Bolnick et al., 2011; Des Roches et al., 2018) and may influence biomass estimates (Momo et al., 2020), we here re-examined this hypothesis at a global scale via the newly assembled Global Wood Density Database v.2, which, compared to the previous version (Zanne et al., 2009), included more than 6x as many records, more than doubled the number of species, and added key information on sources of intraspecific variation.

### Intraspecific variation in wood density matters

In the GWDD v.2, intraspecific variation in wood density (sd = 0.068 g cm^-3^) was substantial and structured according to both internal factors (within-plant structure) and external factors (environment). Intraspecific variation accounted for up to 15% of global wood density variation and followed predictable patterns with environmental factors. Wood density increased with water deficit and more weakly decreased with soil pH. These results were consistent across models and datasets (Table S10-13) and confirmed expectations that plants with high wood densities should be favoured in extreme and nutrient-poor environments (Anderegg et al., 2021; Gourlet-Fleury et al., 2011; Ibanez et al., 2017; Muller-Landau, 2004; Yang et al., 2023). There was also a general trend of increases in wood density with temperature, both in and out of the tropics, which may also reflect higher risks of drought-induced embolisms in hotter environments, but it was not as clear as trends with water availability (Tables S10-15). Crucially, environmental effects at intraspecific level were strongly correlated with interspecific effects (r = 0.83) and were also consistent with a companion study that found that community-means of wood density increased with temperature and aridity (Fischer et al., submitted). This consistency of environmental effects across scales was surprising, since previous studies found that variation in plant traits is often shaped by scale-dependent physiological and ecological mechanisms (Anderegg et al., 2018; Fajardo et al., 2024; Wang et al., 2022; Zhou et al., 2022). Wood density, with its genetic limitations on variation, might thus be an exception, with similar physiological constraints operating across different levels of ecological organization.

We also found that previously hypothesized patterns of variation in wood density within individuals (MacFarlane, 2020; Woodcock & Shier, 2002) generalized to global scales. Across species and studies, wood density varied less in sapwood than in heartwood and less in branchwood than in trunkwood. Species with low-density heartwood had denser sapwood and species with low-density trunkwood had denser branchwood, and vice versa in both cases (Fig. 2a and b). An explanation for the heartwood-sapwood trends may lie in distinct ecological strategies with changes in ontogeny (Hietz et al., 2013; Wiemann & Williamson, 1988; Woodcock & Shier, 2002), i.e., that some species tend to grow fast early in their life, investing little in dense wood, but slow down in later life. The opposite strategy – investing first in dense wood and then accelerating diameter growth later in life – has also been suggested, but appears less common (Osazuwa-Peters et al., 2014). Trunk-branchwood patterns have been explained through stronger functional constraints on branches than on trunks (MacFarlane, 2020; Momo et al., 2020), but it has also been argued that they may be explained by sapwood fractions (Gartner, 1995), since corewood formation at the tip of the stem occurs at the same time as the formation of outer wood at its base (cf. summary in Wiemann & Williamson, 2013). Indeed, when we compared only sapwood samples, differences between branch and trunkwood largely disappeared (Fig. 2c). These results need to be qualified, however. While we found clear global patterns, our findings relied on coarse, binary distinctions between woody tissue types and thus glossed over methodological differences between studies, as well as finer biological details. Changes in wood properties from the pith to bark or from base of trunk to branches often follow non-linear patterns and vary strongly between individual plants (Osazuwa-Peters et al., 2014; Schuldt et al., 2013; Terrasse et al., 2021), both of which remain to be studied in future research.

Our study also demonstrated that intraspecific variation in wood density has implications for applications such as carbon stock assessments or functional ecology. It is common practice to use individual wood density measurements as estimates for species means (Réjou-Méchain et al., 2017), but we found that individuals were generally so variable that a species mean was less accurately estimated from a single species measurement than from a genus mean. When applying hierarchical models, we found strong improvements in predictive accuracy for species means (from RMSEs g cm^-3^ of 0.084 down to 0.038 g cm^-3^ with two samples). However, increased sampling and hierarchical models did not help with predicting an individual plant’s wood density. Even from three samples of local conspecifics, RMSEs did not decrease below 0.082 g cm^-3^, indicating that there was substantial biological variation among and within individuals that could not be explained by site-specific environmental factors or phylogenetic relatedness. We note, however, that our study could not examine the local growth conditions and competitive neighbourhood of individuals, which likely play an important role in explaining intraspecific variation in wood density (Kunstler et al., 2016).

### But intraspecific variation in wood density is also limited

Despite the importance of intraspecific variation in wood density, we still found that its extent was limited. First, it accounted for less of the total variance than taxonomic variation at species (17%), genus (30%) or family (30%) levels. Second, average environmental effects and gradients within individuals were generally small (usually ∼0.01 g cm^-3^ or less, exceptionally ∼0.02 g cm^-3^). They amounted to only 20-30% of species-level effects and were outweighed both by methodological uncertainties, as may arise from differences in drying temperatures or wood coring (estimated at sd = 0.049 g cm^-3^ when counting only between-study differences, model M1, Table S4), and large among-species variation in the direction and magnitude of intraspecific effects. For example, across species, we found an average decrease of -0.023 g cm^-3^ from trunk to branchwood, but species varied in this relationship with sd = 0.075 (model M10, Table S8). This result means that branchwood density was still higher than trunkwood density by more than 0.015 g cm^-3^ in ca. 30% of species. Similarly, despite an intraspecific increase of wood density with water deficit of 0.010 g cm^-3^, species varied in this effect with sd = 0.038 g cm^-3^ (model M16, Table S10), meaning that in ca. 30% of species wood density decreased with water deficit by a similarly-sized 0.010 g cm^-3^.

It is possible that these estimates of variation in effect sizes were slightly too large, since global collections such as the GWDD v.2 accumulate variation from many sources. Also, they do not systematically sample across species’ entire environmental ranges, meaning errors across small environmental gradients and local growth conditions (light, rocks, waterlogging) may introduce uncertainty. Some studies, for example, found that intraspecific variation was better predicted when large aridity or temperature gradients were systematically sampled (Anderegg et al., 2021). However, other studies with systematic sampling designs did not find consistent effects of the environment on intraspecific variation (Richardson et al., 2013; Rosas et al., 2019), and a replication of our analysis with a higher-quality subset of the GWDD v.2 did not change the results: for example, we still found the same small intraspecific increase with water deficit (0.008 g cm^-3^), the same large variation around this mean effect (sd = 0.039 g cm^-3^), and the same much larger effect of 0.045 g cm^-3^ among species (model M18, Table S10).

Overall, this large variability in effect size and direction indicated that intraspecific wood density variation, even when following broad patterns, was difficult to predict. In practice, the inclusion of the location of tissues (branch vs. trunk) into hierarchical models of wood density variation improved predictions at the species level from RMSE = 0.043 to 0.038 g cm^-3^ (assuming two wood density measurements, Table S16), but impacts on individual plant wood density estimates were minimal, with RMSE = 0.085 vs. 0.084 g cm^-3^ (also assuming two wood density measurements, Table S17). These findings are consistent with several previous studies which found clear patterns for community- and species-level wood densities (Chave et al., 2009; Kraft et al., 2010; Swenson & Enquist, 2007) where errors were smaller than total variation among taxa, but could not replicate these results at the intraspecific level, with patterns seemingly unpredictable (Fajardo, 2018; Richardson et al., 2013; Rosas et al., 2019; Umaña & Swenson, 2019).

### When to account for intraspecific variation in wood density: A tale of two scales

Overall, our study showed that the decision of accounting for intraspecific variation in wood density depends on the scale of the research question. Measuring each individual’s wood density and how it changes across its organs is paramount when studying plastic growth responses in individual plants. Intraspecific variation of up to 0.068 g cm^-3^ meant that two measurements of individuals from the same species could easily be separated by as much as ±0.19 g cm^-3^ (95% interval of the difference between two draws from a normal distribution with sd = 0.068 g cm^-3^). This variability should be large enough to overwhelm most species differences at a single site and led to large errors when predicting an individual’s wood density, with R^2^ as low as 0.59 and RMSEs as high as 0.108 g cm^-3^ in this study (Fig. 4). Intraspecific variation in wood density may thus dominate species-level differences within communities or even across communities if these are dominated by only a few species that are close in average wood density values (cf. patterns in Anderegg et al., 2021). Another case where intraspecific variation should be accounted for is the determination of individual tree biomass from terrestrial laser scanning (TLS). TLS-derived 3D volumetric models can give precise volume estimates (Calders et al., 2015), which then must be combined with wood density to transfer volume to biomass. However, TLS estimates can be costly to construct, and their volume estimate precision only matters if wood density variation among individuals and along the hydraulic pathway are accounted for (Demol et al., 2021; Momo et al., 2020). Since we found errors of at least 0.08 g cm^-3^ when predicting wood density at the individual tree level (or between 10-20% of wood density, assuming most plants have densities between 0.40 and 0.80 g cm^-3^), studies likely cannot infer this variation, but need to systematically sample multiple individuals with at least two samples per measurement location.

By contrast, if the aim is to assess average community structure and vegetation dynamics at large scales or across steep environmental gradients, it makes sense to prioritize taxonomic coverage (Phillips et al., 2019) over the exhaustive sampling of individuals from a single species. First, as we showed here, variation at species or higher taxonomic levles accounted for most variation in wood density (77%). Second, environmental effects at the intraspecific level aligned with interspecific effects (r = 0.83). Third, variation among and within individuals within sites is expected to average out at the community level. Therefore, as long as a wide range of communities is sampled, the omission of intraspecific effects should not introduce systematic bias. In some cases, such as wood density predictions via machine learning models (Yang et al., 2024), it may even make sense to ignore intraspecific information on purpose, as the risk of overfitting or mistaking small methodological differences for biological variation outweighs the benefits of small corrections of ∼0.01 g cm^-3^ or less. Similarly, intraspecific variation in wood density should play a minor role when applying pre-calibrated allometric models to estimate tree biomass, as its variance is dwarfed by other sources of uncertainty, such as allometric models and estimates of plant size and shape (Chave et al., 2014; Kindermann et al., 2022; Molto et al., 2013; Réjou-Méchain et al., 2017).

### Global traits databases as backbones for hierarchical models

A key takeaway of our study is that, no matter the level of analysis, wood density measurements should not be treated as monolithic true values. Rather, they are noisy trait estimates that can be refined by including prior information through shared evolutionary history or measurement locations (Funk et al., 2017). Here, we applied simple hierarchical models based on taxonomic relationships and found that they vastly outperformed simple averaging procedures, particularly for undersampled species. A single wood density measurement was as poor an approximation of the species mean as a genus mean that did not involve any sampling of the target species (Tables S16), but errors decreased substantially when combining both in a hierarchical model (Fig. 4, Tables S16-17). Approaches could be further refined by explicitly accounting for phylogenetic relationships, but this approach comes with its own challenges (Revell, 2010), and taxonomic hierarchies may provide a good approximation (cf. https://statmodeling.stat.columbia.edu/2016/02/14/hierarchical-models-for-phylogeny-heres-what-everyones-talking-about/, last accessed on 9 September, 2024).

Overall, our findings suggest that there is great value in open-access trait database such as the GWDD v.2, as they synthesize knowledge across a range of disciplines and help correct noisy local estimates. They also provide insights on ecological strategies, variation across biogeographic realms, and, with careful curation and documentation (Augustine et al., 2024), allow us to explore intraspecific variation. At the time of writing, version 1 of the GWDD has been downloaded almost 20,000 times. Many of the applications of this database have been in assessing forest carbon storage, in connection with REDD+ projects or carbon credit accounting programs. As this sector is coming under closer scrutiny, reducing uncertainty in carbon estimates is a timely ambition, and the GWDD v.2 will be an important contribution. The findings in this study will also be helpful for developing theories about the evolution of carbon investments in plants (Castorena et al., 2022), improving the parameterization of global dynamic vegetation models, and providing guidance on how to account for intraspecific variation in ecological studies. In the future, we hope that the openly available and thoroughly documented GWDD v.2 will encourage the documentation and sharing of more wood density datasets and the construction of similar databases for other traits.

## Acknowledgments

F. J. Fischer acknowledges a European Research Council grant to Rupert Seidl under the European Union’s Horizon 2020 research and innovation program (Grant Agreement 101001905, FORWARD). J. Chave acknowledges an ‘Investissement d’Avenir’ grant managed by the Agence Nationale de la Recherche (CEBA grant: ANR-10-LABX-25-01 and TULIP grant: ANR-10-LABX-0041). T. Jucker acknowledges a UK NERC Independent Research Fellowship (grant: NE/S01537X/1), UKRI Frontier Research grant (grant: EP/Y003810/1) and a Research Project Grant from the Leverhulme Trust (grant: RPG-2020-341). A. Fajardo acknowledges Anid-Fondecyt 1231025. R.A.F. de Lima acknowledges grant 13/08722-5, São Paulo Research Foundation (FAPESP). G. Vieilledent acknowledges the French Foundation for Research on Biodiversity (BIOSCENEMADA project). L. F. Alves acknowledges a Smithsonian Tropical Research Institution, Center for Tropical Forest Science (CTFS) Research Grant (2009). J. Borah acknowledges the village councils of Kiphire (Fakim, Thanamir, and Tsundang village), Phek (Zhipu, Wazeho, and Washelo village), and Kohima (Dzuleke village) district, Nagaland, India. J. Camarero acknowledges the Spanish Ministry of Science and Innovation (projects PID2021-123675OB-C43 and TED2021-129770B-C21). J.A. Dar acknowledges the Anusandhan National Research Foundation (ANRF), Government of India, for funding (Ref. No.: SRG/2022/002286) and SRM University-AP for the Seed Grant (SRMAP/URG/E&PP/2022–23/012). A. K. Das acknowledges a research project grant from the Council of Scientific and Industrial Research, New Delhi (Government of India, project no.38(1349)/13/EMR-II dt.14.2.2013). Philip Fearnside acknowledges the National Council for Scientific and Technological Development (CNPq 12450/2021-4, 406941/2022-0), the Foundation for the Support of Research in the State of Amazonas (FAPEAM: 01.02.016301.02529/2024-87) and the Brazilian Research Network on Climate Change (RedeClima) (FINEP/Rede Clima 01.13.0353-00). S. Kothandaraman acknowledges the Science and Engineering Research Board, Department of Science and Technology (DST-SERB) (Ref. No.: PDF/2021/003742/LS). A. Linstädter, M. Dobler and L. Kindermann acknowledge funding by German Research Foundation (DFG) through funding codes CRC TRR-228/1 and TRR-228/2, and thank the University of Namibia (UNAM) as well as National Botanical Research Institute Windhoek for their support. A.L.A Lima acknowledges the National Council for Scientific and Technological Development (CNPq). Pernambuco Science and Technology Support Foundation (FACEPE). Cate Macinnis-Ng acknowledges a Rutherford Discovery Fellowship from the Royal Society Te Aparangi RDF-UOA1504; Marsden Fund Award from the Royal Society Te Apārangi UOA1207. L.F.S. Magnago acknowledges the Conselho Nacional de Desenvolvimento Científico e Tecnológico (CNPq), grant 307984/2022-2. A. Martin acknowledges the Natural Sciences and Engineering Research Council of Canada. A. Matheny acknowledges a NSF EAR Hydrological Sciences CAREER Award 2046768. E. Nogueira acknowledges the National Institute for Research in Amazonia (INPA). O. Razafindratsima acknowledges the Madagascar National Parks and the Malagasy Ministry of Environment for research permission, and numerous local field workers. S. Richardson acknowledges the Strategic Science Investment Fund administed by the NZ Ministry for Business, Innovation and Employment. Oris Rodriguez-Reyes acknowledges the Instituto de Ciencias Ambientales y Biodiversidad, Universidad de Panama; Sistema Nacional de Investigación Panama; STRI Panama. R. Salguero-Gomez acknowledges a NERC Independent Research Fellowship (NE/M018458/1) and a NERC Pushing the Frontiers grant (NE/X013766/1). L. F. Henao-Diaz and P. Stevenson acknowledge Ecopetrol No. 135-2009. M. van der Sanda acknowledges a Veni grant from the Dutch Research Council (NWO), NWO-VI.Veni.192.027. J.M.D. Torezan acknowledges a grant from the Brazilian science council (CNPq PELD 441510/2020-5). In addition, we thank Jordi Martínez-Vilalta and Teresa Rosas for their data contributions, supported by CGL2013-46808-R grant funded by MINECO, and Dmitry Schepaschenko for his contributions to the database and facilitating the coordination among contributors. Above all, we thank the work of thousands of anonymous trait collectors and wood technicians; taxonomists who maintain and harmonize species records; foresters, ecologists, students, and field assistants who measure woody plants on the ground; meteorologists who man weather stations and collect climate data; satellite engineers, climate scientists and IT developers who develop models and maps as well as the software to analyze them; as well as individual citizens, local communities, and public and private bodies that support this work.

## Author contributions

F.J. Fischer, J. Chave and A. Zanne conceived of the project. F.J. Fischer led the assembly and processing of the database, the analysis for the manuscript, and the writing of a first draft of the manuscript, with assistance from J. Chave, A. Zanne, T. Jucker, A. Fajardo, A. Fayolle, R.A.F. de Lima and G. Vieilledent. All co-authors contributed substantially through data collection, data assembly and revisions of the manuscript.

## Data availability

The Global Wood Density Database v.2 and derived wood density estimates at species and genus level will be made openly available in a dedicated Zenodo repository upon publication (doi: 10.5281/zenodo.16919509). All other data and code underpinning the results of this article are publicly archived in a separate Zenodo archive, which will also be publicly available upon publication (doi: 10.5281/zenodo.16928343).

