## Supplementary Materials for "Beyond species means – the intraspecific contribution to global wood density variation"

### A) Supplementary Methods

#### S1: Updating the first GWDD

*Correcting inconsistencies*

Entries in the first GWDD were updated to reflect structural changes in the GWDD v.2. In the case of large collections that had previously been transformed – through conversion factors or aggregation – or updated with new records since the publication of the first GWDD (Ilic et al., 2000; Langbour et al., 2019; Vieilledent et al., 2018), we reintegrated the entire data sets from scratch to ensure consistency. Where possible, this was done at the level of individual plants instead of species means. If published manuals were based on or overlapped with databases, but provided values for additional species from other sources (e.g., Détienne & Jacquet, 1983 and Langbour et al., 2019), we first included the raw values from the databases and then reincluded any non-matching taxa from the published literature to ensure continuity. All values were transformed with the new correction factors. Where re-integration would have been highly time-consuming or near-impossible (values from hard-to-access print-only manuals such as Desch, 1941, or previous compilations, such as Reyes et al., 1992), we inferred the source value by back-transforming entries with the original wood density conversion factor and reapplying a corrected factor. In this case, the taxon identifier from the original database was kept as *species_reference* and the column *backtransformed* is ticked. In a few cases, sources in the first GWDD could not be accessed anymore (Database of Brazilian Woods 2006, formerly: http://www.ibama.gov.br/lpf/madeira/default.htm) or have been overwritten in the meantime (ICRAF database, http://db.worldagroforestry.org/wd). Records exclusively attributed to these sources were removed (50 taxa in total, 1 lost genus, Table S2).

*Database extension*

In addition to updating the previous database, we searched the literature for new or previously overlooked studies and extracted values either manually or from supplementary files. To cover a wide range of trait values, we also wrote to authors of papers citing the original GWDD paper (Chave et al., 2009) and asked them whether they would be willing to participate in our effort, yielding more than 70 contributors (co-authors on this paper). While not a primary aim, we also received data for a range of wood density types outside of the scope of the first GWDD. For some tissue types (e.g., “bark” in *type_tissue*) and within-plant locations (“root” in *location_sample*), we included these values, as they are directly linked to intraspecific variation and form already part of some wood density assessments (e.g., bark is not always removed before estimating wood density). However, we decided not to include quantities such as green wood density (Niklas & Spatz, 2010) or dry mass fraction (Goodman et al., 2013), as there is no clear conversion to wood specific gravity.

**S2: Converting air- and ovendry density to basic wood density**

For air- and ovendry wood densities robust conversion factors to basic density can be derived (Vieilledent et al., 2018). The theory behind the conversion factors is as follows: wood is composed of a variable fraction of water, some of which is free to move in conduits (tracheids and vessels), and the rest is associated with other wood tissues including parenchyma and fibers. During drying, free water is gradually lost, until reaching the so-called fiber saturation point. The moisture at fiber saturation is quite variable across species from ca 10% to 50%, with a typical value of ~30% (Berry & Roderick, 2005; Vieilledent et al., 2018). Beyond this point, any further drying also shrinks the volume. If $V_s$ is the volume of the sample at fiber saturation moisture *S*, and $V_0$ is the volume when the sample has lost all of its water, then the volumetric shrinkage, or retractability, is the percent loss in volume $R = (V_s - V_0)/V_s \times 100$, which varies from 5% to 25% across species (Vieilledent et al., 2018).

From these values, it is possible to derive a conversion formula between wood density at any moisture content *w* ($D_w$) and basic density ($D_b$):

$$D_b = \frac{1 - (R/100) \times (S - w)}{1 + w/100} \times D_w$$

This formula has been used together with the CIRAD wood technology database to derive robust conversion factors (Vieilledent et al., 2018). The CIRAD wood technology database is a collection of *S*, *R*, and *D* values for 3,832 individual trees with >10 samples per individual and measurements at four moisture contents *w* (from 18% to 0%). The data set allows the estimation of $D_w$ at any *w* as well as an estimation of conversion factors to $D_b$ by fitting regression models with intercepts forced through the origin. For the GWDD v.2, we calculated conversion factors for wood densities at four common moisture levels: 0.819 for airdry densities at w = 15%, 0.828 for airdry densities at w = 12%, 0.840 for airdry densities at w = 8%, and 0.868 for ovendry densities (or w = 0%).

**S3: Details on Bayesian modelling of wood density variation**

We fitted linear mixed-effects either with a Bayesian framework in the *brms* R package (Bürkner, 2018) and the maximum-likelihood framework of *lme4* (Bates et al., 2015). We assume basic familiarity with mixed-effect model fitting in R (formulae provided in Table S3), and will only focus on details particular to the Bayesian context.

1/ <u>Distributional model:</u> Unlike *lme4*, *brms* allows to explicitly model the distributional parameter σ (i.e., the width of the residual distribution). We here use this to allow σ to vary across the same taxonomic groupings as the response variable (wood density). Given that variances are constrained to be positive and typically follow a lognormal distribution, we used the *brms* default of modelling σ on log-scales. In R's common random effects model notation, this can be simply expressed as *log(σ) ~ 1 + (1 | species)* or *log(σ) ~ 1 + (1 | family / genus / species)*, but we note that in *brms*, the log-transformation is carried out automatically, so we redefine the distributional parameter as $\sigma^* = exp(\sigma)$ and write: $\sigma^*$ *~ 1 + (1 | species)* or $\sigma^*$ *~ 1 + (1 | family / genus / species)*. A visual check of the assumption of lognormality is provided in Fig. S4.

2/ <u>Prior specifications:</u> Bayesian modelling requires the specification of priors, i.e. the provision of initial distributions for the parameters to be modelled. Since the data sets used in this study are large, exact prior choices have little influence on the resulting inference. Nevertheless, specification of weakly informative priors is recommended to constrain the initial parameter space and provide weak constraints on expected effect sizes (a form of regularization). Throughout all wood density models, we chose the following priors:

$$Intercept \sim \mathrm{N}(0.5, 0.25); lb = 0$$

$$Population\ level\ (fixed)\ effects \sim \mathrm{N}(0.0, 0.1)$$

$$Group\ level\ (random)\ effects \sim \mathrm{N}(0.1, 0.1); lb = 0$$

Here, *lb* stands for "lower bound". This means that we broadly expect the intercept for any wood density model to lie between 0 and 1.0, with fixed effects broadly constrained between -0.2 and 0.2, and random effects broadly constrained between 0 and 0.3 (all 95% intervals). Given that most wood density values lie between 0.3 and 0.8 and that we always standardize predictors (scaling and centering), effect sizes outside this range would be very large. Compare also to model results, where fixed effect sizes usually lie within a much more restricted [-0.05, 0.05] and never exceed [-0.1, 0.1]. In addition, we specify the following priors on the distributional parameter $\sigma = log(\sigma^*)$:

$$Intercept \sim \mathrm{N}(-3.0, 0.5)$$

$$Group\ level\ (random)\ effects \sim \mathrm{N}(0.5, 0.5); lb = 0$$

This corresponds to a realized $\sigma^*$ = exp(-3.0) ~ 0.5, with an approximate range of [0.02, 0.14].

3/ Model fitting and checks: All *brms* models were fit with *adapt_delta* == 0.95 and *max_treedepth* == 10. We always ran 4 chains in parallel, with 2000 iterations (including 1000 for warmup) and checked against warnings about divergent transitions, effective sampling sizes (ESS), and the mixing of chains (R-hat <= 1.02). If ESS were low, we increased the total number of iterations up to 5000. Model fits were visually checked via the inbuilt pp_check() function (representative examples in Fig. S3).

**S4: Overview over R packages**

For completeness, we here list all R packages used in the study. They include *terra* (Hijmans, 2023), *data.table* (Dowle & Srinivasan, 2023), *lme4* (Bates et al., 2015), *brms* (Bürkner, 2018), *WorldFlora* (Kindt, 2020), *ggplot2* (Wickham, 2016), *ggdist* (Kay, 2024), *patchwork* (Pedersen, 2024), *viridis* (Garnier et al., 2023), *sjPlot* (Lüdecke, 2021), *lmodel2* (Legendre, 2018), *rnaturalearth* (Massicotte & South, 2023), *rnaturalearthdata* (South, 2017), *corrplot* (Wei & Simko, 2021).

#### S5: Literature references for wood density values

Below we provide all the sources of data for the GWDD v.2, and the number of records from these sources. Note that these records also include bark density measurements.

| Source | N |
|---|---|
| A.T.I.B.S. 1975. Nomenclature Generale des Bois Tropicaux. Nogent-sur-Marne, France. | 186 |
| Acevedo Mallque, M. and Kikata, Y. 1994. Atlas of Peruvian Woods. Universidad Nacional Agraria, La Molina, Peru and Nagoya University, Japan. 202 pp. | 11 |
| Ackerly, D.D. 2004. Functional strategies of chaparral shrubs in relation to seasonal water deficit and disturbance. Ecological Monographs 74:25-44. | 20 |
| Aguilar-Rodríguez et al. 2001: Comparación de la gravedad específica y características anatómicas de la madera de dos comunidades vegetales en México. Anales del Instituto de Biología. Universidad Nacional Autónoma de México. Serie Botánica 72, 171-185. | 54 |
| Aiba, M., & Nakashizuka, T. (2009). Architectural differences associated with adult stature and wood density in 30 temperate tree species. Functional Ecology, 23(2), 265-273. | 30 |
| Aiba, M., Kurokawa, H., Onoda, Y., Oguro, M., Nakashizuka, T., & Masaki, T. (2016). Context-dependent changes in the functional composition of tree communities along successional gradients after land-use change. Journal of Ecology, 104(5), 1347-1356. + unpublished data | 200 |
| Alden, H. 1995. Hardwoods of North America. United States Department of Agriculture, Forest Service, Forest Products Laboratory. Gen. Tech. Report FPL-GTR-83. 136 pp. http://www2.fpl.fs.fed.us/TechSheets/hardwood.html. From Wiemann database | 96 |
| Alden, H. 1997. Softwoods of North America. United States Department of Agriculture, Forest Service, Forest Products Laboratory. Gen. Tech. Report FPL-GTR-102. 151 pp. http://www2.fpl.fs.fed.us/TechSheets/softwood.html. From Wiemann database | 51 |
| Alfaro-Sánchez et al. 2020. How do social status and tree architecture influence radial growth, wood density and drought response in spontaneously established oak forests? Annals of Forest Science 77, 49. | 591 |
| Alston, A.S. 1982. Timbers of Fiji: Properties and Potential Uses. Department of Forestry, Suva, Fiji. 183 pp. | 40 |
| Altamirano, V., and Rico, R.L. 1992. Maderas de Bolivia: Caracteristicas y Usos de 55 Maderas Tropicales. Camara Nacional Forestal, Santa Cruz, Bolivia. | 8 |
| Alvarez et al. 2012. Tree above-ground biomass allometries for carbon stocks estimation in the natural forests of Colombia. Forest Ecology and Management 267, 297–308. | 257 |
| Alvarez et al. 2013. Densidad básica del fuste de árboles del bosque seco en la costa Caribe de Colombia. Rev. Intropica 8, 17-28. | 124 |
| Alves, LF and Oliveira, AA (2010) Assessment of aboveground forest carbon pools of a tropical moist forest (Ilha do Cardoso, Brazil). Smithsonian Tropical Research Institute, Center for Tropical Forest Science. Final Report | 20 |
| Amaro, M.A. 2010. Quantificação do estoque de volume, biomassa e carbono em uma Floresta Estacional Semidecidual Montana em Viçosa, MG. Tese (doutorado). Universidade Federal de Viçosa, Viçosa. 168p.; Amaro, M.A., Soares, C.P.B., Souza, A.L.D., Leite, H.G., & Silva, G.F.D. 2013. Volume, biomass and carbon stocks in a seasonal semideciduous forest in viçosa, minas gerais state. Revista Árvore, 37(5), 849-857. | 28 |
| Amorim, L.C. 1991. Variação da densidade basica no sentido radial em madeiras tropicais da Amazônia. Relatorio final Periodo abril 90: março 91. INPA, Manaus, AM 24 pp. In Fearnside, P. M. 1997. Wood density for estimating forest biomass in Brazilian Amazonia. Forest Ecology and Management 90: 59-87. | 1 |
| Anderegg et al. 2021. Aridity drives coordinated trait shifts but not decreased trait variance across the geographic range of eight Australian trees. New Phytologist 229(3), 1375-1387. doi:10.1111/nph.16795 | 1603 |
| Anonymous 1981. Mengenal Sifat-sifat Kayu Indonesia dan Penggunaannya. Penerbit Kanisius. ISBN 979-413-106-7. | 115 |
| Anonymous. 1971. Inventaire Forestier des Terres Basses du Versant Occidental des Monts Cardamones CAMBOOGE. Confidential FAO Technical Report No. 6, FO:SF/CAM 6., Rome, Italy. Reported as air-dry density in Miller-Ilic database | 128 |
| Anonymous. 1974. Standard Nomenclature of Forest Plants, Burma, including commercial timbers. Forest Research and Training Circle, Forest Department, Burma. 121 pp. . Reported as air-dry density in Miller-Ilic database | 233 |
| Anonymous. 1979. La Amazonia Colombiana y sus recursos. Proyecto Radargrametrico de Amazonas. IGAC, Bogota. By H ter Steege. | 68 |
| Anonymous. 1990. Nomenclature of Commercial timbers, including sources of supply. (Revision of BS 881 and 589: 1974). British Standards Association, London, England. | 4 |
| Anufrieva 1976, Schepaschenko et al. (2017) Sci. Data 4:170070. DOI: 10.1038/sdata.2017.70 | 3 |
| Apgaua DMG, Ishida FY, Tng DYP, Laidlaw MJ, Santos RM, Rumman R, Eamus D, Holtum JAM, Laurance SGW (2015) Functional traits and water transport strategies in lowland tropical rainforest trees. PLoS ONE 10, e0130799. | 8 |
| Apgaua DMG, Tng DYP, Cernusak LA, Cheesman AW, Santos RM, Edwards WJ, Laurance SGW (2017) Plant functional groups within a tropical forest exhibit different wood functional anatomy. Functional Ecology 31, 582-591. | 82 |
| Arcanjo, Fátima; Bordignon, Alexandre; Torezan, José Marcelo; Miranda, Carmem (2016). Field data. Source: ARCANJO, Fátima Aparecida. Biomass in semideciduous Atlantic Forest fragments and restoration sites. 2017.89. Dissertation (Master in Biological Sciences) – State University of Londrina, 2017. | 75 |
| Arostegui V., A. 1976. Características Tecnológicas y Usos de la Madera de 145 Especies del País. 483 pp. | 12 |
| Arostegui, A. and Sobral Filho, M. 1986. Usos de las maderas del bosque humedo tropical Colonia Angamos Rio Yavari y Jenaro Herrera, Investigaciones Tecnologicas 1:2, Instituto de Investigaciones de la Amazonia Peruana, Iquitos By T Baker. | 4 |
| Arostegui, A. and Valderamma, F. 1986. Usos de las maderas del bosque humedo tropical Allpahuayo-Iquitos. Investigaciones Tecnologicas 1:5, Instituto de Investigaciones de la Amazonia Peruana, Iquitos By T Baker. | 11 |
| Askarov 1974; Malenko et al. 2015, Schepaschenko et al. (2017) Sci. Data 4:170070. DOI: 10.1038/sdata.2017.70 | 37 |
| Asner et al. 2011. High-resolution carbon mapping on the million-hectare Island of Hawaii. Frontiers in Ecology and the Environment 9(8), 434–439. doi:10.1890/100179. Obtained from Appendix in Flint et al. 2014. | 30 |
| Asrat et al. 2020. Modelling and quantifying tree biometric properties of dry Afromontane forests of south-central Ethiopia. Trees 34, 1411–1426. | 60 |

| | |
|---|---|
| Auburn_Silviculture, from: Radtke et al. 2016. LegacyTreeData: A national database of detailed tree measurements for volume, weight, and physical properties. In: Stanton SM, Christensen GA, (eds). Proceedings of Forest Inventory and Analysis 2015 Science Symposium, December 8–10, 2015, Portland, OR. GTR-PNW-931, USDA Forest Service, Pacific Northwest Research Station, GTR-PNW-931, Portland, OR, pp. 25–30. | 140 |
| Auclair Metayer 1980, Schepaschenko et al. (2017) Sci. Data 4:170070. DOI: 10.1038/sdata.2017.70 | 1 |
| B Davis 1994. Wood densities for fifty-two Australian tree species. CSIROIn: Ilic, J., Boland, D., McDonald, M., Downes, G. and Blakemore, P. 2000. Woody density phase 1. State of knowledge. National carbon accounting system. Technical Report 18 Australian Greenhouse Office, Canberra, Australia. | 61 |
| Baldwin 1987, from: Radtke et al. 2016. LegacyTreeData: A national database of detailed tree measurements for volume, weight, and physical properties. In: Stanton SM, Christensen GA, (eds). Proceedings of Forest Inventory and Analysis 2015 Science Symposium, December 8–10, 2015, Portland, OR. GTR-PNW-931, USDA Forest Service, Pacific Northwest Research Station, GTR-PNW-931, Portland, OR, pp. 25–30. | 260 |
| Baldwin_and_Saucier, from: Radtke et al. 2016. LegacyTreeData: A national database of detailed tree measurements for volume, weight, and physical properties. In: Stanton SM, Christensen GA, (eds). Proceedings of Forest Inventory and Analysis 2015 Science Symposium, December 8–10, 2015, Portland, OR. GTR-PNW-931, USDA Forest Service, Pacific Northwest Research Station, GTR-PNW-931, Portland, OR, pp. 25–30. | 452 |
| Baldwin_et_al_2000, from: Radtke et al. 2016. LegacyTreeData: A national database of detailed tree measurements for volume, weight, and physical properties. In: Stanton SM, Christensen GA, (eds). Proceedings of Forest Inventory and Analysis 2015 Science Symposium, December 8–10, 2015, Portland, OR. GTR-PNW-931, USDA Forest Service, Pacific Northwest Research Station, GTR-PNW-931, Portland, OR, pp. 25–30. | 214 |
| Baldwin_et_al, from: Radtke et al. 2016. LegacyTreeData: A national database of detailed tree measurements for volume, weight, and physical properties. In: Stanton SM, Christensen GA, (eds). Proceedings of Forest Inventory and Analysis 2015 Science Symposium, December 8–10, 2015, Portland, OR. GTR-PNW-931, USDA Forest Service, Pacific Northwest Research Station, GTR-PNW-931, Portland, OR, pp. 25–30. | 34 |
| Barajas-Morales, J. 1987, Wood specific gravity in species from two tropical forests in México. International Association of Wood Anatomists Bulletin, 8, 143-J148. | 212 |
| Barbosa, R.I. and Ferreirea, C.A.C. 2004. Densidade basica da madeira de um ecossistema de 'campina' em Roraima, Amazonia Brasileira. Acta Amazonica 34: 587-591. | 30 |
| Barnard D.M., Meinzer F.C., Lachenbruch B., McCulloh K.A., Johnson D.M., Woodruff D.R. 2011. Climate-related trends in sapwood biophysical properties in two conifers: avoidance of hydraulic dysfunction through coordinated adjustments in xylem efficiency, safety and capacitance. Plant, Cell and Environment 34:643-54. | 4 |
| Bencat 1989, Schepaschenko et al. (2017) Sci. Data 4:170070. DOI: 10.1038/sdata.2017.70 | 18 |
| Bencat 1990, Schepaschenko et al. (2017) Sci. Data 4:170070. DOI: 10.1038/sdata.2017.70 | 1 |
| Bendtsen, B A. and Chudnoff, M. 1981. Properties of Seven Colombian Woods. Research Note FPL-0242. U.S. Department of Agriculture, Forest Service, Forest Products Laboratory. 12 pp. | 3 |
| Benthall, A.P. 1984. The Trees of Calcutta: And its Neighborhood. Thacker Spink and Co. Ltd. Calcutta India. | 136 |
| Berner, L. T., & Law, B. E. (2015). Water limitations on forest carbon cycling and conifer traits along a steep climatic gradient in the Cascade Mountains, Oregon. Biogeosciences, 12(22), 6617. | 3 |
| Bhaskar, R. A. Valiente-Banuet, D.D. Ackerly. 2007. Evolution of hydraulic traits in closely related species pairs from mediterranean and non-mediterranean environments of North America. New Phytologist 176: 718-26. | 12 |
| Bhatt et al. 2017. Fuelwood characteristics of important trees and shrubs of Eastern Himalaya, Energy Sources, Part A: Recovery, Utilization, and Environmental Effects 39(1), 47-50. | 24 |
| Blakemore P. 2003. Density and Shrinkage of four low rainfall plantation grown eucalypts. CSIROIn: Ilic, J., Boland, D., McDonald, M., Downes, G. and Blakemore, P. 2000. Woody density phase 1. State of knowledge. National carbon accounting system. Technical Report 18 Australian Greenhouse Office, Canberra, Australia. | 4 |
| Boanerges Souza 2014. EFEITOS DO SOLO E NÍVEL DO LENÇOL FREÁTICO SOBRE A VARIAÇÃO DA GRAVIDADE ESPECÍFICA DA MADEIRA EM MESOESCALA NO NORTE DA AMAZÔNIA. Dissertation, INPA, Manaus, Amazonas. | 169 |
| Bolza E and Kloot N. H. 1963. The mechanical properties of 174 Australian Timbers. CSIROIn: Ilic, J., Boland, D., McDonald, M., Downes, G. and Blakemore, P. 2000. Woody density phase 1. State of knowledge. National carbon accounting system. Technical Report 18 Australian Greenhouse Office, Canberra, Australia. | 205 |
| Bolza, E. 1975. Properties and uses from 175 timber species from Papua New Guinea and West Irian. C.S.I.R.O. Division of Building Research Report 34. | 232 |
| Bondarenko 1970, Schepaschenko et al. (2017) Sci. Data 4:170070. DOI: 10.1038/sdata.2017.70 | 8 |
| Brink and Achigan-Dako 2012. Plant Resources of Tropical Africa 16. Fibres. Wageningen. | 1 |
| Brodribb T.J. & Cochard H. 2009. Hydraulic failure defines the recovery and point of death in water-stressed conifers. Plant physiology 149:575-84. | 6 |
| Brooks_et_al, from: Radtke et al. 2016. LegacyTreeData: A national database of detailed tree measurements for volume, weight, and physical properties. In: Stanton SM, Christensen GA, (eds). Proceedings of Forest Inventory and Analysis 2015 Science Symposium, December 8–10, 2015, Portland, OR. GTR-PNW-931, USDA Forest Service, Pacific Northwest Research Station, GTR-PNW-931, Portland, OR, pp. 25–30. | 84 |
| Broshtilova 1983, Schepaschenko et al. (2017) Sci. Data 4:170070. DOI: 10.1038/sdata.2017.70 | 6 |
| Brzeziecki, B. and Kienast, F. 1994. Classifying the life-history strategies of trees on the basis of the Grimian model. Forest Ecology and Management 69: 167-187. | 34 |
| Bucci S., Scholz F.G., Peschiutta M.L., Arias M.S., Meinzer F.C., Goldstein G. 2013. The stem xylem of Patagonian shrubs operates far from the point of catastrophic dysfunction and is additionally protected from drought-induced embolism by leaves and roots. Plant, Cell and Environment 36: 2163-2174. | 14 |
| Bucci S.J. Scholz F.G., Goldstein G and Meinzer F.C. 2009. Soil water availability as determinant of the hydraulic architecture in Patagonian woody species. Oecologia DOI 10.1007/s00442-009-1331-z. | 7 |
| Bucci S.J., Goldstein G., Meinzer F.C., Scholz F.G., Franco A.C. and Bustamante M. 2004.. Functional convergence in hydraulic architecture and water relations of savanna trees: from leaf to whole plant. Tree Physiology 24: 891-899. | 2 |
| Bucci, S.J., Scholz F.G., Goldstein G., Meinzer F.C., Franco, A.C., Campanello, P.I., Villalobos-Vega, R., Bustamante, M. and Miralles-Wilhelm, F. 2006. Nutrient availability constrains the hydraulic architecture and water relations of savanna trees. Plant Cell and Environment 29: 2153-2167 | 20 |
| Bucci, S.J., Scholz, F.G., Campanello, L.M., Montti, L., Jimenez-Castillo, M., Rockwell, F.A., La Manna, L., Guerra, P. Bernal, P.L., Troncoso, O. Enricci, J., Holbrook M.N. and Goldstein G. 2012. Hydraulic differences along the water transport system of South American Nothofagus species: do leaves protect the stem functionality? Tree Physiology 32:880-893, doi: 10.1093/treephys/tps054 | 9 |
| Buckeye_Cell_Slash, from: Radtke et al. 2016. LegacyTreeData: A national database of detailed tree measurements for volume, weight, and physical properties. In: Stanton SM, Christensen GA, (eds). Proceedings of Forest Inventory and Analysis 2015 Science Symposium, December 8–10, 2015, Portland, OR. GTR-PNW-931, USDA Forest Service, Pacific Northwest Research Station, GTR-PNW-931, Portland, OR, pp. 25–30. | 160 |
| Buckeye_Planted_Slash, from: Radtke et al. 2016. LegacyTreeData: A national database of detailed tree measurements for volume, weight, and physical properties. In: Stanton SM, Christensen GA, (eds). Proceedings of Forest Inventory and Analysis 2015 Science Symposium, December 8–10, 2015, Portland, OR. GTR-PNW-931, USDA Forest Service, Pacific Northwest Research Station, GTR-PNW-931, Portland, OR, pp. 25–30. | 168 |

| | |
|---|---|
| Budgen B. 1981. Shrinkage and density of Australian and other South-east Pacific woods. CSIROIn: Ilic, J., Boland, D., McDonald, M., Downes, G. and Blakemore, P. 2000. Woody density phase 1. State of knowledge. National carbon accounting system. Technical Report 18 Australian Greenhouse Office, Canberra, Australia. | 22 |
| Burger 1945, Schepaschenko et al. (2017) Sci. Data 4:170070. DOI: 10.1038/sdata.2017.70 | 2 |
| Burger 1947, Schepaschenko et al. (2017) Sci. Data 4:170070. DOI: 10.1038/sdata.2017.70 | 5 |
| Burger 1948, Schepaschenko et al. (2017) Sci. Data 4:170070. DOI: 10.1038/sdata.2017.70 | 4 |
| Burger 1953, Schepaschenko et al. (2017) Sci. Data 4:170070. DOI: 10.1038/sdata.2017.70 | 31 |
| Burkhart_and_Clutter 1971, from: Radtke et al. 2016. LegacyTreeData: A national database of detailed tree measurements for volume, weight, and physical properties. In: Stanton SM, Christensen GA, (eds). Proceedings of Forest Inventory and Analysis 2015 Science Symposium, December 8–10, 2015, Portland, OR. GTR-PNW-931, USDA Forest Service, Pacific Northwest Research Station, GTR-PNW-931, Portland, OR, pp. 25–30. | 702 |
| Burkhart_et_al, from: Radtke et al. 2016. LegacyTreeData: A national database of detailed tree measurements for volume, weight, and physical properties. In: Stanton SM, Christensen GA, (eds). Proceedings of Forest Inventory and Analysis 2015 Science Symposium, December 8–10, 2015, Portland, OR. GTR-PNW-931, USDA Forest Service, Pacific Northwest Research Station, GTR-PNW-931, Portland, OR, pp. 25–30. | 1088 |
| Buzykin et al. 2002, Schepaschenko et al. (2017) Sci. Data 4:170070. DOI: 10.1038/sdata.2017.70 | 14 |
| Buzykin Pshenichnikova 1980, Schepaschenko et al. (2017) Sci. Data 4:170070. DOI: 10.1038/sdata.2017.70 | 8 |
| Caetano Ferreira et al. 2012: Wood anatomy and technological properties of an endangered species: Picconia azorica (Oleaceae). IAWA Journal 33 (4), 375-390. | 1 |
| CAPPS, from: Radtke et al. 2016. LegacyTreeData: A national database of detailed tree measurements for volume, weight, and physical properties. In: Stanton SM, Christensen GA, (eds). Proceedings of Forest Inventory and Analysis 2015 Science Symposium, December 8–10, 2015, Portland, OR. GTR-PNW-931, USDA Forest Service, Pacific Northwest Research Station, GTR-PNW-931, Portland, OR, pp. 25–30. | 498 |
| Carlyn A. Raymond, and Andrew C. MacDonald 1998. Where to shoot your pilodyn: within tree variation in basic density in plantation Eucalyptus globulus and E. nitens in Tasmania. CSIROIn: Ilic, J., Boland, D., McDonald, M., Downes, G. and Blakemore, P. 2000. Woody density phase 1. State of knowledge. National carbon accounting system. Technical Report 18 Australian Greenhouse Office, Canberra, Australia. | 2 |
| Carrasco L.O., Bucci S.J., Francescantonio D.D., Lezcano O.A., Campanello P.I., Scholz F.G., Rodríguez S., Madanes N., Cristiano P.M., Hao G.-Y., Holbrook N.M., Goldstein G. 2014. Water storage dynamics in the main stem of subtropical tree species differing in wood density, growth rate and life history traits. Tree Physiology 35: 354-365. | 10 |
| Carrillo, A., Garza, M., de Jesris Nañez, M., Garza, F., Foroughbakhch, R., & Sandoval, S. (2011). Physical and mechanical wood properties of 14 timber species from Northeast Mexico. Annals of forest science, 68(4), 675-679. | 14 |
| Cartuche Peralta 2022: Caracterización de la madera de 95 especies forestales del sur de Ecuador con base a sus propriedades físicas, organolépticas y anatómicas. Tesis. Universidad Nacional de Loja. | 95 |
| Casas LF, Aldana AM, Henao-Días F, Stevenson PR (2017). Specific gravity of woody tissue from lowland Neotropical plants: Differences among forest types. Ecology 98(5) | 2602 |
| Cavender-Bares, J., Kitajima, K., and Bazzaz, F.A. 2004. Multiple trait associations in relation to habitat differentiation among 17 Floridian oak species. Ecological Monographs 74: 635-J661. | 17 |
| Cerny 1990, Schepaschenko et al. (2017) Sci. Data 4:170070. DOI: 10.1038/sdata.2017.70 | 21 |
| Chapotin S.M., Razanameharizaka J.H., Holbrook N.M. 2006. Water relations of baobab trees (Adansonia spp. L.) during the rainy season: does stem water buffer daily water deficits? Plant Cell and Environment 29: 1021-1032. | 2 |
| Chaturvedi and Raghubanshi 2015. Assessment of carbon density and accumulation in mono- and multi-specific stands in Teak and Sal forests of a tropical dry region in India. Forest Ecology and Management 339, 11-21. | 103 |
| Chaturvedi et al. 2011. Carbon density and accumulation in woody species of tropical dry forest in India. Forest Ecology and Management 262, 1576–1588. | 94 |
| Chen B. Chen C. 1980, Schepaschenko et al. (2017) Sci. Data 4:170070. DOI: 10.1038/sdata.2017.70 | 3 |
| Chen et al. 2016. Time lags between crown and basal sap flows in tropical lianas and co-occurring trees. Tree Physiology 36(6), 736–747. | 45 |
| Chen, J.-W., Zhang, Q., Li, X.-S. and K.-F. Cao. 2009. Independence of leaf and stem hydraulic straits in six Euphorbiaceae species with contrasting leaf phenology. Plant 230: 459-468. Chen J.W., Zhang Q., Cao K.F. 2009. Inter-species variation of photosynthetic and xylem hydraulic traits in the deciduous and evergreen Euphorbiaceae tree species from a seasonally tropical forest in south-western China. Ecological Research 24: 65-73. | 6 |
| Cheng, J. 1980. Chinese tropical and subtropical timbers: their distinction, properties and applications. China Forestry publishing house, Beijing. | 10 |
| Cheng, J. 1985. Wood Science. Chinese Forestry Publishing; Beijing, China. 1379 pp. | 60 |
| Cheng, J.C., Yang, J. and Liu, P. 1992. Anatomy and Properties of Chinese Woods. Chinese Forestry Publishing, Beijing, China. 820 pp. | 922 |
| Chichignoud, M., Deon, G., Detienne, P., Parant, B. and P. Vantomme. 1990. Atlas des Bois Tropicaux d'Amerique Latine. CIRAD-Foret, Nogent-Sur-Marne France, and Organisation internationale des Bois Tropicaux, Yokohama, Japan. | 3 |
| Chimelo et al. (1976) apud Paula, J.E. & Alves, J.L.H. 2010. 922 Madeiras nativas do Brasil: anatomia, dendrologia, dendrometria, produção e uso. Ed. Cinco Continentes, Porto Alegre. 461p. | 1 |
| Choat B, Ball M, Luly J, Holtum J. 2003. Pit membrane porosity and water stress-induced cavitation in four co-existing dry rainforest tree species. Plant Physiology 131: 41–48. | 4 |
| Chudnoff, M. 1984. Tropical Timbers of the World. Washington, DC, USDA Forest Service. http://www2.fpl.fs.fed.us/TechSheets/tropicalwood.html. | 148 |
| Chudnoff, M., 1973. Physical, Mechanical, and other properties of Selected Secondary Species in Surinam, Peru, Colombia, Nigeria, Gabon, Philippines, and Malaysia. FPL-AID-PASA TA(AJ)2-73 (Species Properties). 77 pp. | 20 |
| Cintra, B. B. L., Schietti, J., Emillio, T., Martins, D., Moulatlet, G., Souza, P., ... & Schöngart, J. (2013). Soil physical restrictions and hydrology regulate stand age and wood biomass turnover rates of Purus–Madeira interfluvial wetlands in Amazonia. Biogeosciences, 10(11), 7759-7774. | 17 |
| CIRAD database: Vieilledent et al. 2017, https://doi.org/10.5281/zenodo.1095453 | 4014 |
| Clark 1986, from: Radtke et al. 2016. LegacyTreeData: A national database of detailed tree measurements for volume, weight, and physical properties. In: Stanton SM, Christensen GA, (eds). Proceedings of Forest Inventory and Analysis 2015 Science Symposium, December 8–10, 2015, Portland, OR. GTR-PNW-931, USDA Forest Service, Pacific Northwest Research Station, GTR-PNW-931, Portland, OR, pp. 25–30. | 10658 |
| Clark_and_Saucier 1990, from: Radtke et al. 2016. LegacyTreeData: A national database of detailed tree measurements for volume, weight, and physical properties. In: Stanton SM, Christensen GA, (eds). Proceedings of Forest Inventory and Analysis 2015 Science Symposium, December 8–10, 2015, Portland, OR. GTR-PNW-931, USDA Forest Service, Pacific Northwest Research Station, GTR-PNW-931, Portland, OR, pp. 25–30. | 2264 |

| | |
|---|---|
| Clark_and_Taras 1976, from: Radtke et al. 2016. LegacyTreeData: A national database of detailed tree measurements for volume, weight, and physical properties. In: Stanton SM, Christensen GA, (eds). Proceedings of Forest Inventory and Analysis 2015 Science Symposium, December 8–10, 2015, Portland, OR. GTR-PNW-931, USDA Forest Service, Pacific Northwest Research Station, GTR-PNW-931, Portland, OR, pp. 25–30. | 440 |
| Clark_ChipnSaw_BlackOak 1970, from: Radtke et al. 2016. LegacyTreeData: A national database of detailed tree measurements for volume, weight, and physical properties. In: Stanton SM, Christensen GA, (eds). Proceedings of Forest Inventory and Analysis 2015 Science Symposium, December 8–10, 2015, Portland, OR. GTR-PNW-931, USDA Forest Service, Pacific Northwest Research Station, GTR-PNW-931, Portland, OR, pp. 25–30. | 81 |
| Clark_ChipnSaw_Loblolly 1970, from: Radtke et al. 2016. LegacyTreeData: A national database of detailed tree measurements for volume, weight, and physical properties. In: Stanton SM, Christensen GA, (eds). Proceedings of Forest Inventory and Analysis 2015 Science Symposium, December 8–10, 2015, Portland, OR. GTR-PNW-931, USDA Forest Service, Pacific Northwest Research Station, GTR-PNW-931, Portland, OR, pp. 25–30. | 48 |
| Clark_Cypress_Disk, from: Radtke et al. 2016. LegacyTreeData: A national database of detailed tree measurements for volume, weight, and physical properties. In: Stanton SM, Christensen GA, (eds). Proceedings of Forest Inventory and Analysis 2015 Science Symposium, December 8–10, 2015, Portland, OR. GTR-PNW-931, USDA Forest Service, Pacific Northwest Research Station, GTR-PNW-931, Portland, OR, pp. 25–30. | 116 |
| Clark_Cypress, from: Radtke et al. 2016. LegacyTreeData: A national database of detailed tree measurements for volume, weight, and physical properties. In: Stanton SM, Christensen GA, (eds). Proceedings of Forest Inventory and Analysis 2015 Science Symposium, December 8–10, 2015, Portland, OR. GTR-PNW-931, USDA Forest Service, Pacific Northwest Research Station, GTR-PNW-931, Portland, OR, pp. 25–30. | 144 |
| Clark_IP, from: Radtke et al. 2016. LegacyTreeData: A national database of detailed tree measurements for volume, weight, and physical properties. In: Stanton SM, Christensen GA, (eds). Proceedings of Forest Inventory and Analysis 2015 Science Symposium, December 8–10, 2015, Portland, OR. GTR-PNW-931, USDA Forest Service, Pacific Northwest Research Station, GTR-PNW-931, Portland, OR, pp. 25–30. | 264 |
| Clark_NCSU, from: Radtke et al. 2016. LegacyTreeData: A national database of detailed tree measurements for volume, weight, and physical properties. In: Stanton SM, Christensen GA, (eds). Proceedings of Forest Inventory and Analysis 2015 Science Symposium, December 8–10, 2015, Portland, OR. GTR-PNW-931, USDA Forest Service, Pacific Northwest Research Station, GTR-PNW-931, Portland, OR, pp. 25–30. | 140 |
| Clark_Slsh_Complete_Tree 1977, from: Radtke et al. 2016. LegacyTreeData: A national database of detailed tree measurements for volume, weight, and physical properties. In: Stanton SM, Christensen GA, (eds). Proceedings of Forest Inventory and Analysis 2015 Science Symposium, December 8–10, 2015, Portland, OR. GTR-PNW-931, USDA Forest Service, Pacific Northwest Research Station, GTR-PNW-931, Portland, OR, pp. 25–30. | 132 |
| Clark_Sumter, from: Radtke et al. 2016. LegacyTreeData: A national database of detailed tree measurements for volume, weight, and physical properties. In: Stanton SM, Christensen GA, (eds). Proceedings of Forest Inventory and Analysis 2015 Science Symposium, December 8–10, 2015, Portland, OR. GTR-PNW-931, USDA Forest Service, Pacific Northwest Research Station, GTR-PNW-931, Portland, OR, pp. 25–30. | 60 |
| Clark_WPPP, from: Radtke et al. 2016. LegacyTreeData: A national database of detailed tree measurements for volume, weight, and physical properties. In: Stanton SM, Christensen GA, (eds). Proceedings of Forest Inventory and Analysis 2015 Science Symposium, December 8–10, 2015, Portland, OR. GTR-PNW-931, USDA Forest Service, Pacific Northwest Research Station, GTR-PNW-931, Portland, OR, pp. 25–30. | 60 |
| Clark, from: Radtke et al. 2016. LegacyTreeData: A national database of detailed tree measurements for volume, weight, and physical properties. In: Stanton SM, Christensen GA, (eds). Proceedings of Forest Inventory and Analysis 2015 Science Symposium, December 8–10, 2015, Portland, OR. GTR-PNW-931, USDA Forest Service, Pacific Northwest Research Station, GTR-PNW-931, Portland, OR, pp. 25–30. | 1020 |
| Cochard H, Casella E, Mencuccini M. 2007 Xylem vulnerability to cavitation varies among poplar and willow clones and correlates with yield. Tree Physiology 27: 1761-1767 | 19 |
| Cochard H, Damour G, Bodet C, Tharwat I, Poirier M, Améglio T. 2005. Evaluation of a new centrifuge technique for rapid generation of xylem vulnerability curves. Physiologia Plantarum 124: 410-418. | 2 |
| Cochard H. 2006. Cavitation in trees. CR Physique 7: 1018-1126. | 37 |
| Cochard, H. 1992. Vulnerability of several conifers to air embolism. Tree Physiology 11: 73:83. | 1 |
| Colgan et al. 2013. Structural relationships between form factor, wood density, and biomass in African savanna woodlands. Trees 28, 91-102. | 10 |
| compiled by M.L. Cause, E.J. Rudder and W.T. Kynaston 1989. Queensland Timbers: Their nomenclature, Density and Lyctid Susceptibility.. Department of Forestry, QueenslandIn: Ilic, J., Boland, D., McDonald, M., Downes, G. and Blakemore, P. 2000. Woody density phase 1. State of knowledge. National carbon accounting system. Technical Report 18 Australian Greenhouse Office, Canberra, Australia. | 1093 |
| Cornwell and Ackerly 2009, taken from Nelson et al. 2020, and data from Nelson et al. 2020. The Role of Climate Niche, Geofloristic History, Habitat Preference, and Allometry on Wood Density within a California Plant Community. Forests 11(1), 105. | 9 |
| Cornwell and Ackerly 2009, taken from Nelson et al. 2020. The Role of Climate Niche, Geofloristic History, Habitat Preference, and Allometry on Wood Density within a California Plant Community. Forests 11(1), 105. | 7 |
| Costa, J.C.A. 2004. Fixação de carbono e produção de biomassa pela cupiriva (Tapirira guianensis Aubl.), em um fragmento manejado de mata atlântica, município de Goiana-PE. Dissertação (Mestrado). Universidade Federal Rural do Pernambuco, Recife. 109p. | 24 |
| Costa, T. G., Bianchi, M. L., Protásio, T. P., Trugilho, P. F., & Pereira, A. J. (2014). Qualidade da madeira de cinco espécies de ocorrência no cerrado para produção de carvão vegetal. Cerne 20(1), 37-46. | 5 |
| Crivellaro and Schweingruber 2013. Atlas of Wood, Bark and Pith Anatomy of Eastern Mediterranean Trees and Shrub. With a Special Focus on Cyprus. Springer. | 259 |
| CUNHA, M.P.S.C.; PONTES, C.L.F.; CRUZ, I. A.; CABRAL, M. T. F. D.; CUNHA NETO, Z.B.; BARBOSA, A.P.R. Estudo químico de 55 espécies lenhosas para geração de energia em caldeiras. In: 3o encontro Brasileiro em madeiras e em estruturas de madeira: Anais, v.2, p. 93-121, São Carlos, 1989. | 51 |
| da Páscoa et al. 2020. How many trees and samples are adequate for estimating wood-specific gravity across different tropical forests? Trees 34, 1383–1395. | 487 |
| Dadswell H.E. 1972. The anatomy of Eucalypt Woods. CSIROIn: Ilic, J., Boland, D., McDonald, M., Downes, G. and Blakemore, P. 2000. Woody density phase 1. State of knowledge. National carbon accounting system. Technical Report 18 Australian Greenhouse Office, Canberra, Australia. | 22 |
| Damásio, R.A.P., Pereira, B.L.C., Oliveira, A. C., Cardoso, M. T., Vital, B. R., & Carvalho, A. M. L. M. (2013). Caracterização anatômica e qualidade do carvão vegetal da madeira de pau-jacaré (Piptadenia gonoacantha). Pesquisa Florestal Brasileira, 33(75), 261-267. | 1 |
| Danilin et al. 2015, Schepaschenko et al. (2017) Sci. Data 4:170070. DOI: 10.1038/sdata.2017.70 | 49 |
| De Ridder, M., Van den Bulcke, J., Vansteenkiste, D., Van Loo, D., Dierick, M., Masschaele, B., ... & Van Hoorebeke, L. (2010). High-resolution proxies for wood density variations in Terminalia superba. Annals of botany, 107(2), 293-302. | 1 |
| Desch, H.E. (1941) Dipterocarp Timbers of the Malay Peninsula. Malaysian Forest Records No. 14. Caxton Press, Kuala Lumpur Malaysia. / Desch, H.E. (1941 and 1954) Manual of Malayan timbers. Malaysian Forest Records No. 15, Vols 1 & 2, Caxton Press, Kuala Lumpur Malaysia. | 860 |
| Detienne, P., Jacquet P., and Mariaux, A. 1982. Manuel d'Identification des Bois Tropicaux, Tome 3 and Pierre Detienne, P. and Jacquet, P. 1983. Atlas d'Identification des Bois de l'Amazonie et des Régions Voisines, both: Centre Technique Forestier Tropical, Nogent-sur-Marne | 74 |
| Devillez et al. 1973, Schepaschenko et al. (2017) Sci. Data 4:170070. DOI: 10.1038/sdata.2017.70 | 1 |

| | |
|---|---|
| Dimitri, M.J. and Biloni, J.S. 1973. Libro del árbol. Tomo 1. Esencias forestales indígenas de la Argentina de aplicación ornamental. Editorial Celulosa Argentina; S.A. Leonardis R.F.J. 1975. Libro del árbol. Tomo 2. Esencias Forestales indígenas de la Argentina de aplicación industrial. Editorial Celulosa Argentina S. A. | 92 |
| Djomo et al. 2017. Variation of wood density in tropical rainforest trees. Journal of Forests 4(2), 16-26. | 30 |
| Do Nascimento, C.C., 1993. Variabilidade da densidade basica e de propriedades mecanicas de madeiras da Amazonia. Masters thesis in Forestry Sciences, Universidade de São Paulo, Escola Superior de Agricultura 'Luiz de Queiroz', Piracicaba, SP Brazil, 129. From Fearnside (1997) Forest Ecology and Management. | 63 |
| Domec J-C., Scholz F.G., Bucci S.J., Meinzer F.C. Goldstein G., Villalobos-Vega R. 2006. Diurnal and seasonal variation in root xylem embolism in tropical savanna woody species: relationship to stomatal behavior and plant water potential. Plant Cell and Environment 29: 26-35. | 4 |
| Domec J.C. & Gartner B.L. 2002. Age- and position-related changes in hydraulic versus mechanical dysfunction of xylem: inferring the design criteria for Douglas-fir wood structure. Tree Physiology 22: 91-104. | 1 |
| Domec J.C. & Gartner B.L. 2002. How do water transport and water storage differ in coniferous earlywood and latewood. Journal of Experimental Botany 53: 2369-2379. | 2 |
| Domec J.C., Schäfer K., Oren R., Kim H.S., McCarthy H.R. 2010. Variable conductivity and embolism in roots and branches of four contrasting tree species and their impacts on whole-plant hydraulic performance under future atmospheric CO_ concentration. Tree Physiology 30: 1001-1015. | 9 |
| Domec, J.-C., B. Lachenbruch, and F. C. Meinzer. 2006. Bordered pit structure and function determine spatial patterns of air-seeding thresholds in xylem of Douglas-fir (Pseudotsuga menziesii; Pinaceae) trees. American Journal of Botany 93: 1588-1600; Domec J.-C., Warren J.M., Meinzer F.C., Lanchenbruch B. 2009. Safety factors for xylem failure by implosion and air-seeding within roots, trunks and branches of young and old conifer trees. IAWA Journal 30: 101-120. | 6 |
| Droste zu Hulshoff 1969, Schepaschenko et al. (2017) Sci. Data 4:170070. DOI: 10.1038/sdata.2017.70 | 4 |
| Dwianto et al. 2020. IOP Conf. Ser.: Earth Environ. Sci 591 012040 | 19 |
| Dylis and Nosova 1977, Schepaschenko et al. (2017) Sci. Data 4:170070. DOI: 10.1038/sdata.2017.70 | 3 |
| Easdale, T. - Universidad Tucuman, Unpublished data. | 60 |
| Edwards, EJ. 2006. Correlated evolution of stem and leaf hydraulic traits in Pereskia (Cactaceae). New Phytologist 172: 479-489 | 7 |
| Ellick 2015. The carbon sequestration potential of Commidendrum robustum Roxb. (DC.) within the Millennium Forest restoration site, St Helena Island. Master Thesis. York University. https://etheses.whiterose.ac.uk/9337/ | 1 |
| Fairbairn E. 1999. Australian Timbers: Volume two. Department of Natural Resources, QueenslandIn: Ilic, J., Boland, D., McDonald, M., Downes, G. and Blakemore, P. 2000. Woody density phase 1. State of knowledge. National carbon accounting system. Technical Report 18 Australian Greenhouse Office, Canberra, Australia. | 11 |
| Falster DS (2003) Plant height strategies. MSc thesis, Macquarie University, Australia | 130 |
| Falster DS, Westoby M (2005) Alternative height strategies among 45 dicot rain forest species from tropical Queensland, Australia. Journal of Ecology 93: 521-535. 10.1111/j.0022-0477.2005.00992.x | 310 |
| Fan Z.-X., Zhang S.-B., Hao G.-Y., Slik J.W.F., Cao K.-F. 2012. Hydraulic conductivity traits predict growth rates and adult stature of 40 Asian tropical tree species better than wood density. Journal of Ecology 100: 732-741. | 42 |
| Fanshawe, D.B. 1961. Forest products of British Guiana I: principal timbers. Forestry Bulletin (New Series). Forest Department, Georgetown, British Guiana By H ter Steege. | 118 |
| Farfan-Rios et al. 2019: Forest responses to climate change along an Andes-to-Amazon elevational gradient. Dissertation. Wake Forest University Graduate School of Arts and Sciences 2019. Data extracted from Appendix and subset to non-genus averages and species not yet found in wood density database (raw data values were not available). | 218 |
| Farias et a. 2020. Dataset on wood density of trees in ecotone forests in Northern Brazilian Amazonia. Data in Brief 30, 105378. | 1360 |
| Fathi 2014. Structural and mechanical properties of the wood from coconut palms, oil palms and date palms. Dissertation. Universität Hamburg. Values provided for different sections, weighted by relative area. | 4 |
| Favrichon, V. 1994. Classification des espèces arborées en groupes fontionnels en vue de la réalisation d'un modèle de dynamique de peuplement en forêt Guyannaise. Revue d'Ecologie Terre et Vie 49:379-402. V. Favrichon, Modele matriciel deterministe en temps discret. Application a l'etude d'un peupleument forestier tropical humide. Unpublished PhD thesis, Universite Claude Bernard, Lyon I, 1995. | 50 |
| Fayolle et al. (2013) FEM | 357 |
| Feger et al. 1991, Schepaschenko et al. (2017) Sci. Data 4:170070. DOI: 10.1038/sdata.2017.70 | 1 |
| Feng Yang 1985 1995, Schepaschenko et al. (2017) Sci. Data 4:170070. DOI: 10.1038/sdata.2017.70 | 3 |
| Fernandes Neto et al. 2019. Alternative functional trajectories along succession after different land uses in central Amazonia. Journal of Applied Ecology 56(11), 2472-2481. | 168 |
| Ferraz, I.D.K., Leal Filho, N., Imakawa, A.M., Varela, V.P., and Pina-Rodrigues, F.C.M. 2004. Caracteristicas basicas para um agrupamento ecologico preliminar de especies madeireiras da floresta de terra firme da Amazonia Central. Acta Amazonica 34: 621-633. | 58 |
| Flynn Jr., J.H. and Holder, C.D. 2001. A Guide to Useful Woods of the World. 2nd ed. Forest Products Society, Madison. | 40 |
| Forest Products Laboratory. 1987 and 1999. Wood Handbook: Wood as an Engineering Material. General Technical Report. FPL-GTR-113. USDA. | 7 |
| França, M.B. 2002. Modelagem de biomassa através do padrão espectral no sudoeste da Amazônia. Dissertação (Mestrado). Instituto Nacional de Pesquisas da Amazônia/Fundação Universidade Federal do Amazonas, Manaus, Amazonas, Brazil. 106pp. | 128 |
| Freyburger, C., Longuetaud, F., Mothe, F., Constant, T., & Leban, J. M. (2009). Measuring wood density by means of X-ray computer tomography. Annals of forest science, 66(8), 804. | 37 |
| Fu, P-L., Jiang, Y-J., Wang, A-Y., Brodribb, T.J., Zhang, J-L., Zhu, S-D. and Cao, K-F. 2012. Stem hydraulic traits and leaf water-stress tolerance are co-ordinated with the leaf phenology of angiosperm trees in an Asian tropical dry karst forest. Annals of Botany 110: 189-99. | 11 |
| Fuentes-Salinas et al. 2008: Características tecnológicas de 16 maderas del estado de Tamaulipas, que influyen en la fabricación de tableros de partículas y de fibras. Revista Chapingo 14, 65-71. | 16 |
| Fujiwara et al. 2007. Basic densities as a parameter for estimating the amount of carbon removal by forests and their variation. Bulletin of FFPRI 405, 215 - 226. | 59 |
| Gabdelkhakov 2005, Schepaschenko et al. (2017) Sci. Data 4:170070. DOI: 10.1038/sdata.2017.70 | 5 |
| Gabdelkhakov 2015, Schepaschenko et al. (2017) Sci. Data 4:170070. DOI: 10.1038/sdata.2017.70 | 186 |
| Ganivet et al. 2019: Ecological strategies of tree species in the laurel forest of Tenerife (Canary Islands): an insight into cloud forest natural dynamics using long-term monitoring data. European Journal of Forest Research 138, 93-110. | 15 |
| García Esteban et al. 2020. Characterisation of Pinus canariensis C.Sm. ex. DC. Sawn Timber from Reforested Trees on the Island of Tenerife, Spain. Forests 11, 769. | 1 |

| | |
|---|---|
| Gardiner, B., Leban, J. M., Auty, D., & Simpson, H. (2011). Models for predicting wood density of British-grown Sitka spruce. Forestry, 84(2), 119-132. | 5 |
| Gartner, B. L. 1991. Stem hydraulic properties of vines vs. shrubs of western poison oak, Toxicodendron diversilobum. Oecologia 87: 180-189. | 2 |
| Gartner, B. L., S. H. Bullock, H. A. Mooney, V. B. Brown, and J. L. Whitbeck. 1990. Water transport properties of vine and tree stems in a tropical deciduous forest. American Journal of Botany 77: 742-749. | 20 |
| Gazel, M. 1983. Croissance des arbres et productivite des peuplements en foret dense equatoriale de Guyane. Unpublished report of the Office National des Forets. | 147 |
| Genet, A., Auty, D., Achim, A., Bernier, M., Pothier, D., & Cogliastro, A. (2013). Consequences of faster growth for wood density in northern red oak (Quercus rubra Liebl.). Forestry, 86(1), 99-110. | 2 |
| Gérard, J., Miller, R.B. and ter Welle, B.J.H. 1996. Major timber trees of Guyana: timber characteristics and utilization. Tropenbos Series 15, Tropenbos Foundation, Wageningen, The Netherlands. | 38 |
| Gillerot et al. 2018. Inter- and intraspecific variation in mangrove carbon fraction and wood specific gravity in Gazi Bay, Kenya. Ecosphere 9(6), e02306. | 12 |
| Gimenez, A.M. and Moglia, J,G. Arboles del Chaco Argentino. 2003. Guia para el Reconocimiento Dendrologico. Facultad de Ciencias Forestales, Universidad Nacional de Santiago del Estero. Santiago del Estero, Argentina, 307 pp. | 74 |
| Ginoga, B. and Karnasudirdja, S. 1978. Sifat Mekanis Sepuluh Jenis Kayu Indonesia. Laporan LPHH Badan Penelitian dan Pengembangan Pertanian. Departemen Pertanian. Bogor. 1-10 pp. | 7 |
| Ginoga, B., Hadjib, N. and Karna Sudjirdja, S. 1982. Sifat Fisis dan Mekanis Beberapa Jenis Kayu Indonesia Bagian 10. Laporan BPHH No. 162. | 19 |
| Ginoga, B., Hadjib, N. and Karnasudirdja, S. 1980. Sifat Fisis dan Mekanis beberapa Jenis Kayu Indonesia Bagian IX. Laporan BPHH No. 153. | 18 |
| Gleason et al. 2018. Shoot growth of woody trees and shrubs is predicted by maximum plant height and associated traits. Functional Ecology 32(2), 247-259. | 44 |
| Gleason, S.M., Butler, D.W., Zieminska, K., Waryszak, P. andWestoby M. 2012. Stem xylem conductivity is key to plant water balance across Australian angiosperm species. Functional Ecology 26: 343-352. doi: 10.1111/j.1365-2435.2012.01962.x | 120 |
| Glover_1981 1981, from: Radtke et al. 2016. LegacyTreeData: A national database of detailed tree measurements for volume, weight, and physical properties. In: Stanton SM, Christensen GA, (eds). Proceedings of Forest Inventory and Analysis 2015 Science Symposium, December 8–10, 2015, Portland, OR. GTR-PNW-931, USDA Forest Service, Pacific Northwest Research Station, GTR-PNW-931, Portland, OR, pp. 25–30. | 124 |
| Goldsmith, B. and D.T. Carter. 1981. The indigenous timbers of Zimbabwe. The Zimbabwe Bulletin of Forestry Research No. 9:x, 406 pp. | 260 |
| Gomes et al. 2021. Functional traits and symbiotic associations of geoxyles and trees explain the dominance of detarioid legumes in miombo ecosystems. New Phytologist 230(2), 510-520. | 25 |
| Goussanou et al. 2016. Specific and generic stem biomass and volume models of tree species in a West African tropical semi-deciduous forest. Silva Fennica 50(2), article id 1474. https://doi.org/10.14214/sf.1474 | 18 |
| Green, D.W., Winandy, J.E. and Kretschmann, D.E. 1999. Mechanical Properties. Forest Products Laboratory, Chapter 4, 463-509 pp. | 5 |
| Gueneau, P. and Gueneau, D. 1969. Proprietes physiques et mecaniques des bois malgaches. Centre Technique Forestier Tropical, Madagascar. P. and Gueneau, D. 1969. | 24 |
| Gutierrez Oliva, A. and Plaza Pulgar, F. 1967. Caracteristicas fisico-mecanicas de las maderas espanolas. Ministerio de Agricultura, Direccion general de montes, caza y pesca fluvial, Instituto Forestal de Investigaciones y Experiencias. Madrid. 10 3 pp., A. and Plaza Pulgar, F. 1967. | 62 |
| Gutierrez Rojas, V.H. and Sandova, l. J.S. (sin fecha). Informacion Technica para el Procesamiento Industrial de 134 Especies Maderables de Bolivia. Serie Tecnica XII. Ministerio de Desarrollo Sostenible y Planificacion. Santa Cruz, La Paz, Bolivia. 352 pp. | 126 |
| Guzman et al. 2016: Trade-offs between water transport capacity and drought resistance in neotropical canopy liana and tree species. Tree physiology 37, 1404-1414. | 12 |
| Hacke U.G. & Jansen S. 2009. Embolism resistance of three boreal conifer species varies with pit structure. New Phytologist 182: 675–686. | 18 |
| Hacke U.G., Sperry J.S., Field T.S., Sano Y., Sikkema H. & Pittermann J. 2007. Water transport in vesselless angiosperms: conducting efficiency and cavitation safety. Int. J. Plant Sci. 168(8):1113–1126. | 7 |
| Hacke, U. G., J. S. Sperry, and J. Pittermann. 2000. Drought experience and cavitation resistance in six desert shrubs of the Great Basin, Utah. Basic and Applied Ecology. 1:31–41. | 9 |
| Hailu Ubuy et al. 2018. Variation in wood basic density within and between tree species and site conditions of exclosures in Tigray, northern Ethiopia. Trees 32, 967–983. | 50 |
| Hamada, J., Pétrissans, A., Mothe, F., Ruelle, J., Pétrissans, M., & Gérardin, P. (2016). Variations in the natural density of European oak wood affect thermal degradation during thermal modification. Annals of forest science, 73(2), 277-286. | 5 |
| Hao G-Z. Hoffmann, W.A., Scholz, F.G., Bucci, S.J. Meinzer, F.C. Franco, A.C. Cao, K-F. and Goldstein, G. 2008. Stem and leaf hydraulic congeneric tree species from adjacent tropical savanna and forest ecosystems. Oecologia 155: 405-415 | 12 |
| Hao G.-Y., Sack L., Wang A.-Y., Cao K.-F. & Goldstein G. (2010) Differentiation of leaf water flux and drought tolerance traits in hemiepiphytic and non-hemiepiphytic Ficus tree species. Functional Ecology 486 24: 731–740. Hao G.-Y., Goldstein G., Sack L., Holbrook N.M., Liu Z.-H., Wang A.-Y., Harrison R.D., Su Z.-H., Cao K-F. (2011) Ecology of hemiepiphytism in fig species is based on evolutionary correlation of hydraulics and carbon economy. Ecology 92: 2117-2130. | 19 |
| Harrison_et_al 2009, from: Radtke et al. 2016. LegacyTreeData: A national database of detailed tree measurements for volume, weight, and physical properties. In: Stanton SM, Christensen GA, (eds). Proceedings of Forest Inventory and Analysis 2015 Science Symposium, December 8–10, 2015, Portland, OR. GTR-PNW-931, USDA Forest Service, Pacific Northwest Research Station, GTR-PNW-931, Portland, OR, pp. 25–30. | 84 |
| He and Deane 2016. The relationship between trunk- and twigwood density shifts with tree size and species stature. Forest Ecology and Management 372, 137–142. | 140 |
| Heady, R.D., Banks, J.G. and Evans, P.D. 2002. Wood Anatomy of Wollemi Pine (Wollemia nobilis, Araucariaceae). IAWA Journal 23(4): 339-357. | 1 |
| Henry, M., Besnard, A., Asante, W. A., Eshun, J., Adu-Bredu, S., Valentini, R., ... & Saint-André, L. (2010). Wood density, phytomass variations within and among trees, and allometric equations in a tropical rainforest of Africa. Forest Ecology and Management, 260(8), 1375-1388. | 42 |
| Hervé, V., Mothe, F., Freyburger, C., Gelhaye, E., & Frey-Klett, P. (2014). Density mapping of decaying wood using X-ray computed tomography. International Biodeterioration & Biodegradation, 86, 358-363. | 1 |
| Hidayat & Simpson 1994: Use of Green Moisture Content and Basic Specific Gravity to Group Tropical Woods for Kiln Drying. Forest Products Laboratory Research Note FPL-RN-0263. Values derived from: Ali et al. 1968 | 14 |
| Hidayat & Simpson 1994: Use of Green Moisture Content and Basic Specific Gravity to Group Tropical Woods for Kiln Drying. Forest Products Laboratory Research Note FPL-RN-0263. Values derived from: Anonymous 1990 | 10 |
| Hidayat & Simpson 1994: Use of Green Moisture Content and Basic Specific Gravity to Group Tropical Woods for Kiln Drying. Forest Products Laboratory Research Note FPL-RN-0263. Values derived from: Arostegui 1982 | 59 |

| | |
|---|---|
| Hidayat & Simpson 1994: Use of Green Moisture Content and Basic Specific Gravity to Group Tropical Woods for Kiln Drying. Forest Products Laboratory Research Note FPL-RN-0263. Values derived from: Brazier & Franklin 1967 | 49 |
| Hidayat & Simpson 1994: Use of Green Moisture Content and Basic Specific Gravity to Group Tropical Woods for Kiln Drying. Forest Products Laboratory Research Note FPL-RN-0263. Values derived from: Darkwa 1973 | 8 |
| Hidayat & Simpson 1994: Use of Green Moisture Content and Basic Specific Gravity to Group Tropical Woods for Kiln Drying. Forest Products Laboratory Research Note FPL-RN-0263. Values derived from: Echenique-Manrique 1971 | 36 |
| Hidayat & Simpson 1994: Use of Green Moisture Content and Basic Specific Gravity to Group Tropical Woods for Kiln Drying. Forest Products Laboratory Research Note FPL-RN-0263. Values derived from: Hon & Pun 1965 | 128 |
| Hidayat & Simpson 1994: Use of Green Moisture Content and Basic Specific Gravity to Group Tropical Woods for Kiln Drying. Forest Products Laboratory Research Note FPL-RN-0263. Values derived from: Karasujana & Martawijaya 1979 | 254 |
| Hidayat & Simpson 1994: Use of Green Moisture Content and Basic Specific Gravity to Group Tropical Woods for Kiln Drying. Forest Products Laboratory Research Note FPL-RN-0263. Values derived from: Limaye & Sen 1956 | 217 |
| Hidayat & Simpson 1994: Use of Green Moisture Content and Basic Specific Gravity to Group Tropical Woods for Kiln Drying. Forest Products Laboratory Research Note FPL-RN-0263. Values derived from: Okoh 1971 | 14 |
| Hidayat & Simpson 1994: Use of Green Moisture Content and Basic Specific Gravity to Group Tropical Woods for Kiln Drying. Forest Products Laboratory Research Note FPL-RN-0263. Values derived from: Rocafort & Siopongco 1974 | 214 |
| Hidayat & Simpson 1994: Use of Green Moisture Content and Basic Specific Gravity to Group Tropical Woods for Kiln Drying. Forest Products Laboratory Research Note FPL-RN-0263. Values derived from: Sharma 1975 | 1 |
| Hidayat & Simpson 1994: Use of Green Moisture Content and Basic Specific Gravity to Group Tropical Woods for Kiln Drying. Forest Products Laboratory Research Note FPL-RN-0263. Values derived from: van Vuuren et al. 1978 | 39 |
| Hidayat & Simpson 1994: Use of Green Moisture Content and Basic Specific Gravity to Group Tropical Woods for Kiln Drying. Forest Products Laboratory Research Note FPL-RN-0263. Values derived from: Vilela 1969 | 132 |
| Hidayat & Simpson 1994: Use of Green Moisture Content and Basic Specific Gravity to Group Tropical Woods for Kiln Drying. Forest Products Laboratory Research Note FPL-RN-0263. Values derived from: Vilela 1974 | 106 |
| Holiaka D.M., Bilous A.M., Holiaka M.A. (2018). Live biomass of willow bushes in natural phytocenoses Chernihiv Polissia. Kyiv: NULES of Ukraine. ISBN 978-617-7630-47-9 | 55 |
| Huerta Crespo, J. and Becerra Martinez, J. 1982. Anatomia Macroscopica y Algunas Caracteristicas Fisicas de Diecisiete Maderas Tropicales Mexicanas. Boletin Divulgativo No. 46, Instituto Nacional de Investigaciones Forestales, Mexico. 61 pp. | 6 |
| Hultine, K. R., D. F. Koepke, W. T. Pockman, A. Fravolini, J. S. Sperry, and D. G. Williams. 2006. Influence of soil texture on hydraulic properties and water relations of a dominant warm-desert phreatophyte. Tree Physiology 26:313-323. | 4 |
| IBDF. 1981. Madeiras de Amazônia, caracteristicas e utilização. Vol 1. Floresta Nacional do Tapajos. Instituto Brasileiro de Desenvolvimento Florestal, Brasilia, DF Brasil, 113 pp. In Fearnside, P.M. 1997. Wood density for estimating forest biomass in Brazilian Amazonia. Forest Ecology and Management 90: 59-87. | 8 |
| IBDF. 1983. Potencial madeireira do Grande Carajas, Instituto Brasileiro de Desenvolvimento Florestal, Brasilia, DF Brasil, 134 pp. In Fearnside, P.M. 1997. Wood density for estimating forest biomass in Brazilian Amazonia. Forest Ecology and Management 90: 59-87. | 31 |
| IBDF. 1988. Madeiras de Amazônia, caracteristicas e utilização. Estação experimental de Curua-Una. Vol 2. Instituto Brasileiro de Desenvolvimento Florestal, Brasilia, DF Brasil, 134 pp. In Fearnside, P.M. 1997. Wood density for estimating forest biomass in Brazilian Amazonia. Forest Ecology and Management 90: 59-87. | 38 |
| Iida, Y., Poorter, L., Sterck, F. J., Kassim, A. R., Kubo, T., Potts, M. D., & Kohyama, T. S. (2012). Wood density explains architectural differentiation across 145 co-occurring tropical tree species. Functional Ecology, 26(1), 274-282. | 37 |
| IMANA-ENCINAS, J. et al. 2012. Florística, volume e biomassa lenhosa de um fragmento de Mata Atlántica no municipio de Santa Maria de Jetibá - Espirito Santo. Floresta 42(3): 565 - 576. | 73 |
| Inga, P.R. and Castillo, M.U. 1987. Caracteristicas fisico-quimicas de la madera y carbon de once especies forestales de la Amazonia peruana. Revista forestal del Peru 14:62-73. By T Baker. | 11 |
| INPA 1991. Catalogo de madeiras da Amazônia. Instituto Nacional de Pesquisas da Amazônia, Coodenação de Pesquisas em Produtos Forestais, Manaus, AM Brasil, 163 pp. In Fearnside, P.M. 1997. Wood density for estimating forest biomass in Brazilian Amazonia. Forest Ecology and Management 90: 59-87. | 43 |
| Instituto Nacional de Tecnologia Industrial (INTI), Argentina 2003. Densidad de maderas (Kg/m3). Prepared by: Ing. Ftal. María Elena Atencia. https://www.inti.gob.ar/publicaciones/descargac/365, last accessed on 15/07/2019. | 389 |
| Iogna PA, Bucci SJ, Scholz FG, Goldstein G (2011) Water relations and hydraulic architecture of two Patagonian steppe shrubs: effect of slope orientation and microclimate. J Arid Environ 75: 763–772. | 6 |
| Ivanchikov 1971, Schepaschenko et al. (2017) Sci. Data 4:170070. DOI: 10.1038/sdata.2017.70 | 9 |
| Jacobsen A.L., Agenbag L., Esler K.J., Pratt R.B., Ewers F.W., Davis S.D. 2007. Xylem density, biomechanics and anatomical traits correlate with water stress in 17 evergreen shrub species of the Mediterranean-type climate region of South Africa. Journal of Ecology 95: 171-183. | 19 |
| Jacobsen A.L., Ewers F.W., Pratt R.B., Paddock W.A. III, Davis S.D. 2005. Do xylem fibers affect vessel cavitation resistance? Plant Physiology 139: 546-556. | 6 |
| Jacobsen AL, Pratt RB, Davis SD, Ewers FW. 2008. Comparative community physiology: non-convergence in water relations among three semi-arid shrub communities. New Phytologist 180: 100-113. | 29 |
| Jacobsen AL, Pratt RB, Ewers FW, Davis SD. 2007. Cavitation resistance among twenty-six chaparral species of southern California. Ecological Monographs 77: 99-115. | 26 |
| Jasinska AK, Alber M, Tullus A, Rahi M, Sellin A (2015) Impact of elevated atmospheric humidity on anatomical and hydraulic traits of xylem in hybrid aspen. Funct Plant Biol 42:565–578. doi:10.1071/FP14224 | 2 |
| Jati et al. 2014. Densidade da madeira de árvores em savanas do norte da Amazônia brasileira. Acta Amazonica 44 (1), 79-86. | 225 |
| Jenkins, K.L. and Coomes, D.A. unpublished data. | 46 |
| JMPROV, from: Radtke et al. 2016. LegacyTreeData: A national database of detailed tree measurements for volume, weight, and physical properties. In: Stanton SM, Christensen GA, (eds). Proceedings of Forest Inventory and Analysis 2015 Science Symposium, December 8–10, 2015, Portland, OR. GTR-PNW-931, USDA Forest Service, Pacific Northwest Research Station, GTR-PNW-931, Portland, OR, pp. 25–30. | 288 |
| Johnson D.M., Domec J.C., Woodruff D.R., McCulloh K.A., Meinzer F.C. 2013. Contrasting hydraulic strategies in two tropical lianas and their host trees. American Journal of Botany 100: 374-83. | 5 |

| | |
|---|---|
| K.R. Bootle 1983. Wood in Australia Types, properties and uses. In: Ilic, J., Boland, D., McDonald, M., Downes, G. and Blakemore, P. 2000. Woody density phase 1. State of knowledge. National carbon accounting system. Technical Report 18 Australian Greenhouse Office, Canberra, Australia. | 1 |
| Kanawjia et al. 2013. Specifi c gravity of some woody species in the Srinagar Valley of the Garhwal Himalayas, India. For. Sci. Pract. 15(1), 85-88. | 31 |
| Karizumi 1974, Schepaschenko et al. (2017) Sci. Data 4:170070. DOI: 10.1038/sdata.2017.70 | 76 |
| Kasatkin et al. 2015a, Schepaschenko et al. (2017) Sci. Data 4:170070. DOI: 10.1038/sdata.2017.70 | 43 |
| Kasatkin et al. 2015b, Schepaschenko et al. (2017) Sci. Data 4:170070. DOI: 10.1038/sdata.2017.70 | 49 |
| Kasatkin et al. 2016, Schepaschenko et al. (2017) Sci. Data 4:170070. DOI: 10.1038/sdata.2017.70 | 39 |
| Kazimirov and Morozova 1973, Schepaschenko et al. (2017) Sci. Data 4:170070. DOI: 10.1038/sdata.2017.70 | 61 |
| Kazimirov et al. 1977, Schepaschenko et al. (2017) Sci. Data 4:170070. DOI: 10.1038/sdata.2017.70 | 21 |
| Kazimirov et al. 1978, Schepaschenko et al. (2017) Sci. Data 4:170070. DOI: 10.1038/sdata.2017.70 | 35 |
| Keduolhouvonuo and Kumar 2017. Variation in wood specific gravity of selected tree species of Kohima district of Nagaland North Eastern parts of India. Journal of Pharmacognosy and Phytochemistry 6(6), 70-74. | 28 |
| Keenan, F.J. and Tejada, M. 1984. Tropical Timber for Building Materials in the Andean Group Countries of South America. IDRC-TS 49. | 31 |
| Kimura 1963, Schepaschenko et al. (2017) Sci. Data 4:170070. DOI: 10.1038/sdata.2017.70 | 1 |
| Kindermann et al. 2021. Dataset on Woody Aboveground Biomass, Disturbance Losses, and Wood Density from an African Savanna Ecosystem. Mendeley Data, V1, doi: 10.17632/3cs85wd3gb.1. Also preprint: Improving estimation of woody aboveground biomass in drylands by accounting for disturbances and spatial heterogeneity. | 61 |
| King, D.A. 1996. Allometry and life history of tropical trees. Journal of Tropical Ecology 12: 25-44. | 14 |
| Kingston R.S.T. and Risdon C.J.E. 1961. Shrinkage and density of Australian and other South-west Pacific woods. CSIROIn: Ilic, J., Boland, D., McDonald, M., Downes, G. and Blakemore, P. 2000. Woody density phase 1. State of knowledge. National carbon accounting system. Technical Report 18 Australian Greenhouse Office, Canberra, Australia. | 352 |
| Kininmonth, J.A. 1982. Properties and uses of the timbers of West Samoa. Indigenous hardwoods. Forest Research Institute, Rotorua, New Zealand. 57 pp. | 9 |
| Kizha, from: Radtke et al. 2016. LegacyTreeData: A national database of detailed tree measurements for volume, weight, and physical properties. In: Stanton SM, Christensen GA, (eds). Proceedings of Forest Inventory and Analysis 2015 Science Symposium, December 8–10, 2015, Portland, OR. GTR-PNW-931, USDA Forest Service, Pacific Northwest Research Station, GTR-PNW-931, Portland, OR, pp. 25–30. | 52 |
| Kleinschmidt et al. 2020. Successional habitat filtering of rainforest trees is explained by potential growth more than by functional traits. Functional Ecology 34(7), 1438-1447. | 42 |
| Koltunova et al. 2007, Schepaschenko et al. (2017) Sci. Data 4:170070. DOI: 10.1038/sdata.2017.70 | 45 |
| Kostov et al. 1992, Schepaschenko et al. (2017) Sci. Data 4:170070. DOI: 10.1038/sdata.2017.70 | 7 |
| Kotowska, M. M., Hertel, D., Rajab, Y. A., Barus, H., & Schuldt, B. (2015). Patterns in hydraulic architecture from roots to branches in six tropical tree species from cacao agroforestry and their relation to wood density and stem growth. Frontiers in plant science, 6. | 6 |
| Kryn & Fobes 1959: The Woods of Liberia. | 130 |
| Kukachka, B.F. 1970. Properties of imported tropical woods. Forest Products Laboratory Research Paper 125,Forest Service, United States Department of Agriculture. | 51 |
| Kukachka, B.F., McClay, T.A. and Beltranena M.E. 1968. Selected Properties of 52 Timber Species from the Department of El Peten, Guatemala. (also Kukachka, McClay, Beltranena. 1968. Propiedades seleccionadas de 52 especies de madera del Departamento del Peten, Guatemala. Boletin Numero 2, Proyecto de Evaluacion Forestal -- FAO-FYDEP -- Guatemala, C.A.; 88 pp.) | 7 |
| Kusumoto et al. 2014. Functional response of plant communities to clearcutting: management impacts differ between forest vegetation zones. J Appl Ecol, 52: 171-180. https://doi.org/10.1111/1365-2664.12367 | 2516 |
| Lachenbruch, B., Johnson, G. R., Downes, G. M., & Evans, R. (2010). Relationships of density, microfibril angle, and sound velocity with stiffness and strength in mature wood of Douglas-fir. Canadian Journal of Forest Research, 40(1), 55-64. | 1 |
| Lakyda et al. 2018, Schepaschenko et al. (2017) Sci. Data 4:170070. DOI: 10.1038/sdata.2017.70 | 222 |
| Lakyda P.I., Vasylyshyn R.D., Blyshchyk V.I., Lakyda I.P., Bilous A.M., Matushevych L.M., ... Dubrovets B.V. (2020). Experimental data on live biomass of Ukrainian deciduous forests. Kyiv: PC Komprynt LLC. ISBN 978-617-7986-50-8 | 726 |
| Lakyda P.I., Vasylyshyn R.D., Blyshchyk V.I., Lakyda I.P., Terentiev A.Yu., Domashovets H.S., ... Stratii N.V. (2018). Experimental data on live biomass of Ukrainian coniferous forests. Kyiv: PC Komprynt LLC. ISBN 978-966-929-806-5 | 142 |
| Langbour, P., Sébastien, P., and Thibaut, B. 2019. Description of the Cirad wood collection in Montpellier, France, representing eight thousand identified species. BOIS ET FORÈTS DES TROPIQUES n° 339, p.7-16. Available at: / Previously included through: Detienne, P., Jacquet P., and Mariaux, A. 1982. Manuel d'Identification des Bois Tropicaux, Tome 3, Guyane Francaise. Centre Technique Forestier Tropical, Nogent-sur-Marne, France.; Pierre Détienne, P. and Jacquet, P. 1983. Atlas d'Identification des Bois de l'Amazonie et des Régions Voisines. Centre Technique Forestier Tropical, Nogent-sur-Marne, 640 pp. | 19488 |
| Larter et al. 2017. Aridity drove the evolution of extreme embolism resistance and the radiation of conifer genus Callitris. New Phytol 215, 97-112. https://doi.org/10.1111/nph.14545 | 23 |
| Lastra Rivera, J.A. 1986. Compilacion de las propiedades fisico-mecanicas y usos posibles de 178 maderas de Colombia. Bogota. 75 pp. | 15 |
| Lauricio, F.M. and Siopongco, J.D. 1970. Sixth progress report on the mechanical and related properties of Philippine woods. The Philippine Lumberman 16(6): 17-24. | 16 |
| Lavers, G.M. 1983. The Strength Properties of Timber. 3rd ed. Building Research Establishment Report, HMSO, London, 60 pp. | 52 |
| Lawson et al. 2015. Hydrological conditions explain variation in wood density in riparian plants of south-eastern Australia. Journal of Ecology 103(4), 945-956. https://doi.org/10.1111/1365-2745.12408 | 66 |
| Leban et al. 2021. Wood basic density for 125 tree forest species from the French forests, https://doi.org/10.15454/XFOPL1, Portail Data INRAE, V1, UNF:6:RD2eei4/fHwqpfXGUpfbew== [fileUNF] | 125 |
| Lens, F., Sperry, J. S., Christman, M. A., Choat, B., Rabaey, D., & Jansen, S. (2011). Testing hypotheses that link wood anatomy to cavitation resistance and hydraulic conductivity in the genus Acer. New Phytologist, 190(3), 709-723. | 7 |
| Li 2011. Light-use strategies and biomass accumulation of woody species in a subtropical forest in southwest China. Dissertation. University of Zurich, Faculty of Science. | 41 |
| Li et al. 1981, Schepaschenko et al. (2017) Sci. Data 4:170070. DOI: 10.1038/sdata.2017.70 | 2 |
| Li et al. 2015. Are functional traits a good predictor of global change impacts on tree species abundance dynamics in a subtropical forest? Ecology Letters 18(11), 1181–1189. | 48 |

| | |
|---|---|
| Lima, J.A.S. 2009. Biomassa arbórea e estoques de nutrientes em fragmentos florestais da APA Rio São João: o efeito da fragmentação sobre a Mata Atlântica da Baixada Litorânea Fluminense. Tese (Doutorado). Universidade Estadual do Norte Fluminense. Campo dos Goytacazes. 180p. | 24 |
| Liogier, A.H. 1978. Arboles Dominicanos. Academia de Ciencias de la Republica Dominicana, Santo Domingo, Dominican Republic. | 26 |
| Little, E.L., Jr., and F.H. Wadesworth. 1964. Common trees of Puerto Rico and the Virgin Islands, US Department of Agriculture, Agricultural Handbook 249, Superintendent of Documents, US Government Printing Office, Washington DC. | 92 |
| Little, E.L., Jr., R.O. Wodbury, and F.H. Wadesworth. 1974. Trees of Puerto Rico and the Virgin Islands. Second Volume. US Department of Agriculture, Agricultural Handbook 449, Superintendent of Documents, US Government Printing Office, Washington DC. | 23 |
| Litvinova et al. 2009, Schepaschenko et al. (2017) Sci. Data 4:170070. DOI: 10.1038/sdata.2017.70 | 5 |
| Llach Cordero. (no date) Report on a Wood Testing Programme Carried out for UNDP/SF Project 234, Inventory and Forest Demonstrations Panama, Part III, Physical and Mechanical Properties of 113 Species. | 4 |
| Lohrey 1985, from: Radtke et al. 2016. LegacyTreeData: A national database of detailed tree measurements for volume, weight, and physical properties. In: Stanton SM, Christensen GA, (eds). Proceedings of Forest Inventory and Analysis 2015 Science Symposium, December 8–10, 2015, Portland, OR. GTR-PNW-931, USDA Forest Service, Pacific Northwest Research Station, GTR-PNW-931, Portland, OR, pp. 25–30. | 915 |
| Lohrey_direct_seeded, from: Radtke et al. 2016. LegacyTreeData: A national database of detailed tree measurements for volume, weight, and physical properties. In: Stanton SM, Christensen GA, (eds). Proceedings of Forest Inventory and Analysis 2015 Science Symposium, December 8–10, 2015, Portland, OR. GTR-PNW-931, USDA Forest Service, Pacific Northwest Research Station, GTR-PNW-931, Portland, OR, pp. 25–30. | 572 |
| Lorenzi, H. 1992/2000. Arboles brasileiras: Manual de identificação e cultivo de plantas arbóreas nativas do Brasil. Vol.1 & 2. Nova Odessa, SP, Brazil, Instituto Plantarum de Estudos da Flora Ltda. www.plantarum.com.br | 446 |
| Lorenzi, H. 2009. Árvores brasileiras: manual de identificação e cultivo de plantas arbóreas nativas do Brasil, vol. 3 (1st Ed.). Instituto Plantarum, Nova Odessa. | 231 |
| Loureiro, A. A. and Braga Lisboa, P. L. 1979. Madeiras do Município de Aripuana e suas utilidades (Mato Grosso). Acta Amazonica 9(1): 1-79. | 62 |
| Machado, J. L., and M. T. Tyree. 1994. Patterns of hydraulic architecture and water relations of two tropical canopy trees with contrasting leaf phenologies: Ochroma pyramidale and Pseudobombax septenatum. Tree Physiology 14:219–240. | 2 |
| Maeshiro et al. 2013. Using tree functional diversity to evaluate management impacts in a subtropical forest. Ecosphere4(6):70. http://dx.doi.org/10.1890/ES13-00125.1 | 501 |
| Magalhaes et al. 2020. Data on dendrometric parameters, basic wood density, below- and aboveground biomass of tree species from Mangrove, Miombo, Mopane, and Mecrusse woodlands. Data in brief 2020, 105154. | 497 |
| Mahato et al. 2019. Wood Specific Gravity of Temperate Forest Species of Garhwal Himalaya, India. Indian Forester 145 (11), 1035-1038. | 26 |
| Maherali H., Moura C.F., Caldeira M.C., Willson C.J. & Jackson R.B. 2006. Functional coordination between leaf gas exchange and vulnerability to xylem cavitation in temperate forest trees. Plant, Cell and Environment, 29: 571-583. | 14 |
| Maksimov 2003, Schepaschenko et al. (2017) Sci. Data 4:170070. DOI: 10.1038/sdata.2017.70 | 13 |
| Malavassi, I.M.C. 1992. Maderas de Costa Rica: 150 Especies forestales, Editorial de la Universidad de Costa Rica. | 129 |
| Malenko et al. 2015, Schepaschenko et al. (2017) Sci. Data 4:170070. DOI: 10.1038/sdata.2017.70 | 21 |
| Malimbwi and Solberg 1994. Estimation of biomass and volume in miombo woodland at Kitulangalo Forest Reserve, Tanzania. Journal of Tropical Forest Science 7(2), 230-242. | 34 |
| Malkonen 1977, Schepaschenko et al. (2017) Sci. Data 4:170070. DOI: 10.1038/sdata.2017.70 | 1 |
| Mallque, M.A., Yoza, L.Y. and Garcia, A.Q. 1991. Determinacion de las propiedades electricas en seis maderas tropicales. Revista Forestal del Peru, 18:5-21 By T Baker. | 1 |
| Mamers H., Balodis V., Garland C.P., Langfors N.G. and Menz D.N.J., 1991. Kraft pulping of East Gippsland eucalypt regrowth. CSIROIn: Ilic, J., Boland, D., McDonald, M., Downes, G. and Blakemore, P. 2000. Woody density phase 1. State of knowledge. National carbon accounting system. Technical Report 18 Australian Greenhouse Office, Canberra, Australia. | 7 |
| Mamers H., Balodis V., Garland C.P., Langfors N.G., Menz D.N.J., and Chin C.W.J. 1990. An assessment of the kraft pulping properties of residual mature eucalypt roundwood from East Gippsland. CSIROIn: Ilic, J., Boland, D., McDonald, M., Downes, G. and Blakemore, P. 2000. Woody density phase 1. State of knowledge. National carbon accounting system. Technical Report 18 Australian Greenhouse Office, Canberra, Australia. | 6 |
| Mani & Parthasarathy 2007: Above-ground biomass estimation in ten tropical dry evergreen forest sites of peninsular India | 59 |
| Markesteijn and Poorter 2009. Seedling root morphology and biomass allocation of 62 tropical tree species in relation to drought- and shade-tolerance. Journal of Ecology 97, 311-325. https://doi.org/10.1111/j.1365-2745.2008.01466.x | 70 |
| Markwardt, L.J. and Wilson, T.R. 1935. Strength and Related Properties of Woods Grown in the United States. USDA Forest Service, Tech. Bull. No. 479. | 33 |
| Marrs 1980, from: Radtke et al. 2016. LegacyTreeData: A national database of detailed tree measurements for volume, weight, and physical properties. In: Stanton SM, Christensen GA, (eds). Proceedings of Forest Inventory and Analysis 2015 Science Symposium, December 8–10, 2015, Portland, OR. GTR-PNW-931, USDA Forest Service, Pacific Northwest Research Station, GTR-PNW-931, Portland, OR, pp. 25–30. | 129 |
| Martawijaya, A. et al. 1992. Indonesian Wood Atlas Vol. I. and II AFPRDC AFRD Department of Forestry Bogor Indonesia. | 92 |
| Martínez-Cabrera et al. 2009. Wood anatomy and wood density in shrubs: responses to varying aridity along transcontinental transects. American Journal of Botany 96(8), 1388–1398. | 62 |
| Martínez-Sancho et al. 2020. The GenTree Dendroecological Collection, tree-ring and wood density data from seven tree species across Europe. Sci Data 7, 1. https://doi.org/10.1038/s41597-019-0340-y | 3071 |
| Martínez-Vilalta J, Cochard H, Mencuccini M, Sterck F, Herrero A, Korhonen JFJ, Llorens P, Nikinmaa E, Nolè A, Poyatos R, Ripullone F, Sass-Klaassen U and Zweifel R in press. Hydraulic adjustment of Scots pine across Europe. New Phytologist. | 12 |
| Martinez-Yrizar A., Sarukhan J., Perez-Jimenez A., Rincon E., Maass J.M., Solis-Magallanes A., Cervantes L. 1992 Above-Ground Phytomass of a Tropical Deciduous Forest on the Coast of Jalisco, Mexico. J Trop Ecol 8(1), 87-96. | 9 |
| Martins, R. 1944. Livro das Arvores do Parana. Edicao do Diretorio Regional de Geografia do Estado do Parana, Curitiba, Brasil. | 127 |
| Mayr S , Wofschwenger M, Bauer H 2002. Winter-drought induced embolism in Norway spruce Picea abies. at the alpine timberline. Physiologia Plantarum 115, 74-80; Mayr S, Hacke U, Schmid P, Schwienbacher F, Gruber A 2006. Frost drought in conifers at the alpine timberline: xylem dysfunction and adaptations. Ecology 87, 3175-3185 | 1 |
| Mayr S, Beikircher B, Obkircher M-A, Schmid P. Hydraulic plasticity and limitations of alpine Rhododendron species. Oecologia 164: 321-30. | 3 |
| Mayr S, Hacke U, Schmid P, Schwienbacher F, Gruber A 2006. Frost drought in conifers at the alpine timberline: xylem dysfunction and adaptations. Ecology 87, 3175-3185 | 7 |
| Mayr S, Schwienbacher F, Bauer H 2003. Winter at the alpine timberline: why does embolism occur in Norway spruce but not in stone pine? Plant Physiology 131, 780-792. Mayr S, Hacke U, Schmid P, Schwienbacher F, Gruber A 2006. Frost drought in conifers at the alpine timberline: xylem dysfunction and adaptations. Ecology 87, 3175-3185 | 2 |

| | |
|---|---|
| McCulloh K.A., Johnson D.M., Meinzer F.C., Voelker S.L., Lachenbruch B., Domec J.C. 2012. Hydraulic architecture of two species differing in wood density: opposing strategies in co-occurring tropical pioneer trees. Plant, Cell and Environment 35:116-25. | 2 |
| McCulloh, K. A., Meinzer, F. C., Sperry, J. S., Lachenbruch, B., Voelker, S. L., Woodruff, D. R., & Domec, J. C. (2011). Comparative hydraulic architecture of tropical tree species representing a range of successional stages and wood density. Oecologia, 167(1), 27-37. | 20 |
| McNab_and_Clark 1982, from: Radtke et al. 2016. LegacyTreeData: A national database of detailed tree measurements for volume, weight, and physical properties. In: Stanton SM, Christensen GA, (eds). Proceedings of Forest Inventory and Analysis 2015 Science Symposium, December 8–10, 2015, Portland, OR. GTR-PNW-931, USDA Forest Service, Pacific Northwest Research Station, GTR-PNW-931, Portland, OR, pp. 25–30. | 268 |
| Medeiros et al. 2019. An extensive suite of functional traits distinguishes Hawaiian wet and dry forests and enables prediction of species vital rates. Functional Ecology 33(4), 712-734. | 85 |
| Meinzer F.C., James S.A., Goldstein G., Woodruff D. 2003. Whole-tree water transport scales with sapwood capacitance in tropical forest canopy trees. Plant Cell and Environment 26: 1147-1155. | 4 |
| MEJÍA M., N.E.; UCEDA C., M. Poder calorífico de cinco espécies de Bombacaceas. Revista Florestal Del Peru, 19(1):93-97. 1992. | 5 |
| Meniado, J.A., Valbuena, R.R. and Tamolang, F.N. 1974. Timbers of the Philippines, Vol. I. Manila. | 6 |
| Miles and Smith 2009. Specific Gravity and Other Properties of Wood and Bark for 156 Tree Species Found in North America. USDA Research Note NRS-38. | 3 |
| Mirzoev 1975, Schepaschenko et al. (2017) Sci. Data 4:170070. DOI: 10.1038/sdata.2017.70 | 1 |
| Missio, F.F., Silva, A.C., Higuchi, P., Longhi, S.J., Brand, M.A., Dalla Rosa, A., ... & Pscheidt, F. (2017). ATRIBUTOS FUNCIONAIS DE ESPÉCIES ARBÓREAS EM UM FRAGMENTO DE FLORESTA OMBRÓFILA MISTA EM LAGES-SC. Ciência Florestal, 27(1), 215-224. | 20 |
| Mitchell et al. 2018. Correlated evolution between climate and suites of traits along a fast–slow continuum in the radiation of Protea. Ecol Evol 8, 1853– 1866. https://doi.org/10.1002/ece3.3773. | 775 |
| Mitchell P.J., Veneklaas E., Lambers H., Burgess S.S.O. Submitted. Soil moisture and hydraulic limitation of seasonal transpiration patterns in contrasting plant functional types along a topographical gradient ins south-western Australia. Functional Ecology. | 7 |
| Mitchell P.J., Veneklaas E.J., Lambers H. & Burgess S.S.O. (2008) Using multiple trait associations to define hydraulic functional types in plant communities of south-western Australia. Oecologia 158, 385-397. | 14 |
| Molchanov 1971 1974a; Molchanov Polyakova 1974, Schepaschenko et al. (2017) Sci. Data 4:170070. DOI: 10.1038/sdata.2017.70 | 106 |
| Molchanov 1971, Schepaschenko et al. (2017) Sci. Data 4:170070. DOI: 10.1038/sdata.2017.70 | 63 |
| Molchanov 1974b, Schepaschenko et al. (2017) Sci. Data 4:170070. DOI: 10.1038/sdata.2017.70 | 44 |
| Monteiro, R.F.R., Pimenta de Franca, O.M.V. and Sardinha, R.M.A. 1971. Essencias Florestais de Angola: Estudo das suas Madeiras II: Regiao Dos Dembos. Instituto de Investigacao Cientifica de Angola, Luanda, Angola. | 11 |
| Morales et al. 2002: Laurel forests in Tenerife, Canary Islands. I. Xylem structure in stems and petioles of Laurus azorica trees. Trees 16, 529-537. | 6 |
| Moser et al. 2019. Interaction between extreme weather events and mega-dams increases tree mortality and alters functional status of Amazonian forests. Journal of Applied Ecology 00, 1-11. | 263 |
| Moskalyuk 2015, Schepaschenko et al. (2017) Sci. Data 4:170070. DOI: 10.1038/sdata.2017.70 | 26 |
| Mroz, from: Radtke et al. 2016. LegacyTreeData: A national database of detailed tree measurements for volume, weight, and physical properties. In: Stanton SM, Christensen GA, (eds). Proceedings of Forest Inventory and Analysis 2015 Science Symposium, December 8–10, 2015, Portland, OR. GTR-PNW-931, USDA Forest Service, Pacific Northwest Research Station, GTR-PNW-931, Portland, OR, pp. 25–30. | 218 |
| Mugasha et al. 2016. Allometric Models for Estimating Tree Volume and Aboveground Biomass in Lowland Forests of Tanzania. International Journal of Forestry Research, Article ID 8076271 \| https://doi.org/10.1155/2016/8076271. | 60 |
| Mukuralinda et al. 2021. Allometric equations, wood density and partitioning of aboveground biomass in the arboretum of Ruhande, Rwanda. Trees, Forests and People 3, 100050. | 45 |
| Muller-Landau HC, unpublished data | 210 |
| Munishi, P.K.T., Maliondo, S.M., Temu, R.P.C., and Msanya, B.M. 2004. The Potential of Afromontane Rain Forests to mitigate carbon emissions in Tanzania. Journal of the Tanzania Association of Foresters. | 30 |
| Myint et al. 2017. Estimating Quantity of Water Contained in Different Timber Species from Myanmar. The Republic of the Union of Myanmar Ministry of Natural Resources and Environmental Conservation Forest Department. Leaflet No. 27. | 104 |
| NA. Unkown source. In: Ilic, J., Boland, D., McDonald, M., Downes, G. and Blakemore, P. 2000. Woody density phase 1. State of knowledge. National carbon accounting system. Technical Report 18 Australian Greenhouse Office, Canberra, Australia. | 5 |
| Nagimov et al. 2013; Usoltsev 2015, Schepaschenko et al. (2017) Sci. Data 4:170070. DOI: 10.1038/sdata.2017.70 | 44 |
| Naidu1998 1998, from: Radtke et al. 2016. LegacyTreeData: A national database of detailed tree measurements for volume, weight, and physical properties. In: Stanton SM, Christensen GA, (eds). Proceedings of Forest Inventory and Analysis 2015 Science Symposium, December 8–10, 2015, Portland, OR. GTR-PNW-931, USDA Forest Service, Pacific Northwest Research Station, GTR-PNW-931, Portland, OR, pp. 25–30. | 60 |
| Naji et al. 2021. Variations in Wood Density, Annual Ring Widths and Other Anatomical Properties of Quercus brantii Affected by Crown Dieback. Preprint. Data at: https://figshare.com/articles/dataset/Oak_Decline_Naji_xlsx/13138583/1. | 48 |
| Nam et al. 2016. Allometric Equations for Aboveground and Belowground Biomass Estimations in an Evergreen Forest in Vietnam. PLoS ONE 11(6), e0156827. https://doi.org/10.1371/journal.pone.0156827. | 45 |
| Nandika et al. 2020. Evaluation of Color Change and Biodeterioration Resistance of Gewang (Corypha utan Lamk.) Wood. Applied Sciences 10, 7501. doi: 10.3390/app10217501. Values provided for outer and inner wood, weighted by relative area. | 1 |
| Neisch 1980, from: Radtke et al. 2016. LegacyTreeData: A national database of detailed tree measurements for volume, weight, and physical properties. In: Stanton SM, Christensen GA, (eds). Proceedings of Forest Inventory and Analysis 2015 Science Symposium, December 8–10, 2015, Portland, OR. GTR-PNW-931, USDA Forest Service, Pacific Northwest Research Station, GTR-PNW-931, Portland, OR, pp. 25–30. | 28 |
| Nelson et al. 2020. The Role of Climate Niche, Geofloristic History, Habitat Preference, and Allometry on Wood Density within a California Plant Community. Forests 11(1), 105. | 4 |
| Neves et al. 2017. The roles of rainfall, soil properties, and species traits in flowering phenology along a savanna-seasonally dry tropical forest gradient. Braz. J. Bot 40, 665–679. | 68 |
| New Zealand Forest Service. 1957. Forest Trees and Timbers of New Zealand. | 8 |
| Ngoma et al. 2018: Data for developing allometric models and evaluating carbon stocks of the Zambezi Teak Forests in Zambia. Data in Brief 17, 1361-1373. | 178 |

| | |
|---|---|
| Nguyen Ngoc Chinh et al. 1996. Vietnam Forest Trees. Forest Inventory and Planning Institute Agricultural Publishing House Hanoi. | 131 |
| Nihlgard 1972, Schepaschenko et al. (2017) Sci. Data 4:170070. DOI: 10.1038/sdata.2017.70 | 2 |
| Niu et al. 2017. Divergence in strategies for coping with winter embolism among co-occurring temperate tree species: the role of positive xylem pressure, wood type and tree stature. Functional Ecology 31(8), 1550-1560. | 22 |
| Njana, M. A., Meilby, H., Eid, T., Zahabu, E. and Malimbwi, R. E. (2016). Importance of tree basic density in biomass estimation and associated uncertainties: A case of three mangrove species in Tanzania. Annals of Forest Science 73: 1073 - 1087. | 9 |
| Njiti_1982, from: Radtke et al. 2016. LegacyTreeData: A national database of detailed tree measurements for volume, weight, and physical properties. In: Stanton SM, Christensen GA, (eds). Proceedings of Forest Inventory and Analysis 2015 Science Symposium, December 8–10, 2015, Portland, OR. GTR-PNW-931, USDA Forest Service, Pacific Northwest Research Station, GTR-PNW-931, Portland, OR, pp. 25–30. | 20 |
| Nogueira, E.M., Nelson, B.W., and Fearnside, P.M. 2005. Wood density in dense forest in central Amazonia, Brazil Forest Ecology and Management 208: 261-286. | 147 |
| Nogueira, E.M., P.M. Fearnside, B.W. Nelson & M.B. França. 2007. Wood density in forests of Brazil's 'arc of deforestation': Implications for biomass and flux of carbon from land-use change in Amazonia. Forest Ecology and Management 248(3): 119-135. http://dx.doi.org/10.1016/j.foreco.2007.04.047 | 454 |
| Nunes, E.S. & Guimarães, J.B. 2012. DETERMINAÇÃO DA DENSIDADE BÁSICA DA MADEIRA DE QUALEA PARVIFLORA MART. E QUALEA GRANDIFLORA MART. (PAU-TERRA) PARA PRODUÇÃO DE CARVÃO VEGETAL. In: XXI Seminário de Iniciação Científica. | 2 |
| Nygard and Elfving 2000. Stem basic density and bark proportion of 45 woody species in young savanna coppice forests in Burkina Faso. Ann. For. Sci. 57, 143–153. | 149 |
| Nyirambangutse, B., Zibera, E., Uwizeye, F. K., Nsabimana, D., Bizuru, E., Pleijel, H., ... & Wallin, G. (2017). Carbon stocks and dynamics at different successional stages in an Afromontane tropical forest. Biogeosciences, 14(5), 1285. | 39 |
| Oey Djoen Seng. 1951. (Transl. into Indonesian by Soewarsono, P.H. 1990). Specific gravity of Indonesian Woods and Its Significance for Practical Use. FRPDC Forestry Department, Bogor, Indonesia. Indonesian Title: Berat jenis dari jenis-jenis kayu Indonesia dan pengertian beratnya kayu untuk keperluan praktek. Original Dutch title: De soortelijke gewichten van Indonesische houtsoorten en hun betekenis voor de praktijk. | 1225 |
| Oliveira 2003. Características anatômicas, químicas e térmicas da madeira de três espécies de maior ocorrência no semi-árido nordestino. UFV, Viçosa,122p. | 27 |
| Oliveras I., Martínez-Vilalta J., Jimenez-Ortiz T., Lledó M.J., Escarré A. and Piñol J. 2003. Hydraulic properties of Pinus halepensis, Pinus pinea and Tetraclinis articulata in a dune ecosystem of Eastern Spain. Plant Ecology 169: 131-141. | 6 |
| Omog Hardwood Consulting: lumber. www.omog-hardwood.com/english/speciesanduses.htm. | 4 |
| Onoda, Y., Richards, A. E., & Westoby, M. (2010). The relationship between stem biomechanics and wood density is modified by rainfall in 32 Australian woody plant species. New Phytologist, 185(2), 493-501. | 32 |
| Osborne, N. L., Høibø, Ølav. A., & Maguire, D. A. (2016). Estimating the density of coast Douglas-fir wood samples at different moisture contents using medical X-ray computed tomography. Computers and Electronics in Agriculture, 127, 50-55. | 3 |
| Osuri et al. 2014: Altered stand structure and tree allometry reduce carbon storage in evergreen forest fragments in India's Western Ghats. Forest Ecology and Management. doi:10.1016/j.foreco.2014.01.039 | 71 |
| Ovington 1957, Schepaschenko et al. (2017) Sci. Data 4:170070. DOI: 10.1038/sdata.2017.70 | 2 |
| Ovington Madgwick 1959a, Schepaschenko et al. (2017) Sci. Data 4:170070. DOI: 10.1038/sdata.2017.70 | 2 |
| own field data (Apgaua) | 72 |
| own field data (Bastin) | 475 |
| own field data (Borah) | 437 |
| own field data (Boukili) | 28 |
| own field data (Camarero) | 30 |
| own field data (Cazzolla Gatti) | 4 |
| own field data (Dar) | 66 |
| own field data (De la Riva) | 128 |
| own field data (Fajardo) | 35 |
| own field data (Flores) | 69 |
| own field data (Fyllas) | 333 |
| own field data (Hietz) | 178 |
| own field data (Homeier) | 405 |
| own field data (Jansen) | 26 |
| own field data (Jucker) | 912 |
| own field data (Kotowska) | 93 |
| own field data (Kubota) | 87 |
| own field data (Lawson) | 133 |
| own field data (Lopez) | 20 |
| own field data (Luiz Alves de Lima) | 45 |
| own field data (Macinnis-Ng) | 53 |
| own field data (Magnago) | 599 |
| own field data (Martin) | 165 |
| own field data (McCarthy) | 984 |

| | |
|---|---|
| own field data (Oliveira) | 372 |
| own field data (Olson) | 524 |
| own field data (Ramananantoandro) | 17 |
| own field data (Razafindratsima) | 53 |
| own field data (Richardson) | 716 |
| own field data (Rinkumoni) | 9 |
| own field data (Rosas) | 449 |
| own field data (Rosell) | 340 |
| own field data (Salguero-Gomez) | 210 |
| own field data (Schuldt) | 40 |
| own field data (Schwendenmann) | 33 |
| own field data (Staples) | 137 |
| own field data (Tanaka) | 223 |
| own field data (Tng) | 21 |
| own field data (Van der Sande) | 225 |
| own field data (Wright) | 3750 |
| own field data / Wannes Hubau, Tom De Mil, Mélissa Rousseau, Hans Beeckman | 4451 |
| own field data; Data also partially reported in Goodman et al. (2014) Ecological Applications 24(4): 680–698 & Goodman et al. (2013) Dryad Digital Repository. http://dx.doi.org/10.5061/dryad.p281g | 39 |
| Own field data; Matheny, A. M., S. R. Garrity and G. Bohrer (2017). The Calibration and Use of Capacitance Sensors to Monitor Stem Water Content in Trees. Journal of Visual Experiments. In Press. | 5 |
| Panshin, A.J. and de Zeeuw, C. 1980. Textbook of Wood Technology. 4th ed. McGraw-Hill, New York. | 2 |
| Parolin, P., Ferreira, L.V., and Junk, W.J. 1998. Central Amazonian floodplains: effect of two water types on the wood density of trees. Verh. Internat. Verein. Limnol. 26, 1106-1112. Data also used in Parolin, P. and M. Worbes. 2000. Wood density of trees in black water floodplains of Rio Jari National Park, Amazonia, Brazil. Acta Amazonica 30:441-448. and in Parolin, P, Ferreira LV, 1998. Are there differences in specific wood gravities between trees in varzea and igapo (Central Amazonia)? Ecotropica 4, 25-32. | 61 |
| Patino et al. 2009: Branch xylem density variations across the Amazon Basin. Biogeosciences 6, 545-568. | 1033 |
| Paul, K.I., Larmour, J., Specht, A., et al. (2019). Testing the generality of below-ground biomass allometry across plant functional types at the continent scale. Forest Ecology and Management 432, 102-114. | 80 |
| Paul, K.I., Roxburgh, S.H., Chave, J. et al. (2016). Testing the generality of above-ground biomass allometry across plant functional types at the continent scale. Global Change Biology, 22, 2106-2124. | 180 |
| Paula, J.E. & Alves, J.L.H. 2010. 922 Madeiras nativas do Brasil: anatomia, dendrologia, dendrometria, produção e uso. Ed. Cinco Continentes, Porto Alegre. 461p. | 538 |
| PAULA, J.E., IMAÑA-ENCIMAS, J.; PEREIRA, B.A.S. 1993. Inventário de um hectare de Mata Ripária. Pesquisa Agropecuária Brasileira 28(2):143-152. | 36 |
| PAULA, J.E.; IMAÑA-ENCINAS, J. & PEREIRA, B.A.S. 1996. Parámetros volumétricos e da biomassa da mata ripária do Córrego dos Macacos. Cerne 2 (2): 91-105. | 49 |
| PAULA, J.E.; IMAÑA-ENCINAS, J. & SUGIMOTO, N. 1998. Levantamento quantitativo em três hectares de vegetação de cerrado. Pesquisa Agropecuária Brasileira 33 (5): 613-620. | 22 |
| Pearson, R.S. and Brown, H.P. 1932. Commercial timbers of India. Vols. I and II. Goernment of India, Central Publication Branch, Calcutta. | 50 |
| Peichl_and_Arain 2007, from: Radtke et al. 2016. LegacyTreeData: A national database of detailed tree measurements for volume, weight, and physical properties. In: Stanton SM, Christensen GA, (eds). Proceedings of Forest Inventory and Analysis 2015 Science Symposium, December 8–10, 2015, Portland, OR. GTR-PNW-931, USDA Forest Service, Pacific Northwest Research Station, GTR-PNW-931, Portland, OR, pp. 25–30. | 20 |
| Pellinen 1986, Schepaschenko et al. (2017) Sci. Data 4:170070. DOI: 10.1038/sdata.2017.70 | 17 |
| Perala_Alban 1994, from: Radtke et al. 2016. LegacyTreeData: A national database of detailed tree measurements for volume, weight, and physical properties. In: Stanton SM, Christensen GA, (eds). Proceedings of Forest Inventory and Analysis 2015 Science Symposium, December 8–10, 2015, Portland, OR. GTR-PNW-931, USDA Forest Service, Pacific Northwest Research Station, GTR-PNW-931, Portland, OR, pp. 25–30. | 953 |
| Peschiutta M.L., Bucci S.J., Scholz F.G., Kowal R.F., Goldstein G.H. 2013. Leaf and stem hydraulic traits in relation to growth, water use and fruit yield in Prunus avium L. cultivars. Trees 27: 1559-1569. | 3 |
| Phillips_and_McNab 1982, from: Radtke et al. 2016. LegacyTreeData: A national database of detailed tree measurements for volume, weight, and physical properties. In: Stanton SM, Christensen GA, (eds). Proceedings of Forest Inventory and Analysis 2015 Science Symposium, December 8–10, 2015, Portland, OR. GTR-PNW-931, USDA Forest Service, Pacific Northwest Research Station, GTR-PNW-931, Portland, OR, pp. 25–30. | 434 |
| Phillips_and_McNab, from: Radtke et al. 2016. LegacyTreeData: A national database of detailed tree measurements for volume, weight, and physical properties. In: Stanton SM, Christensen GA, (eds). Proceedings of Forest Inventory and Analysis 2015 Science Symposium, December 8–10, 2015, Portland, OR. GTR-PNW-931, USDA Forest Service, Pacific Northwest Research Station, GTR-PNW-931, Portland, OR, pp. 25–30. | 600 |
| Pittermann J., Sperry J.S., Hacke U.G., Wheeler J.K., Sikkema E.L. 2006. Inter-tracheid pitting and the hydraulic efficiency of conifer wood: the role of tracheid allometry and cavitation protection. American Journal of Botany 93: 1265–1273; Pittermann J., Sperry J.S., Wheeler J.K., Hacke U.G., Sikkema E. 2006. Mechanical reinforcement against tracheid implosion compromises the hydraulic efficiency of conifer xylem. Plant, Cell and Environment 29: 1618-1628 | 30 |
| Plavcová, L. and Hacke, U.G. (2012). Phenotypic and developmental plasticity of xylem in hybrid poplar saplings subjected to experimental drought, nitrogen fertilization, and shading. J. Exp.Bot. 63: 6481-6491. Plavcová, L., Hacke, U.G., and Sperry, J.S. (2011). Linking irradiance-induced changes in pit membrane ultrastructure with xylem vulnerability to cavitation. Plant Cell Environ. 34: 501–513. | 11 |
| Ploton, P., Barbier, N., Momo, S. T., Réjou-Méchain, M., Boyemba Bosela, F., Chuyong, G. B., ... & Henry, M. (2016). Closing a gap in tropical forest biomass estimation: taking crown mass variation into account in pantropical allometries. Biogeosciences, 13, 1571-1585. | 77 |
| Plourde, B. T., Boukili, V. K., & Chazdon, R. L. (2015). Radial changes in wood specific gravity of tropical trees: inter-and intraspecific variation during secondary succession. Functional Ecology, 29(1), 111-120. https://doi.org/10.1111/1365-2435.12305 AND Plourde BT, Boukili V, Chazdon R (2014 Dryad Digital Repository. https://doi.org/10.5061/dryad.sv181 | 925 |

| | |
|---|---|
| PMRC_1976 1976, from: Radtke et al. 2016. LegacyTreeData: A national database of detailed tree measurements for volume, weight, and physical properties. In: Stanton SM, Christensen GA, (eds). Proceedings of Forest Inventory and Analysis 2015 Science Symposium, December 8–10, 2015, Portland, OR. GTR-PNW-931, USDA Forest Service, Pacific Northwest Research Station, GTR-PNW-931, Portland, OR, pp. 25–30. | 685 |
| PMRC_1977 1977, from: Radtke et al. 2016. LegacyTreeData: A national database of detailed tree measurements for volume, weight, and physical properties. In: Stanton SM, Christensen GA, (eds). Proceedings of Forest Inventory and Analysis 2015 Science Symposium, December 8–10, 2015, Portland, OR. GTR-PNW-931, USDA Forest Service, Pacific Northwest Research Station, GTR-PNW-931, Portland, OR, pp. 25–30. | 724 |
| PMRC_1981 1981, from: Radtke et al. 2016. LegacyTreeData: A national database of detailed tree measurements for volume, weight, and physical properties. In: Stanton SM, Christensen GA, (eds). Proceedings of Forest Inventory and Analysis 2015 Science Symposium, December 8–10, 2015, Portland, OR. GTR-PNW-931, USDA Forest Service, Pacific Northwest Research Station, GTR-PNW-931, Portland, OR, pp. 25–30. | 275 |
| PMRC_1982 1982, from: Radtke et al. 2016. LegacyTreeData: A national database of detailed tree measurements for volume, weight, and physical properties. In: Stanton SM, Christensen GA, (eds). Proceedings of Forest Inventory and Analysis 2015 Science Symposium, December 8–10, 2015, Portland, OR. GTR-PNW-931, USDA Forest Service, Pacific Northwest Research Station, GTR-PNW-931, Portland, OR, pp. 25–30. | 288 |
| Poorter, L., McNeil, A., Hurtado, V. H., Prins, H. H., & Putz, F. E. (2014). Bark traits and life-history strategies of tropical dry-and moist forest trees. Functional Ecology, 28(1), 232-242. | 89 |
| Powers and Tiffin 2010. Plant functional type classifications in tropical dry forests in Costa Rica: leaf habit versus taxonomic approaches. Functional Ecology 24(4), 927-936. Re-used and published in other publications (e.g. Werden et al. 2018, Functional Ecology). | 87 |
| Pratt R.B., Black R.A. 2006. Do invasive trees have a hydraulic advantage over native trees? Biological Invasions 8: 1331-341. | 10 |
| Pratt RB, Jacobsen AL, Golgotiu KA, Sperry JS, Ewers FW, Davis SD. 2007. Life history type coupled to water stress tolerance in nine Rhamnaceae species of the California chaparral. Ecological Monographs 77: 239-253; Pratt RB, Jacobsen AL, Ewers FW, Davis SD. 2007. Relationships among xylem transport, biomechanics, and storage in stems and roots of nine Rhamnaceae species of the California chaparral. New Phytologist 174: 787-798. | 18 |
| Pritzkow, C., Heinrich, I., Grudd, H., & Helle, G. (2014). Relationship between wood anatomy, tree-ring widths and wood density of Pinus sylvestris L. and climate at high latitudes in northern Sweden. Dendrochronologia, 32(4), 295-302. | 1 |
| Prokopovich 1995, Schepaschenko et al. (2017) Sci. Data 4:170070. DOI: 10.1038/sdata.2017.70 | 20 |
| Prospect: The Wood Database Version 2.1 to 2.3. Formerly accessible: http://www.plants.ox.ac.uk/ofi/prospect/downloads/index.htm | 272 |
| Proyectos Andinos de Desarrollo Technologico en el area de los recursos forestales tropicales. 1981. Tablas de propiedades fisicas y mecanicas de la madera de 20 especies del Peru. PADT-REFORT, Lima. By T Baker. | 48 |
| Pulgar Lorenzo and Riesco Muñoz 2018: Inter-tree and intra-tree variation in the physical properties of wood of laurel (Laurus nobilis). European Journal of Forest Research 137(4), 507-515. | 1 |
| Purkayastha, S.K. 1982. Indian Woods: Their identifications, properties and uses, Vol. IV, Myrlacene to Symploceae, Controller of Publications, New Delhi. | 3 |
| Quirino, W.F., Vale, A.T., Andrade, A.P.A., Abreu, V.L.S. & AZEVEDO, A.D.S. 2005. Poder calorífico da madeira e de materiais ligno-celulósicos. Revista da Madeira, 89, 100-106. | 5 |
| Rai & Proctor 1986: Ecological Studies on Four Rainforests in Karnataka, India: I. Environment, Structure, Floristics and Biomass. Journal of Ecology 74, 439-454. | 12 |
| Ramananantoandro et al. (2015) Forest aboveground biomass estimates in a tropical rainforest in Madagascar: new insights from the use of wood specific gravity data. Journal of Forestry Science 26(1):47–55 | 44 |
| Ramananantoandro et al. (2017) Wood specific gravity for trees in Ankeniheny Zahamena forest corridor, Madagascar. https://doi.org/10.5285/5e9fe20b-8d00-4bd3-99cb-7989fa781348 | 413 |
| Razafindratsima OH, Brown KA, Carvalho F, Johnson SE, Wright PC, Dunham AE (2017) Data from: Edge effects on components of diversity and above-ground biomass in a tropical rainforest. Dryad Digital Repository. https://doi.org/10.5061/dryad.jn743 | 132 |
| Read et al. 2011. Wood properties and trunk allometry on co-occurring rainforest canopy trees in a cyclone-prone environment. American Journal of Botany 98(11), 1762-1772. | 39 |
| Reid, Collins and Associates, 1977. Jari hog fueld study: investigation of moisture content, specific gravity, rate of drying and other related properties of indigenous hardwood species at Jari, Brazil. Progress Report, dry season sampling and results, Vancouver BC, 63 pp. In Fearnside, P.M. 1997. Wood density for estimating forest biomass in Brazilian Amazonia. Forest Ecology and Management 90: 59-87. | 13 |
| Research Group... 1964, Schepaschenko et al. (2017) Sci. Data 4:170070. DOI: 10.1038/sdata.2017.70 | 14 |
| Reyes, G., Brown, S., Chapman, J. and Lugo, A.E. 1992. Wood densities of tropical tree species. General Technical Report SO-88, United States Department of Agriculture, Forest Service, Southern Forest Experiment Station. Brown's 1997 FAO Primer is a summary of these data. | 623 |
| Ribeiro et al. 2019. Functional diversity and composition of Caatinga woody flora are negatively impacted by chronic anthropogenic disturbance. Journal of Ecology 107, 2291-2302. | 39 |
| Ribeiro, S.C., Fehrmann, L., Soares, C.P.B., Jacovine, L.A.G., Klein, C., & Gaspar, R.O. 2011. Above-and belowground biomass in a Brazilian Cerrado. Forest Ecology and Management, 262(3), 491-499. | 18 |
| Rich, P.M. 1987. Mechanical structure of the stem of arborescent palms. Botanical Gazette 148: 42-50. | 6 |
| Rijsdijk, J.F. and Laming, P.B. 1994. Physical and Related Properties of 145 Timbers. Kluwer Academic Publishers, Dordrecht, Netherlands, 380 pp. | 44 |
| Rodriguez et al. 2016: Variability in Wood Density and Wood Fibre Characterization of Woody Species and Their Possible Utility in Northeastern Mexico. American Journal of Plant Sciences 7, 1139-1150. | 37 |
| Roeder et al. 2019. Wood density, growth and mortality relationships of lianas on environmental gradients in fragmented forests of montane landscapes. Journal of Vegetation Science 00, 1-10. | 116 |
| Romero et al. 2020: Wood density, deposits and mineral inclusions of successional tropical dry forest species. European Journal of Forest Research 139, 369-381. | 21 |
| Rosner, S., Klein, A., Müller, U., & Karlsson, B. (2008). Tradeoffs between hydraulic and mechanical stress responses of mature Norway spruce trunk wood. Tree physiology, 28(8), 1179-1188. | 2 |
| Rungwattana, K., & Hietz, P. Radial variation of wood functional traits reflect size-related adaptations of tree mechanics and hydraulics. Functional Ecology. (in press). | 5 |
| Salazar C.A. 1966. Identification of Trees of Peru. Final Report Collection of wood samples and herbarium voucher specimens from the forest trees of Peru. Servicio Forestal y de Caza, Ministerio de Agricultura, Lima, Peru. 15 pp. | 16 |
| Sanaphre-Villanueva et al. 2016. Functional Diversity of Small and Large Trees Along Secondary Succession in a Tropical Dry Forest. Forests 7, 163. Obtained indirectly from: Herhández-Stefanoni et al. 2020. Improving aboveground biomass maps of tropical dry forests by integrating LiDAR, ALOS PALSAR, climate and field data. Carbon Balance and Management 15, 15. | 64 |
| Santiago, L.S., G. Goldstein, F.C. Meinzer, J.B. Fisher, K. Machado, D. Woodruff and T. Jones. 2004. Leaf photosynthetic traits scale with hydraulic conductivity and wood density in Panamanian forest canopy trees. Oecologia 140: 543-550. | 20 |
| Santini et al. 2012, DOI 10.1007/s00468-012-0729 | 19 |

| | |
|---|---|
| Santini et al. 2013, http://dx.doi.org/10.1071/FP12204 | 6 |
| Santini et al. 2016, DOI 10.1007/s00468-015-1301 | 14 |
| Santini et al. 2017, Santini NS, Cleverly J, Faux R, McBean K, Nolan R, Eamus D. Root xylem characteristics and hydraulic strategies of species co occurring in semi arid Australia. In Press IAWA Journal + FIELD | 6 |
| Sargent 1884. Report on the Forests of North America (Exclusive of Mexico). Washington, Government Printing Office. Part II. The Woods of the United States. | 425 |
| Sarman 1984, Schepaschenko et al. (2017) Sci. Data 4:170070. DOI: 10.1038/sdata.2017.70 | 1 |
| Sarmiento et al. 2011. Within-individual variation of trunk and branch xylem density in tropical trees. American Journal of Botany 98(1),140-149. | 1032 |
| Saucier_and_Boyd 1982, from: Radtke et al. 2016. LegacyTreeData: A national database of detailed tree measurements for volume, weight, and physical properties. In: Stanton SM, Christensen GA, (eds). Proceedings of Forest Inventory and Analysis 2015 Science Symposium, December 8–10, 2015, Portland, OR. GTR-PNW-931, USDA Forest Service, Pacific Northwest Research Station, GTR-PNW-931, Portland, OR, pp. 25–30. | 50 |
| Saurina Kamenetskaya 1969; Kamenetskaya 1970; Kamenetskaya et al. 1973, Schepaschenko et al. (2017) Sci. Data 4:170070. DOI: 10.1038/sdata.2017.70 | 58 |
| Savva, Y., Koubaa, A., Tremblay, F., & Bergeron, Y. (2010). Effects of radial growth, tree age, climate, and seed origin on wood density of diverse jack pine populations. Trees, 24(1), 53-65. | 21 |
| Scarascia-Mugnozza 2000, Schepaschenko et al. (2017) Sci. Data 4:170070. DOI: 10.1038/sdata.2017.70 | 1 |
| Scarascia-Mugnozza et al. 2000, Schepaschenko et al. (2017) Sci. Data 4:170070. DOI: 10.1038/sdata.2017.70 | 4 |
| Schlaegal_1975 1975, from: Radtke et al. 2016. LegacyTreeData: A national database of detailed tree measurements for volume, weight, and physical properties. In: Stanton SM, Christensen GA, (eds). Proceedings of Forest Inventory and Analysis 2015 Science Symposium, December 8–10, 2015, Portland, OR. GTR-PNW-931, USDA Forest Service, Pacific Northwest Research Station, GTR-PNW-931, Portland, OR, pp. 25–30. | 856 |
| Schlaegel_MS, from: Radtke et al. 2016. LegacyTreeData: A national database of detailed tree measurements for volume, weight, and physical properties. In: Stanton SM, Christensen GA, (eds). Proceedings of Forest Inventory and Analysis 2015 Science Symposium, December 8–10, 2015, Portland, OR. GTR-PNW-931, USDA Forest Service, Pacific Northwest Research Station, GTR-PNW-931, Portland, OR, pp. 25–30. | 1174 |
| Scholz FG, S.J. Bucci, G. Goldstein, F.C. Meinzer, A.C. Franco, A. Salazar. 2008. Plant- and Stand level variation in biophysical and physiological traits along tree density gradients in the Cerrado Brazilian Journal of Plant Physiology. 20 3.:217-232 | 18 |
| Scholz, F. G., Bucci, S.J., Goldstein, G., Meinzer, F.C., Franco, A.C. and Miralles-Wilhelm, F. 2007. Biophysical properties and functional significance of stem water storage tissues in neo-tropical savanna trees. Plant Cell and Environment 30: 236-248. | 8 |
| Scholz, F.G., Phillips, N.G., Bucci, S.J., Meinzer, F.C. and Goldstein, G. 2011. Hydraulic Capacitance: Biophysics and Functional Significance of Internal Water Sources in Relation to Tree Size. Tree Physiology 4: 341-361. | 1 |
| Schöngart, J., Arieira, J., Felfili Fortes, C., Cezarine de Arruda, E., & Nunes da Cunha, C. (2011). Age-related and stand-wise estimates of carbon stocks and sequestration in the aboveground coarse wood biomass of wetland forests in the northern Pantanal, Brazil. Biogeosciences, 8(11), 3407-3421. | 12 |
| Schoonmaker, A.L., Hacke, U.G., Landhäusser, S.M., Lieffers, V.J. & Tyree, M.T. 2010. Hydraulic acclimation to shading in boreal conifers of varying shade tolerance. 2010. Plant, Cell and Environment 33:382-393. doi: 10.1111/j.1365-3040.2009.02088.x | 8 |
| Schreiber, S.G., Hacke, U.G., Hamann, A. and Thomas, B.R. 2010. Genetic variation of hydraulic and wood anatomical traits in hybrid poplar and trembling aspen. 2010. New Phytologist 190:150-160. doi: 10.1111/j.1469-8137.2010.03594.x | 13 |
| Schuldt, B., Leuschner, C., Brock, N., & Horna, V. (2013). Changes in wood density, wood anatomy and hydraulic properties of the xylem along the root-to-shoot flow path in tropical rainforest trees. Tree physiology, 33(2), 161-174. | 15 |
| Schütt, P., Schuck, H.J., Aas, G. and Lang, U.A. 1994. Enzyklopädie der Holzgewächse. Handbuch und Atlas der Dendrologie. Ecomed, Landsberg am Lech, Germany, ISBN 3-609-72030-1. | 16 |
| Schwartz et al. 2020. Topography and Traits Modulate Tree Performance and Drought Response in a Tropical Forest. Front. For. Glob. Change 3:596256. doi: 10.3389/ffgc.2020.596256 | 46 |
| Sellin A, Tullus A, Niglas A, Õunapuu E, Karusion A, Lõhmus K (2013) Humidity-driven changes in growth rate, photosynthetic capacity, hydraulic properties and other functional traits in silver birch (Betula pendula). Ecol Res 28:523–535. doi:10.1007/s11284-013-1041-1 | 2 |
| Semechkina 1978, Schepaschenko et al. (2017) Sci. Data 4:170070. DOI: 10.1038/sdata.2017.70 | 101 |
| Serra-Maluquer et al. 2021. Impacts of recurrent dry and wet years alter long-term tree growth trajectories. J Ecol. 109, 1561-1574. https://doi.org/10.1111/1365-2745.13579. | 509 |
| Shchepashchenko 2015, Schepaschenko et al. (2017) Sci. Data 4:170070. DOI: 10.1038/sdata.2017.70 | 30 |
| Sheikh et al. 2011. Wood specific gravity of some tree species in the Garhwal Himalayas, India. For. Stud. China 13(3), 225-230. | 34 |
| Siemon G. R. and Kealley I. G. 1999. Goldfields Timber: Research Report. Research Project Steering CommitteeIn: Ilic, J., Boland, D., McDonald, M., Downes, G. and Blakemore, P. 2000. Woody density phase 1. State of knowledge. National carbon accounting system. Technical Report 18 Australian Greenhouse Office, Canberra, Australia. | 148 |
| Silman M, unpublished data | 66 |
| Silva, M.S., Santos, F.A.R., Silva, C.R.A. & Silva, L.B. 2012. Características das fibras e densidade básica da madeira de quatro espécies de Mata Atlántica (Serra da Jibóia, Elísio Medrado, Bahia, Brasil): qualificação para uso e preservação. I Simpósio sobre a Biodiversidade da Mata Atlántica (Junho de 2012, Santa Teresa, ES). | 13 |
| Silveira, P. 2008. Métodos indiretos de estimativa do conterido de biomassa e do estoque de carbono em um fragmento de floresta ombrófila densa. Tese (Doutorado). UFPR, Curitiba. | 75 |
| Sirotkin and Gruk 1980, Schepaschenko et al. (2017) Sci. Data 4:170070. DOI: 10.1038/sdata.2017.70 | 5 |
| Smirnov 1971, Schepaschenko et al. (2017) Sci. Data 4:170070. DOI: 10.1038/sdata.2017.70 | 57 |
| Soboleski et al. 2017. Variação de atributos funcionais do componente arbóreo em função de gradientes edáficos em uma floresta nebular no sul do Brasil. Rodriguésia, 68(2): 291-300. | 126 |
| Sobrado, M. A. 1996. Embolism vulnerability of an evergreen tree. Biologia Plantarum 38:297–301. | 1 |
| Soerjanegara and Lemmens, R.H.M.J. (Editors), 1994. PROSEA 5(1): Timber trees: Major commercial timbers; Lemmens, R.H.M.J., Soerjanegara, I. and Wong, W.C. (Editors), 1995. PROSEA 5(2): Timber trees: Minor commercial timbers; Sosef, M.S.M., Hong, L.T. and Prawirohatmodjo, S. (Editors), 1998. PROSEA 5(3): Timber trees: Lesser-known timbers. Hong, L.T., Lemmens, R.H.M.J., Prawirohatmodjo, S., Soerianegara, I., Sosef, M.S.M. and Wong, W.C. (Editors). CD-ROM PROSEA Timber trees. Also available at: https://uses.plantnet-project.org/en/Introduction_to_PROSEA_on_Pl@ntUse, last accessed 11/07/2019 | 688 |
| Sotomayor Castellanos et al. 2010: Características acristicas de la madera de 152 especies mexicanas. Investigación e Ingeniería de la Madera. Michoacán. Vol. 24 | 152 |
| Souza et al. 2017. Niche partitioning by functional groups of tree species in a subtropical forest. Rodriguésia 68(4). | 20 |

| | |
|---|---|
| SOUZA, A.L.; BOINA, A.; SOARES, C.P.B.; VITAL, B.R.; GASPAR, R.O. & LANA, J.M. 2012. Estrutura fitossociológica, estoques de volume, biomassa, carbono e dióxido de carbono em floresta estacional semidecidual. Revista Árvore 36 (1): 169-179.; Boina, A. 2008 Quantificação de estoques de biomassa e de carbono em floresta estacional semidecidual, Vale do Rio Doce, Minas Gerais. Dissertação (Mestrado). Universidade Federal de Viçosa, Viçosa. 89p. | 68 |
| Spalt, H.A. and Stern, W.L. 1956. Survey of African Woods. I. Tropical Woods 105: 13-38. | 1 |
| Startsev 2005, Schepaschenko et al. (2017) Sci. Data 4:170070. DOI: 10.1038/sdata.2017.70 | 21 |
| Startsev 2006, Schepaschenko et al. (2017) Sci. Data 4:170070. DOI: 10.1038/sdata.2017.70 | 10 |
| Stewart, A.M. and Kloot N. H. 1957. The Mechanical Properties of Australian, New Guinea and Other Timbers. CSIROIn: Ilic, J., Boland, D., McDonald, M., Downes, G. and Blakemore, P. 2000. Woody density phase 1. State of knowledge. National carbon accounting system. Technical Report 18 Australian Greenhouse Office, Canberra, Australia. | 3 |
| Sundarapandian, S., Mageswaran, K., Gandhi, Sanjay, and Dar, Javid Ahmad 2014. Impact of Thane Cyclone on Tree Damage in Pondicherry University Campus, Puducherry, India. Current World Environment 9, 287-300. http://dx.doi.org/10.12944/CWE.9.2.09 | 26 |
| Sungpalee et al. 2009. Intra- and interspecific variation in wood density and fine-scale spatial distribution of stand-level wood density in a northern Thai tropical montane forest. Journal of Tropical Ecology 25, 359-370. | 72 |
| Suzuki, E. 1999. Diversity in speciп¬Ѓc gravity and water content of wood among Bornean tropical rainforest trees. Ecological Research 14: 211-224. | 166 |
| Szefer et al. 2017. Determinants of litter decomposition rates in a tropical forest: functional traits, phylogeny and ecological succession. Oikos 126 (8), 1101-1111. | 57 |
| T. Ibanez, J. Chave, L. Barrabé, E. Blanchard, T. Boutreux, S. Trueba, H. Vandrot, P. Birnbaum (2017). Journal of Vegetation Science. | 231 |
| Tadaki et al. 1967, Schepaschenko et al. (2017) Sci. Data 4:170070. DOI: 10.1038/sdata.2017.70 | 7 |
| Takahashi, A. 1978. Compilation of data on the mechanical properties of foreign woods: Part 3: Africa. Research Report of Foreign Wood No. 7, Shimane University, Matsue, Japan. | 66 |
| Tamarit-Urias & Fuentes-Salinas 2003: Parámetros de humedad de 63 maderas latifoliadas mexicanas en función de su densidad básica. Revista Chapingo 9, 155-164. Itself a compilation of values from Bárcenas 1985 and Fuentes 1998. | 63 |
| ter Steege & Hammond 2001: Character convergence, diversity and disturbance in tropical rainforest in Guyana. Ecology 82: 3197-3212 | 221 |
| Terekhov Usoltsev 2008, Schepaschenko et al. (2017) Sci. Data 4:170070. DOI: 10.1038/sdata.2017.70 | 159 |
| Terekhov Usoltsev 2009, Schepaschenko et al. (2017) Sci. Data 4:170070. DOI: 10.1038/sdata.2017.70 | 1 |
| Terekhov Usoltsev 2015, Schepaschenko et al. (2017) Sci. Data 4:170070. DOI: 10.1038/sdata.2017.70 | 51 |
| Tetemke et al. 2019. Allometric Models for Predicting Aboveground Biomass of Trees in the Dry Afromontane Forests of Northern Ethiopia.Forests 10(12), 1114. https://doi.org/10.3390/f10121114 | 6 |
| Thomas and Malczewski 2007. Wood carbon content of tree species in Eastern China: Interspecific variability and the importance of the volatile fraction. Journal of Environmental Management 85, 659-662. | 14 |
| Thomas_et_al, from: Radtke et al. 2016. LegacyTreeData: A national database of detailed tree measurements for volume, weight, and physical properties. In: Stanton SM, Christensen GA, (eds). Proceedings of Forest Inventory and Analysis 2015 Science Symposium, December 8–10, 2015, Portland, OR. GTR-PNW-931, USDA Forest Service, Pacific Northwest Research Station, GTR-PNW-931, Portland, OR, pp. 25–30. | 292 |
| Tng, D. Y., Jordan, G. J., & Bowman, D. M. (2013). Plant traits demonstrate that temperate and tropical giant eucalypt forests are ecologically convergent with rainforest not savanna. PloS one, 8(12), e84378. | 129 |
| Torelli 1982: Estudio promocional de 43 especies forestales tropicales Mexicanas. Programa de Cooperación Cientifica y Técnica México-Yugoslavia. Values taken from Sotomayor Castellanos et al. 2012. | 41 |
| Tossey 1982, from: Radtke et al. 2016. LegacyTreeData: A national database of detailed tree measurements for volume, weight, and physical properties. In: Stanton SM, Christensen GA, (eds). Proceedings of Forest Inventory and Analysis 2015 Science Symposium, December 8–10, 2015, Portland, OR. GTR-PNW-931, USDA Forest Service, Pacific Northwest Research Station, GTR-PNW-931, Portland, OR, pp. 25–30. | 106 |
| Tree Talk. 2005. Woods of the World Pro CD. Springer-Verlag. | 122 |
| Treurnicht et al. 2020. Functional traits explain the Hutchinsonian niches of plant species. Global Ecology and Biogeography 29, 534-545. | 26 |
| Tue et al. 2014. Carbon storage of a tropical mangrove forest in Mui Ca Mau National Park, Vietnam | 6 |
| Tullus, A, Sellin A, Kupper P, Lutter R, Pärn L, Jasinska AK, Alber M, Kukk M, Tullus T, Tullus H Lõhmus K, Sõber A (2014) Increasing air humidity-a climate trend predicted for northern latitudes-alters the chemical composition of stemwood in silver birch and hybrid aspen. Silva Fenn 48:article id 1107. doi:10.14214/sf.1107 | 2 |
| Tyree MT, Cochard H. 1996. Summer and winter embolism in oak. Impact on water relations. Annales des Sciences Forestières 53: 173-180. | 1 |
| Uller et al. 2019: Aboveground biomass quantification and tree-level prediction models for the Brazilian subtropical Atlantic Forest, Southern Forests: a Journal of Forest Science 8 (3), 261-271, DOI: 10.2989/00306525.2019.1581498. Partial overlap with Zimmermann Oliveira et al. 2019, here keeping only individual trees for stem density, and all bark measurements. | 47 |
| Umaña et al. 2016. Interspecific functional convergence and divergence and intraspecific negative density dependence underlie the seed-to-seedling transition in tropical trees. The American Naturalist 187(1), 99-109. | 72 |
| unpublished (Cochard), provided by Jansen | 38 |
| unpublished (Pittermann), provided by Jansen | 22 |
| unpublished, own field data (Blyshchyk) | 97 |
| USFS_SE3101_50, from: Radtke et al. 2016. LegacyTreeData: A national database of detailed tree measurements for volume, weight, and physical properties. In: Stanton SM, Christensen GA, (eds). Proceedings of Forest Inventory and Analysis 2015 Science Symposium, December 8–10, 2015, Portland, OR. GTR-PNW-931, USDA Forest Service, Pacific Northwest Research Station, GTR-PNW-931, Portland, OR, pp. 25–30. | 816 |
| Usoltsev 1985, Schepaschenko et al. (2017) Sci. Data 4:170070. DOI: 10.1038/sdata.2017.70 | 58 |
| Usoltsev 1997, Schepaschenko et al. (2017) Sci. Data 4:170070. DOI: 10.1038/sdata.2017.70 | 200 |
| Usoltsev 1997; 1998, Schepaschenko et al. (2017) Sci. Data 4:170070. DOI: 10.1038/sdata.2017.70 | 263 |
| Usoltsev 1998, Schepaschenko et al. (2017) Sci. Data 4:170070. DOI: 10.1038/sdata.2017.70 | 1 |
| Usoltsev 1999, Schepaschenko et al. (2017) Sci. Data 4:170070. DOI: 10.1038/sdata.2017.70 | 1 |
| Usoltsev 2000, Schepaschenko et al. (2017) Sci. Data 4:170070. DOI: 10.1038/sdata.2017.70 | 1 |

| | |
|---|---|
| Usoltsev 2001, Schepaschenko et al. (2017) Sci. Data 4:170070. DOI: 10.1038/sdata.2017.70 | 1 |
| Usoltsev 2002, Schepaschenko et al. (2017) Sci. Data 4:170070. DOI: 10.1038/sdata.2017.70 | 1 |
| Usoltsev 2003, Schepaschenko et al. (2017) Sci. Data 4:170070. DOI: 10.1038/sdata.2017.70 | 1 |
| Usoltsev 2004, Schepaschenko et al. (2017) Sci. Data 4:170070. DOI: 10.1038/sdata.2017.70 | 1 |
| Usoltsev 2005, Schepaschenko et al. (2017) Sci. Data 4:170070. DOI: 10.1038/sdata.2017.70 | 1 |
| Usoltsev 2006, Schepaschenko et al. (2017) Sci. Data 4:170070. DOI: 10.1038/sdata.2017.70 | 1 |
| Usoltsev 2015, Schepaschenko et al. (2017) Sci. Data 4:170070. DOI: 10.1038/sdata.2017.70 | 24 |
| Usoltsev et al. 1994; Usoltsev Antropov 2001, Schepaschenko et al. (2017) Sci. Data 4:170070. DOI: 10.1038/sdata.2017.70 | 10 |
| Usoltsev et al. 2004a, Schepaschenko et al. (2017) Sci. Data 4:170070. DOI: 10.1038/sdata.2017.70 | 32 |
| Usoltsev et al. 2004b, Schepaschenko et al. (2017) Sci. Data 4:170070. DOI: 10.1038/sdata.2017.70 | 53 |
| Usoltsev et al. 2012a, Schepaschenko et al. (2017) Sci. Data 4:170070. DOI: 10.1038/sdata.2017.70 | 55 |
| Usoltsev et al. 2012a, Schepaschenko et al. (2017) Sci. Data 4:170070. DOI: 10.1038/sdata.2017.70 | 36 |
| Usoltsev et al. 2012b, Schepaschenko et al. (2017) Sci. Data 4:170070. DOI: 10.1038/sdata.2017.70 | 14 |
| Usoltsev et al. 2013, Schepaschenko et al. (2017) Sci. Data 4:170070. DOI: 10.1038/sdata.2017.70 | 26 |
| Utkin and Dylis 1966; Dylis and Nosova 1977, Schepaschenko et al. (2017) Sci. Data 4:170070. DOI: 10.1038/sdata.2017.70 | 2 |
| Utrecht xylarium / National Herbarium Netherlands (www.naturalis.nl) | 761 |
| Vale, A.T. 2000. CARACTERIZAÇÃO DA BIOMASSA LENHOSA DE UM CERRADO SENSU STRICTO DA REGIÃO DE BRASÍLIA PARA USO ENERGÉTICO. Doutorado em Agronomia (Energia na Agricultura). Universidade Estadual Paulista Jrilio de Mesquita Filho, UNESP, Brasil.; VALE, A.T.; BRASIL, M.A.M.; LEÃO, A.L. 2002. Quantificação e caracterização energética da madeira e casca de espécies do cerrado. Ciência Florestal, 12(1), p. 71-80.; Vale, A.T. & Felfili, J.M. 2005. Dry biomass distribution in a cerrado sensu stricto site in Brazil central. Revista Árvore, 29(5), 661-669. | 94 |
| Vale, A.T., Dias, Í.S. & Santana, M.A.E. 2010. Relações entre propriedades químicas, físicas e energéticas da madeira de cinco espécies de cerrado. Ciência Florestal, 20(1), 137-145. | 5 |
| Values compiled from various sources by the BDFFP project | 79 |
| Van der Slooten, H.J., Richter, H.G., Aune, J.E. and Cordero, L.L.. 1971. Inventariacion y demonstracionces forestales Panama: Propriedades y usos de ciento trece especies maderables de Panama. Panama, UNFAO: SF/PAN 6. | 9 |
| van Gelder, H.A., Poorter, L. and Sterck, F.J. 2006. Wood mechanics, allometry, and life-history variation in a tropical rain forest tree community. New Phytologist 171(2): 367-378. | 30 |
| Various sources. In Fearnside, P.M. 1997. Wood density for estimating forest biomass in Brazilian Amazonia. Forest Ecology and Management 90: 59-87. | 25 |
| Venezuela database. By H ter Steege. | 82 |
| Vieilledent et al. 2012 [doi: 10.1890/11-0039.1] | 1878 |
| Villagra M., Campanello P.I., Bucci S.J., Goldstein G. 2013. Functional relationships between leaf hydraulics and leaf economic traits in response to nutrient addition in subtropical tree species. Tree Physiology 33:1308-1318. Villagra M., Campanello P.I., Montti L., Goldstein G. 2013. Removal of nutrient limitations in forest gaps enhances growth rate and resistance to cavitation in subtropical canopy tree species differing in shade tolerance. Tree Physiology 33: 285-296. | 5 |
| Vink, A.T. 1983. Surinam Timbers. State Forest Industries, Paramaribo, Suriname | 57 |
| Vins Sika 1977, Schepaschenko et al. (2017) Sci. Data 4:170070. DOI: 10.1038/sdata.2017.70 | 1 |
| von Maydell, H.J. 1983. Arbres et Arbustes du Sahel: Leurs caracteristiques et leurs utilisations. Deutsche Gesellschaft fur Technische Zusammenarbeit (GTZ) GmbH, Eschborn, Germany. | 10 |
| Vyskot 1976, Schepaschenko et al. (2017) Sci. Data 4:170070. DOI: 10.1038/sdata.2017.70 | 14 |
| Vyskot 1981, Schepaschenko et al. (2017) Sci. Data 4:170070. DOI: 10.1038/sdata.2017.70 | 37 |
| Vyskot 1982, Schepaschenko et al. (2017) Sci. Data 4:170070. DOI: 10.1038/sdata.2017.70 | 14 |
| Vyskot 1983, Schepaschenko et al. (2017) Sci. Data 4:170070. DOI: 10.1038/sdata.2017.70 | 15 |
| Vyskot 1989ab, Schepaschenko et al. (2017) Sci. Data 4:170070. DOI: 10.1038/sdata.2017.70 | 3 |
| Vyskot 1990, Schepaschenko et al. (2017) Sci. Data 4:170070. DOI: 10.1038/sdata.2017.70 | 5 |
| Vyskot 1991, Schepaschenko et al. (2017) Sci. Data 4:170070. DOI: 10.1038/sdata.2017.70 | 5 |
| Wagenführ R. 2007. Holzatlas. Leipzig, Germany: Carl Hanser Verlag. | 181 |
| Wangaard, F.F. and Muschler, A.F. 1952. Tropical Woods 98: 1-190. | 2 |
| Wangaard, F.F., Koehler, A. and Muschler, A.F. 1954. Tropical Woods 99: 1-187. | 1 |
| WDS_Pitch_Pine, from: Radtke et al. 2016. LegacyTreeData: A national database of detailed tree measurements for volume, weight, and physical properties. In: Stanton SM, Christensen GA, (eds). Proceedings of Forest Inventory and Analysis 2015 Science Symposium, December 8–10, 2015, Portland, OR. GTR-PNW-931, USDA Forest Service, Pacific Northwest Research Station, GTR-PNW-931, Portland, OR, pp. 25–30. | 200 |
| WDS_Pond_Pine, from: Radtke et al. 2016. LegacyTreeData: A national database of detailed tree measurements for volume, weight, and physical properties. In: Stanton SM, Christensen GA, (eds). Proceedings of Forest Inventory and Analysis 2015 Science Symposium, December 8–10, 2015, Portland, OR. GTR-PNW-931, USDA Forest Service, Pacific Northwest Research Station, GTR-PNW-931, Portland, OR, pp. 25–30. | 200 |
| WDS_Sand_Pine, from: Radtke et al. 2016. LegacyTreeData: A national database of detailed tree measurements for volume, weight, and physical properties. In: Stanton SM, Christensen GA, (eds). Proceedings of Forest Inventory and Analysis 2015 Science Symposium, December 8–10, 2015, Portland, OR. GTR-PNW-931, USDA Forest Service, Pacific Northwest Research Station, GTR-PNW-931, Portland, OR, pp. 25–30. | 276 |
| WDS_SFS_Pine, from: Radtke et al. 2016. LegacyTreeData: A national database of detailed tree measurements for volume, weight, and physical properties. In: Stanton SM, Christensen GA, (eds). Proceedings of Forest Inventory and Analysis 2015 Science Symposium, December 8–10, 2015, Portland, OR. GTR-PNW-931, USDA Forest Service, Pacific Northwest Research Station, GTR-PNW-931, Portland, OR, pp. 25–30. | 80 |

| | |
|---|---|
| WDS_Spruce_Pine, from: Radtke et al. 2016. LegacyTreeData: A national database of detailed tree measurements for volume, weight, and physical properties. In: Stanton SM, Christensen GA, (eds). Proceedings of Forest Inventory and Analysis 2015 Science Symposium, December 8–10, 2015, Portland, OR. GTR-PNW-931, USDA Forest Service, Pacific Northwest Research Station, GTR-PNW-931, Portland, OR, pp. 25–30. | 148 |
| WDS_Table_mountain_Pine, from: Radtke et al. 2016. LegacyTreeData: A national database of detailed tree measurements for volume, weight, and physical properties. In: Stanton SM, Christensen GA, (eds). Proceedings of Forest Inventory and Analysis 2015 Science Symposium, December 8–10, 2015, Portland, OR. GTR-PNW-931, USDA Forest Service, Pacific Northwest Research Station, GTR-PNW-931, Portland, OR, pp. 25–30. | 200 |
| WDS_Virginia_Pine, from: Radtke et al. 2016. LegacyTreeData: A national database of detailed tree measurements for volume, weight, and physical properties. In: Stanton SM, Christensen GA, (eds). Proceedings of Forest Inventory and Analysis 2015 Science Symposium, December 8–10, 2015, Portland, OR. GTR-PNW-931, USDA Forest Service, Pacific Northwest Research Station, GTR-PNW-931, Portland, OR, pp. 25–30. | 150 |
| Wells, J. unpublished data | 48 |
| Whittaker 9999, from: Radtke et al. 2016. LegacyTreeData: A national database of detailed tree measurements for volume, weight, and physical properties. In: Stanton SM, Christensen GA, (eds). Proceedings of Forest Inventory and Analysis 2015 Science Symposium, December 8–10, 2015, Portland, OR. GTR-PNW-931, USDA Forest Service, Pacific Northwest Research Station, GTR-PNW-931, Portland, OR, pp. 25–30. | 30 |
| Whittaker_et_al 1974, from: Radtke et al. 2016. LegacyTreeData: A national database of detailed tree measurements for volume, weight, and physical properties. In: Stanton SM, Christensen GA, (eds). Proceedings of Forest Inventory and Analysis 2015 Science Symposium, December 8–10, 2015, Portland, OR. GTR-PNW-931, USDA Forest Service, Pacific Northwest Research Station, GTR-PNW-931, Portland, OR, pp. 25–30. | 186 |
| Whittaker_Niering 1975, from: Radtke et al. 2016. LegacyTreeData: A national database of detailed tree measurements for volume, weight, and physical properties. In: Stanton SM, Christensen GA, (eds). Proceedings of Forest Inventory and Analysis 2015 Science Symposium, December 8–10, 2015, Portland, OR. GTR-PNW-931, USDA Forest Service, Pacific Northwest Research Station, GTR-PNW-931, Portland, OR, pp. 25–30. | 60 |
| Whittaker_Woodwell 1968, from: Radtke et al. 2016. LegacyTreeData: A national database of detailed tree measurements for volume, weight, and physical properties. In: Stanton SM, Christensen GA, (eds). Proceedings of Forest Inventory and Analysis 2015 Science Symposium, December 8–10, 2015, Portland, OR. GTR-PNW-931, USDA Forest Service, Pacific Northwest Research Station, GTR-PNW-931, Portland, OR, pp. 25–30. | 94 |
| Wiemann, M.C. and Williamson, G.B. 1989. Wood specific gravity gradients in tropical dry and montane rain forest trees. American Journal of Botany 76(6): 924-928; Wiemann, M.C., and Williamson, , G.B. 1989. Radial gradients in the specific gravity of wood in some tropical and temperate trees. Forest Science 35:197-210; MacDonald, S.S., Williamson, G.B., and Wiemann, M.C. 1995. Wood specific gravity and anatomy in Heliocarpus appendiculatus (Tiliaceae). American Journal of Botany 82, 855-861.; Perez Cordero, L.D., and Kanninen, M. 2000. Wood specific gravity and aboveground biomass of Bombacopsis quinata plantations in Costa Rica. Forest Ecology Management 165: 1-9. | 43 |
| Wigley et al. 2019. Ants, fire, and bark traits affect how African savanna trees recover following damage. 51(5), 682-691. | 10 |
| Williams et al. 2008. Carbon sequestration and biodiversity of re-growing miombo woodlands in Mozambique. Forest Ecology and Management 254, 145–155. | 26 |
| Willson CJ, PS Manos, RB Jackson. 2008. Hydraulic traits are influenced by phylogenetic history in the drought-resistant and invasive genus Juniperus (Cupressaceae). American Journal of Botany 95:299-314. | 28 |
| Willson, CJ, RB Jackson. 2006. Xylem cavitation caused by drought and freezing stress in four co-occurring Juniperus species. Physiologia Plantarum 127: 374-382. | 4 |
| Wirth et al. 1999 2002, Schepaschenko et al. (2017) Sci. Data 4:170070. DOI: 10.1038/sdata.2017.70 | 47 |
| Wittmann et al. 2008. Tree Species Composition, Structure, and Aboveground Wood Biomass of a Riparian Forest of the Lower Miranda River, Southern Pantanal, Brazil. Folia Geobotanica 43, 397. | 15 |
| Wood Industry Research Institute, Chinese Forestry Academy. 1982. Physical and Mechanical Properties of Wood from China's Important Trees. Beijing, China. 154 pp. | 16 |
| Woodcock, D.W. 2000. Wood specific gravity of trees and forest types in the Southern Peruvian Amazon. Acta Amazonica 30(4): 589-599. | 58 |
| Woodrum C.L., Ewers F.W., Telewski F.W. 2003. Hydraulic, biomechanical and anatomical study of the xylem of five species of Acer (Aceraceae). American Journal of Botany 90: 693-699. | 5 |
| WQC, from: Radtke et al. 2016. LegacyTreeData: A national database of detailed tree measurements for volume, weight, and physical properties. In: Stanton SM, Christensen GA, (eds). Proceedings of Forest Inventory and Analysis 2015 Science Symposium, December 8–10, 2015, Portland, OR. GTR-PNW-931, USDA Forest Service, Pacific Northwest Research Station, GTR-PNW-931, Portland, OR, pp. 25–30. | 814 |
| Wyatt-Smith, J. 1963. Manual of Malayan Silviculture for Inland Forest. Malayan Forest Record No. 23. Forest Research Institute, Kepong, pp. 152-167. | 8 |
| Xiao 1990, Schepaschenko et al. (2017) Sci. Data 4:170070. DOI: 10.1038/sdata.2017.70 | 6 |
| Xiao et al. 2021. Divergence of stem biomechanics and hydraulics between Bauhinia lianas and trees, AoB PLANTS, plab016, https://doi.org/10.1093/aobpla/plab016. | 120 |
| Xu et al. 1988, Schepaschenko et al. (2017) Sci. Data 4:170070. DOI: 10.1038/sdata.2017.70 | 3 |
| Yang et al. 1995, Schepaschenko et al. (2017) Sci. Data 4:170070. DOI: 10.1038/sdata.2017.70 | 7 |
| Yeboah et al. 2014. Variation in wood density and carbon content of tropical plantation tree species from Ghana. New Forests 45, 35–52, DOI 10.1007/s11056-013-9390-8. | 21 |
| Zhang YJ, Meinzer FC, Hao GY, Scholz FG, Bucci SJ, Takahashi FS, Villalobos-Vega R, Giraldo JP, Cao KF, Hoffmann WA, Goldstein G. 2009. Size-dependent mortality in a Neotropical savanna tree: the role of height-related adjustments in hydraulic architecture and carbon allocation. Plant Cell and Environment DOI: 10.1111/j.1365-3040.2009.02012.x | 4 |
| Zhang, Shi-Bao, Slik, J.W. Ferry, Zhang, Jiao-Lin, and Kun-Fang Cao 2011. Spatial patterns of wood traits in China are controlled by phylogeny and the environment. Global Ecology and Biogeography 20, 241-250. | 210 |
| Zhu et al. 1993, Schepaschenko et al. (2017) Sci. Data 4:170070. DOI: 10.1038/sdata.2017.70 | 1 |
| Zhu, S-D. & Cao, K-F. 2009. Hydraulic properties and photosynthetic rates in co-occuring lianes and trees in a seasonal tropical rainforest in southwestern China. Plant Ecol 204:295-304. DOI: 10.1007/s11258-009-9592-5 | 6 |
| Zhu, S-D., Song, J-j., Li, R-H. & Ye, Q. 2012 Plant hydraulics and photosynthesis of 34 woody species from different successional stages of subtropical forests. Plant, Cell and Environment 36: 879-891. | 34 |
| Ziemińska K, Westoby M, Wright IJ. (2015) Broad Anatomical Variation within a Narrow Wood Density Range-A Study of Twig Wood across 69 Australian Angiosperms. PLoS ONE 10: e0124892 | 69 |
| Zimmermann Oliveira et al. 2019. Towards the Fulfillment of a Knowledge Gap: Wood Densities for Species of the Subtropical Atlantic Forest. Data 2019, 4, 104. doi:10.3390/data4030104. Partial overlap with Uller et al. 2019, but no bark density data. The Zimmermann Oliveira et al. 2019 data set has been kept for stem density, except for individual trees that were identifiable as equivalent in both and thus allow for bark/stem density comparisons. | 160 |
| Zorger et al. 2019. Functional organization of woody plant assemblages along precipitation and human disturbance gradients in a seasonally dry tropical forest. Biotropica. doi:10.1111/btp.12721 | 1008 |
| Zotz, G., Tyree M.T. & Patino S. 1997. Hydraulic architecture and water relations of a flood-tolerant tropical tree, Annona glabra. Tree Physiology 17, 359-365. | 2 |

| | |
|---|---|
| Zou et al. 1995, Schepaschenko et al. (2017) Sci. Data 4:170070. DOI: 10.1038/sdata.2017.70 | 1 |

#### B) Supplementary Tables

| # | Field | Description |
|---|---|---|
| 1 | *id* | overall primary id of the record |
| 2 | *species* | species after spell-checking and resolution (WFO) |
| 3 | *group* | group after spell-checking and resolution (WFO) |
| 4 | *order* | order after spell-checking and resolution (WFO) |
| 5 | *family* | family after spell-checking and resolution (WFO) |
| 6 | *genus* | genus after spell-checking and resolution (WFO) |
| 7 | *epithet* | species epithet after spell-checking and resolution (WFO) |
| 8 | *epithet_infraspecific* | infraspecific epithet after spell-checking and resolution (WFO) |
| 9 | *authority* | taxonomic authority after spell-checking and resolution (WFO) |
| 10 | *status_taxonomic* | taxonomic status after spell-checking and resolution (WFO): "accepted", "unresolved", “unchecked” |
| 11 | *rank_taxonomic* | taxonomic rank after spell-checking and resolution (WFO): "genus", "species", "subspecies", "variety", "hybrid" |
| 12 | *source_taxonomic* | identifier after spell-checking and resolution (WFO) |
| 13 | *wsg* | wood specific gravity / basic wood density |
| 14 | *plant_agg* | is measurement aggregated over several individuals: 0 or 1, or NA if unclear |
| 15 | *plants_sampled* | number of individuals aggregated over: > 1, if *plant_agg* == 1, else 1 or NA |
| 16 | *location_sample* | the within-tree location of the sample: "root","bole","branch","twig", "shoot”, NA |
| 17 | *type_tissue* | the type of woody tissue sampled: "sapwood","heartwood","bark", "total (bark to pith)" |
| 18 | *region* | one of nine regions: "South America", "Central America and West Indies", "North America", "Asia", "South-East Asia", "Africa", "Indian Ocean", "Europe", "Oceania" |
| 19 | *country* | country of measurement |
| 20 | *site* | site of measurement, at various levels of precision |
| 21-22 | *longitude/latitude* | coordinates of the site of measurement |
| 23 | *type_forest* | type of forest, can contain locally specific types |
| 24-26 | *id_plant/age/dbh* | Individual plant-level information on IDs, age and diameter |
| 27 | *experiment* | is the study an experiment? if collected during an experiment 1, else 0 |
| 28 | *experiment_design* | design of the study or experiment if applicable |
| 29 | *source_short* | the short name of the source of measurements |
| 30 | *source_long* | the full name of the source of measurements |
| 31 | *contributor* | the contributor of the data set |
| 32 | *id_dboriginal* | the id of the sample in the original database |
| 33 | *species_reference* | the taxon supplied to the GWDD v.2 |
| 34 | *genus_reference* | the genus supplied to the GWDD v.2 |
| 35 | *epithet_reference* | the epithet supplied to the GWDD v.2 |
| 36 | *epithet_infraspecific_ reference* | the infraspecific epithet supplied to the GWDD v.2 |
| 37 | *species_reference_ canonical* | the taxon supplied to the GWDD v.2., stripped of additional information such as taxonomic authorities |
| 38 | *value_reference* | the (untransformed) wood density supplied to the GWDD v.2 |
| 39 | *quantity_reference* | the type of wood density that was measured: "Airdry SG/Density", "Ovendry SG/Density", "Basic SG/Density" |
| 40 | *moisture_airdry* | the moisture at which wood was considered airdry (%): 8, 12, 15 if airdry, else NA |
| 41 | *wsg_conversion* | the conversion factor to convert airdry or ovendry densities: 1.0 (Basic SG), 0.8676046 (Ovendry SG), 0.8404015 (8% SG), 0.8281316 (12% SG), 0.8194401 (15% SG) |
| 42 | *backtransformed* | is the value backconverted from the GWDD v.1? 0 or 1 (only applied to legacy data) |
| 43 | *type_sample* | the type of sample: "core", "disk" |
| 44 | *instrument* | the instrument used to obtain wood density |
| 45 | *temperature_drying* | the temperature at which samples have been dried |

**Table S1: Documentation of GWDD v.2 columns.** Given is the column ID, their name and a short description. WorldFloraOnline, which was used for taxonomic resolution, is abbreviated as WFO.

| *Binomial* | *Resolved* | *N* |
|---|---|---|
| Acacia alleniana | Acacia alleniana | 1 |
| Acacia colei | Acacia colei | 2 |
| Acacia trinervata | Acacia trinervata | 1 |
| Acioa edulis | Acioa edulis | 1 |
| Aglaia hiernii | Aglaia hiernii | 1 |
| Albizia dinklagei | Albizia dinklagei | 1 |
| Albizia lebbekoides | Albizia lebbekoides | 1 |
| Alloxylon flammeum | Alloxylon flammeum | 2 |
| Anacardium tenuifolium | Anacardium tenuifolium | 1 |
| Aspidosperma decussatum | Aspidosperma decussatum | 1 |
| Atalaya australiana | Atalaya australiana | 1 |
| Borassus flabellifer | Borassus flabellifer | 1 |
| Breonia madagascariensis | Breonia madagascariensis | 1 |
| Bruguiera exaristata | Bruguiera exaristata | 1 |
| Calliandra calothyrsus | Calliandra houstoniana var. calothyrsus | 1 |
| Calliandra tweediei | Calliandra tweediei | 1 |
| Corymbia dallachiana | Corymbia dallachiana | 4 |
| Couepia robusta | Couepia robusta | 1 |
| Cullen australasicum | Cullen australasicum | **1** |
| Dicymbe corymbosa | Dicymbe corymbosa | 1 |
| Diospyros cayennensis | Diospyros cayennensis | 1 |
| Dipteryx ferrea | Dipteryx ferrea | 1 |
| Endiandra dielsiana | Endiandra dielsiana | 1 |
| Eucalyptus glomerosa | Eucalyptus glomerosa | 1 |
| Eucalyptus urophylla | Eucalyptus urophylla | 2 |
| Grewia crenata | Grewia prunifolia | 1 |
| Huertea cubensis | Huertea cubensis | 1 |
| Lacistema grandifolium | Lacistema grandifolium | 1 |
| Leucaena diversifolia | Leucaena diversifolia | 1 |
| Licania fanshawei | Licania fanshawei | 1 |
| Licania jimenezii | Licania jimenezii | 1 |
| Licania pallida | Licania pallida | 1 |
| Licania sparsipilis | Leptobalanus sparsipilis | 2 |
| Licaria rigida | Licaria rigida | 2 |
| Litsea breviumbellata | Litsea breviumbellata | 1 |
| Magnolia yoroconte | Magnolia yoroconte | 1 |
| Micrandra minor | Micrandra minor | 1 |
| Mischocarpus stipitatus | Mischocarpus stipitatus | 3 |
| Mouriri pseudogeminata | Mouriri pseudogeminata | 1 |
| Nectandra krugii | Nectandra krugii | 1 |
| Parashorea aptera | Parashorea aptera | 1 |
| Pinus hartwegii | Pinus hartwegii | 1 |
| Pittosporum angustifolium | Pittosporum angustifolium | 2 |
| Polyalthia asteriella | Monoon asteriellum | 1 |
| Pouteria izabalensis | Pouteria izabalensis | 1 |
| Pouteria oblanceolata | Pouteria oblanceolata | 1 |
| Pouteria obscura | Pouteria obscura | 1 |
| Pterocarpus osun | Pterocarpus osun | 1 |
| Rauvolfia paraensis | Rauvolfia paraensis | 1 |
| Rhodamnia glauca | Rhodamnia glauca | 1 |

**Table S2: Removed taxa in GWDD v.2.** Shown are taxa from the first GWDD whose references could not be verified and whose records are not available anymore in the GWDD v.2. We also provide their resolved names from World Flora Online (WFO) and their number of entries in the first GWDD. With one exception (Cullen australicum), the removal of the taxa did not result in the loss of the corresponding genus.

| Model | Question | Data subset | Subsetting procedure | Model formula |
|---|---|---|---|---|
| **M1** | **Extent of intraspecific variation** | n = 79,488, $n_{species}$ = 10,780 | full set of measurements from individual plants | wd ~ 1 + (1 \| family / genus / species) + (1 \| source) |
| **M2** | | n = 79,488, $n_{species}$ = 10,780 | full set of measurements from individual plants | wd ~ 1 + (1 \| family / genus / species) |
| **M3** | | n = 49,991, $n_{species}$ = 2735 | higher quality (>= 3 individuals per species, >= 3 species per genus, >= 3 genera per family) | wd ~ 1 + (1 \| family / genus / species) + (1 \| source) |
| **M4** | | n = 49,991, $n_{species}$ = 2735 | higher quality (>= 3 individuals per species, >= 3 species per genus, >= 3 genera per family) | wd ~ 1 + (1 \| family / genus / species) |
| **M5** | **Partitioning of intraspecific variance** | n = 19,246, $n_{species}$ = 147 | minimum quality (>= 2 sites per species, >= 2 individuals for one site, >= 2 records for one individual) | wd ~ 1 + (1 \| species / site / id_plant) + (1 \| source) |
| **M6** | | n = 19,246, $n_{species}$ = 147 | minimum quality (>= 2 sites per species, >= 2 individuals for one site, >= 2 records for one individual) | wd ~ 1 + (1 \| species / site / id_plant) |
| **M7** | | n = 2,494, $n_{species}$ = 35 | higher quality (>= 3 sites per species, >= 3 individuals for one site, >= 3 records for one individual) | wd ~ 1 + (1 \| species / site / id_plant) + (1 \| source) |
| **M8** | | n = 2,494, $n_{species}$ = 35 | higher quality (>= 3 sites per species, >= 3 individuals for one site, >= 3 records for one individual) | wd ~ 1 + (1 \| species / site / id_plant) |
| **M9** | **Within-plant gradients** | n = 679, $n_{species}$ = 150 | species with >= 1 measurement each from heartwood and sapwood | wd ~ 1 + sapwood + (1 + sapwood \| species) + (1 \| source_short) |
| **M10** | | n = 48,494, $n_{species}$ = 2,018 | species with >= 1 measurement each from branch and trunk | wd ~ 1 + branch + (1 + branch \| species) + (1 \| source_short) |
| **M11** | | $n_{species}$ = 150 | species with >= 1 measurement each from heartwood and sapwood; species means | $wd_{sapwood}$ ~ $wd_{heartwood}$ (MA regression) |
| **M12** | | $n_{species}$ = 2,018 | species with >= 1 measurement each from branch and trunk; species means | $wd_{branch}$ ~ $wd_{trunk}$ (MA regression) |
| **M13** | | $n_{species}$ = 523 | species with >= 1 measurement each from branch sapwood and trunk sapwood; species means | $wd_{branch}$ ~ $wd_{trunk}$ (MA regression) |
| **M14** | | $n_{species}$ = 189 | species with 5 randomly drawn measurements from both branch and trunk; species means | $wd_{branch}$ ~ $wd_{trunk}$ (MA regression) |
| **M15** | | n = 3,527, $n_{species}$ = 145 | Individual plants with 1 randomly drawn measurement from both branch and trunk | $wd_{branch}$ ~ $wd_{trunk}$ (MA regression) |
| **M16** | **Environmental predictors of intraspecific variation** | n = 41,893, $n_{species}$ = 2,160 | species with >= 2 distinct locations | wd ~ 1 + $env1km_{species}$ + $env1km_{intrasp}$ + (1 + $env1km_{intrasp}$ \| species) + (1 \| source_short) |
| **M17** | | n = 41,893, $n_{species}$ = 2,160 | species with >= 2 distinct locations | wd ~ 1 + $env5km_{species}$ + $env5km_{intrasp}$ + (1 + $env5km_{intrasp}$ \| species) + (1 \| source_short) |
| **M18** | | n = 30,128, $n_{species}$ = 692 | species with >= 2 distinct locations + wide intraspecific predictor range (one p. in top 10% of ranges) | wd ~ 1 + $env1km_{species}$ + $env1km_{intrasp}$ + (1 + $env1km_{intrasp}$ \| species) + (1 \| source_short) |
| **M19** | | n = 30,128, $n_{species}$ = 692 | species with >= 2 distinct locations + wide intraspecific predictor range (one p. in top 10% of ranges) | wd ~ 1 + $env5km_{species}$ + $env5km_{intrasp}$ + (1 + $env5km_{intrasp}$ \| species) + (1 \| source_short) |
| **M20** | **Environmental predictors (tropical / extratropical)** | n = 8,783, $n_{species}$ = 700 | species with >= 3 distinct locations in the tropics | wd ~ 1 + $env1km_{species}$ + $env1km_{intrasp}$ + (1 + $env1km_{intrasp}$ \| species) + (1 \| source_short) |
| **M21** | | n = 8,783, $n_{species}$ = 700 | species with >= 3 distinct locations in the tropics | wd ~ 1 + $env5km_{species}$ + $env5km_{intrasp}$ + (1 + $env5km_{intrasp}$ \| species) + (1 \| source_short) |
| **M22** | | n = 26,437, $n_{species}$ = 247 | species with >= 3 distinct locations outside the tropics | wd ~ 1 + $env1km_{species}$ + $env1km_{intrasp}$ + (1 + $env1km_{intrasp}$ \| species) + (1 \| source_short) |
| **M23** | | n = 26,437, $n_{species}$ = 247 | species with >= 3 distinct locations outside the tropics | wd ~ 1 + $env5km_{species}$ + $env5km_{intrasp}$ + (1 + $env5km_{intrasp}$ \| species) + (1 \| source_short) |
| **M24** | **Environmental predictors (gymnosperms)** | n = 12,089, $n_{species}$ = 59 | species with >= 2 distinct locations; gymnosperms only | wd ~ 1 + $env1km_{species}$ + $env1km_{intrasp}$ + (1 + $env1km_{intrasp}$ \| species) + (1 \| source_short) |
| **M25** | | n = 12,089, $n_{species}$ = 59 | species with >= 2 distinct locations; gymnosperms only | wd ~ 1 + $env5km_{species}$ + $env5km_{intrasp}$ + (1 + $env5km_{intrasp}$ \| species) + (1 \| source_short) |
| **M26** | **Predictivity** | variable | full GWDD v.2, but tested on $n_{species}$ = 1,667 (records from >= 5 sources) | wd ~ 1 + (1 \| family / genus / species) + (1 \| source / site) |
| **M27** | | variable | full GWDD v.2, but tested on $n_{species}$ = 318 (species with >= 3 sites, one site with >= 4 records) | wd ~ 1 + branch + (1 + branch \| family / genus / species) + (1 \| source / site) |

**Table S3: Overview over models and data subsets.** This is a quick reference for combinations of data subsets and models used in this study, the questions they address, and the subsetting procedure. Model formulas are provided in the mixed-effects model notation employed by the R packages *lme4* (Bates et al., 2015) and *brms* (Bürkner, 2018) and fitted with both approaches. In the latter, we explicitly model the distributional parameter $\sigma^* = \exp(\sigma)$ as $\sigma^*$ ~ 1 + (1 | species) or $\sigma^*$ ~ 1 + (1 | family / genus / species), in line with random effects used for the response variable *wd*. The predictor *env1km* is a shorthand for six predictors from CHELSA (Brun et al., 2022; Karger et al., 2017) and soilgrids at 1km resolution (Hengl et al., 2017), *env5km* for the equivalents from TerraClimate (Abatzoglou et al., 2018) and soilgrids (5 km). All environmental and edaphic predictors are split into species means (subscript “species”) and within-species deviations from species means (subscript “intrasp”).

| *brms/STAN* | **M1: Full dataset (incl. source effect)** | | **M2: Full dataset (no source effect)** | | **M3: High quality subset (incl. source effect)** | | **M4: High quality subset (no source effect)** | |
|---|---|---|---|---|---|---|---|---|
| **Wood density (g cm$^{-3}$)** | *Estimate* | *CI* | *Estimate* | *CI* | *Estimate* | *CI* | *Estimate* | *CI* |
| (Intercept) | 0.557 | [0.540; 0.575] | 0.554 | [0.538; 0.569] | 0.579 | [0.554; 0.605] | 0.575 | [0.552; 0.599] |
| **Random effects** | *Estimate* | *(%Var)* | *Estimate* | *(%Var)* | *Estimate* | *(%Var)* | *Estimate* | *(%Var)* |
| family | 0.098 | (30%) | 0.098 | (32%) | 0.067 | (17%) | 0.065 | (17%) |
| family:genus | 0.097 | (30%) | 0.098 | (32%) | 0.092 | (33%) | 0.094 | (36%) |
| family:genus:species | 0.074 | (17%) | 0.077 | (20%) | 0.077 | (23%) | 0.079 | (26%) |
| source | 0.049 | (8%) | | | 0.049 | (9%) | | |
| residual (σ) | 0.068 | (15%) | 0.071 | (17%) | 0.068 | (18%) | 0.071 | (21%) |
| **log(**σ) | *Estimate* | *CI* | *Estimate* | *CI* | *Estimate* | *CI* | *Estimate* | *CI* |
| (Intercept) | -2.796 | [-2.830; -2.764] | -2.786 | [-2.818; -2.753] | -2.740 | [-2.794; -2.686] | -2.727 | [-2.780; -2.679] |
| **Random effects** | *Estimate* | *(%Var)* | *Estimate* | *(%Var)* | *Estimate* | *(%Var)* | *Estimate* | *(%Var)* |
| family | 0.128 | (9%) | 0.122 | (8%) | 0.119 | (8%) | 0.114 | (7%) |
| family:genus | 0.173 | (16%) | 0.167 | (15%) | 0.194 | (20%) | 0.189 | (19%) |
| family:genus:species | 0.372 | (75%) | 0.375 | (77%) | 0.367 | (72%) | 0.367 | (73%) |
| **Sampling size (N)** | | | | | | | | |
| family | 249 | | 249 | | 40 | | 40 | |
| family:genus | 2576 | | 2576 | | 367 | | 367 | |
| family:genus:species | 10780 | | 10780 | | 2735 | | 2735 | |
| source | 293 | | | | 265 | | | |
| total | 79488 | | 79488 | | 49991 | | 49991 | |

**Table S4: Variance components of wood density across the taxonomic hierarchy (brms).** Shown are the results of four different random effects models (varying intercept models) fitted with the *brms* R package and the STAN software(Bürkner, 2018; Carpenter et al., 2017.The default model (M1) is fitted to the full set of measurements from individual plants and includes nested random effects for taxonomic groupings (species-level variation nested within genera, genus-level variation nested within families), as well as a crossed random effect for study methodology ("source"). Shown are also a simpler model (M2, no random effect for methodology) and the same two models fitted to a subset of records with higher quality (>= 3 records per species, >= 3 species per genus, >= 3 genera per family, M3 and M4). Throughout, the spread of residuals, σ, is itself modelled as varying between species with a nested random effects structure on log-scales (species within genera, genera within families). For simplicity, we also provide the residual variance on untransformed scales ($\sigma$ = root of variance of residuals, where residuals are calculated as posterior means). The percentage of total variance explained is provided in brackets next to each variance component (random effects + residual variance). Point estimates are posterior means, intervals 95% credibility intervals.

| *lme4* | **M1: Full dataset (incl. source effect)** | | **M2: Full dataset (no source effect)** | | **M3: High quality subset (incl. source effect)** | | **M4: High quality subset (no source effect)** | |
|---|---|---|---|---|---|---|---|---|
| **Wood density (g cm$^{-3}$)** | *Estimate* | *CI* | *Estimate* | *CI* | *Estimate* | *CI* | *Estimate* | *CI* |
| (Intercept) | 0.561 | [0.544; 0.577] | 0.555 | [0.540; 0.570] | 0.582 | [0.557; 0.606] | 0.576 | [0.553; 0.599] |
| **Random effects** | *Estimate* | *(%Var)* | *Estimate* | *(%Var)* | *Estimate* | *(%Var)* | *Estimate* | *(%Var)* |
| family | 0.096 | (29%) | 0.096 | (30%) | 0.063 | (16%) | 0.062 | (16%) |
| family:genus | 0.095 | (29%) | 0.096 | (30%) | 0.092 | (33%) | 0.093 | (36%) |
| family:genus:species | 0.079 | (20%) | 0.080 | (21%) | 0.079 | (24%) | 0.080 | (26%) |
| source | 0.046 | (7%) | | | 0.046 | (8%) | | |
| residual (σ) | 0.071 | (16%) | 0.074 | (18%) | 0.069 | (19%) | 0.073 | (22%) |
| **Sampling size (N)** | | | | | | | | |
| family | 249 | | 249 | | 40 | | 40 | |
| family:genus | 2576 | | 2576 | | 367 | | 367 | |
| family:genus:species | 10780 | | 10780 | | 2735 | | 2735 | |
| source | 293 | | | | 265 | | | |
| total | 79488 | | 79488 | | 49991 | | 49991 | |

**Table S5: Variance components of wood density across the taxonomic hierarchy (lme4).** Same as Table S4, but using reduced maximum likelihood modelling instead of a Bayesian approach (Bates et al., 2015) and not explicitly modelling the residual distribution. Intervals are 95% confidence intervals (Wald method).

| *brms/STAN* | M5: Minimum quality (incl. source effect) | | M6: Minimum quality (no source effect) | | M7: High quality (incl. source effect) | | M8: High quality (no source effect) | |
|---|---|---|---|---|---|---|---|---|
| **Wood density (g cm$^{-3}$)** | *Estimate* | *CI* | *Estimate* | *CI* | *Estimate* | *CI* | *Estimate* | *CI* |
| (Intercept) | 0.560 | [0.542; 0.578] | 0.564 | [0.547; 0.582] | 0.571 | [0.524; 0.615] | 0.575 | [0.536; 0.610] |
| **Random effects** | *Estimate* | *(%Var)* | *Estimate* | *(%Var)* | *Estimate* | *(%Var)* | *Estimate* | *(%Var)* |
| species | 0.098 | (72%) | 0.101 | (78%) | 0.100 | (58%) | 0.104 | (70%) |
| species:site | 0.025 | (5%) | 0.032 | (8%) | 0.035 | (7%) | 0.042 | (11%) |
| species:site:idplant | 0.017 | (2%) | 0.017 | (2%) | 0.028 | (5%) | 0.028 | (5%) |
| source | 0.035 | (9%) | | | 0.057 | (19%) | | |
| residual | 0.040 | (12%) | 0.040 | (12%) | 0.045 | (12%) | 0.045 | (13%) |
| **log(σ)** | *Estimate* | *CI* | *Estimate* | *CI* | *Estimate* | *CI* | *Estimate* | *CI* |
| (Intercept) | -3.015 | [-3.116; -2.916] | -3.029 | [-3.127; -2.929] | -3.049 | [-3.233; -2.870] | -3.040 | [-3.223; -2.859] |
| **Random effects** | *Estimate* | | *Estimate* | | *Estimate* | | *Estimate* | |
| species | 0.588 | | 0.588 | | 0.531 | | 0.538 | |
| **Sampling size (N)** | | | | | | | | |
| source | 95 | | | | 20 | | | |
| species | 147 | | 147 | | 35 | | 35 | |
| species:site | 1270 | | 1270 | | 233 | | 233 | |
| species:site:idplant | 14373 | | 14373 | | 1052 | | 1052 | |
| total | 19246 | | 19246 | | 2494 | | 2494 | |

**Table S6: Variance components of wood density within species (brms).** Shown are the results of four different random effects models fitted with *brms*/STAN (Bates et al., 2015; Carpenter et al., 2017), in order to partition within-species variance. The default model (M5) is fitted to a GWDD v.2 subset with minimum quality requirements (only species with >= 2 sites, at least one site with >= 2 plants, at least one plant with >= 2 records) and includes nested random effects (plant-level variation nested within sites, site-level variation nested within species), as well as a crossed random effect for study methodology ("source"). In addition, a simpler model is presented (M6, no random effect for methodology) and the same two models fitted to a higher quality subset (species with >= 3 sites, at least one site with >= 3 plants, at least one plant with >= 3 records, M7 and M8). As described in Table S4, σ is modelled on log-scales, but residual variance is calculated on the original scales. The percentage of total variance explained is provided in brackets next to each variance component. Point estimates are posterior means, intervals 95% credibility intervals.

| *lme4* | M5: Minimum quality (incl. source effect) | | M6: Minimum quality (no source effect) | | M7: High quality (incl. source effect) | | M8: High quality (no source effect) | |
|---|---|---|---|---|---|---|---|---|
| **Wood density (g cm$^{-3}$)** | *Estimate* | *CI* | *Estimate* | *CI* | *Estimate* | *CI* | *Estimate* | *CI* |
| (Intercept) | 0.560 | [0.541; 0.579] | 0.565 | [0.549; 0.581] | 0.573 | [0.530; 0.615] | 0.575 | [0.541; 0.608] |
| **Random effects** | *Estimate* | *(%Var)* | *Estimate* | *(%Var)* | *Estimate* | *(%Var)* | *Estimate* | *(%Var)* |
| species | 0.094 | (64%) | 0.097 | (68%) | 0.092 | (50%) | 0.096 | (58%) |
| species:site | 0.045 | (15%) | 0.050 | (18%) | 0.050 | (15%) | 0.060 | (23%) |
| species:site:idplant | 0.018 | (2%) | 0.018 | (2%) | 0.028 | (5%) | 0.028 | (5%) |
| source | 0.031 | (7%) | | | 0.054 | (17%) | | |
| residual | 0.041 | (12%) | 0.041 | (12%) | 0.047 | (13%) | 0.047 | (14%) |
| **Sampling size (N)** | | | | | | | | |
| source | 95 | | | | 20 | | | |
| species | 147 | | 147 | | 35 | | 35 | |
| species:site | 1270 | | 1270 | | 233 | | 233 | |
| species:site:idplant | 14373 | | 14373 | | 1052 | | 1052 | |
| total | 19246 | | 19246 | | 2494 | | 2494 | |

**Table S7: Variance components of wood density within species (lme4).** Same as Table S6, but using reduced maximum likelihood modelling instead of a Bayesian approach (Bates et al., 2015) and not explicitly modelling the residual distribution. Intervals are 95% confidence intervals (Wald method).

| | M9: Sapwood (brms) | | M9: Sapwood (lme4) | | M10: Branchwood (brms) | | M10: Branchwood (lme4) | |
|---|---|---|---|---|---|---|---|---|
| **Wood density (g cm$^{-3}$)** | *Estimate* | *CI* | *Estimate* | *CI* | *Estimate* | *CI* | *Estimate* | *CI* |
| (Intercept) | 0.572 | [0.533; 0.613] | 0.575 | [0.536; 0.613] | 0.601 | [0.593; 0.609] | 0.603 | [0.595; 0.611] |
| sapwood | -0.001 | [-0.028; 0.025] | -0.003 | [-0.030; 0.024] | | | | |
| branch | | | | | -0.023 | [-0.029; -0.017] | -0.027 | [-0.033; -0.021] |
| **Random effects** | *Estimate* | | *Estimate* | | *Estimate* | | *Estimate* | |
| species | 0.174 | | 0.174 | | 0.137 | | 0.139 | |
| source | 0.035 | | 0.032 | | 0.044 | | 0.045 | |
| sapwood | 0.060 | | 0.055 | | | | | |
| branch | | | | | 0.075 | | 0.083 | |
| residual | 0.068 | | 0.080 | | 0.065 | | 0.067 | |
| **log(σ)** | *Estimate* | *CI* | | | *Estimate* | *CI* | | |
| | -2.787 | [-2.925; -2.655] | | | -2.714 | [-2.738; -2.690] | | |
| **Random effects** | *Estimate* | | | | *Estimate* | | | |
| species | 0.448 | | | | 0.401 | | | |
| **Sampling size (N)** | | | | | | | | |
| source | 15 | | 15 | | 529 | | 529 | |
| species | 150 | | 150 | | 2018 | | 2018 | |
| total | 679 | | 679 | | 48494 | | 48494 | |

**Table S8: Within-plant effects on wood density variation (brms + lme4).** Shown are the results of two different mixed effects models (random intercept/random slope models) fitted either with *brms*/STAN software (Bürkner, 2018; Carpenter et al., 2017) or *lme4* (Bates et al., 2015) to explore within-plant variation in wood density. In M9, we model wood density as varying from heartwood to sapwood via a fixed effect for heartwood-sapwood (sapwood == 0 vs. sapwood == 1) as well as random intercepts and slopes at species level and a crossed random effect for study methodology ("source"). M10 has the same model structure, but with a fixed effect and random slopes for the trunkwood-branchwood distinction (branch == 0 vs. branch == 1). In the Bayesian models (*brms*), we also allow the spread of residuals (σ) to vary on log-scales across species. Intervals are 95% credibility/confidence intervals (Wald method for ML estimates).

| *Model* | | *Estimate* | *CI* |
|---|---|---|---|
| **M11: Sapwood-Heartwood** | Intercept | 0.160 | [0.106; 0.210] |
| | Slope | 0.673 | [0.589; 0.764] |
| | | | r = 0.78, $n_{species}$ = 150 |
| **M12: Branchwood-Trunkwood** | Intercept | 0.123 | [0.100; 0.145] |
| | Slope | 0.757 | [0.721; 0.795] |
| | | | r = 0.67, $n_{species}$ = 2018 |
| **M13: Branchwood-Trunkwood (sapwood only)** | Intercept | 0.062 | [0.020; 0.102] |
| | Slope | 0.871 | [0.808; 0.938] |
| | | | r = 0.76, $n_{species}$ = 523 |
| **M14: Branchwood-Trunkwood (high quality subset)** | Intercept | 0.070 | [0.006; 0.128] |
| | Slope | 0.861 | [0.761; 0.972] |
| | | | r = 0.76, $n_{species}$ = 189 |
| **M15: Branchwood-Trunkwood (individual plants)** | Intercept | 0.014 | [0.004; 0.025] |
| | Slope | 0.988 | [0.969; 1.009] |
| | | | r = 0.85, n = 3,527, $n_{species}$ = 145 |

**Table S9: Within-plant convergence in wood density (lmodel2).** Shown are results from five Major Axis regression models, fitted with the *lmodel2* package in R (Legendre, 2018), to explore whether wood density converges towards sapwood and branchwood. Model M11 regresses mean sapwood densities of 150 species against their mean heartwood densities, Model M12 regresses mean branchwood densities of 2,018 species against their mean trunkwood densities. M13-15 are equivalent to M12, but M13 uses only sapwood measurements from branches and trunks, M14 a high-quality subset of measurements (>= 5 measurements of branches and >= 5 measurements of trunks), and M15 only measurements where both branch and trunk samples have been taken from the same plant, and regresses them at the individual plant instead of the species level. Intervals are parametric 95% confidence intervals.

| *brms/STAN* | M16: Full dataset CHELSA 1 km | | M17: Full dataset TerraClimate 5 km | | M18: High quality subset CHELSA 1 km | | M19: High quality subset TerraClimate 5 km | |
|---|---|---|---|---|---|---|---|---|
| **Wood density (g cm$^{-3}$)** | *Estimate* | *CI* | *Estimate* | *CI* | *Estimate* | *CI* | *Estimate* | *CI* |
| (Intercept) | 0.554 | [0.544; 0.564] | 0.559 | [0.548; 0.569] | 0.540 | [0.526; 0.554] | 0.546 | [0.532; 0.559] |
| Temperature (intrasp) | 0.012 | [0.002; 0.022] | 0.003 | [-0.007; 0.013] | 0.013 | [0.001; 0.024] | 0.004 | [-0.007; 0.015] |
| Water deficit (intrasp) | 0.010 | [0.005; 0.015] | 0.012 | [0.006; 0.018] | 0.008 | [0.001; 0.014] | 0.011 | [0.005; 0.018] |
| Wind speed (intrasp) | -0.003 | [-0.007; 0.000] | -0.006 | [-0.011; -0.001] | -0.001 | [-0.006; 0.003] | -0.003 | [-0.008; 0.002] |
| Sand fraction. (intrasp) | 0.003 | [-0.002; 0.007] | -0.003 | [-0.007; 0.002] | 0.006 | [0.002; 0.011] | 0.004 | [-0.001; 0.008] |
| Soil pH (intrasp) | -0.003 | [-0.008; 0.002] | -0.005 | [-0.010; -0.000] | -0.002 | [-0.008; 0.003] | -0.005 | [-0.010; 0.000] |
| Cat. ex. cap. (intrasp) | 0.009 | [0.004; 0.013] | 0.001 | [-0.004; 0.006] | 0.003 | [-0.002; 0.008] | -0.002 | [-0.008; 0.004] |
| Temperature (species) | 0.022 | [0.009; 0.034] | 0.013 | [0.000; 0.025] | 0.002 | [-0.015; 0.019] | -0.005 | [-0.024; 0.012] |
| Water deficit (species) | 0.038 | [0.027; 0.048] | 0.036 | [0.025; 0.047] | 0.045 | [0.027; 0.062] | 0.042 | [0.026; 0.058] |
| Wind speed (species) | -0.017 | [-0.027; -0.007] | -0.013 | [-0.022; -0.004] | -0.021 | [-0.038; -0.003] | -0.016 | [-0.032; -0.001] |
| Sand fraction. (species) | 0.002 | [-0.008; 0.012] | -0.000 | [-0.010; 0.011] | -0.008 | [-0.025; 0.010] | -0.008 | [-0.025; 0.010] |
| Soil pH (species) | -0.013 | [-0.024; -0.001] | -0.014 | [-0.028; -0.001] | -0.010 | [-0.029; 0.008] | -0.018 | [-0.038; 0.002] |
| Cat. ex. cap. (species) | 0.020 | [0.008; 0.032] | 0.001 | [-0.010; 0.012] | 0.012 | [-0.005; 0.030] | -0.001 | [-0.017; 0.016] |
| **Random effects** | *Estimate* | | *Estimate* | | *Estimate* | | *Estimate* | |
| species | 0.114 | | 0.114 | | 0.112 | | 0.112 | |
| source | 0.051 | | 0.051 | | 0.053 | | 0.052 | |
| Temperature (intrasp) | 0.058 | | 0.061 | | 0.060 | | 0.059 | |
| Water deficit (intrasp) | 0.038 | | 0.032 | | 0.039 | | 0.033 | |
| Wind speed (intrasp) | 0.019 | | 0.025 | | 0.022 | | 0.025 | |
| Sand fraction. (intrasp) | 0.029 | | 0.033 | | 0.018 | | 0.019 | |
| Soil pH (intrasp) | 0.035 | | 0.030 | | 0.029 | | 0.024 | |
| Cat. ex. cap. (intrasp) | 0.024 | | 0.031 | | 0.021 | | 0.032 | |
| residual | 0.062 | | 0.062 | | 0.054 | | 0.055 | |
| **log(**σ) | *Estimate* | *CI* | *Estimate* | *CI* | *Estimate* | *CI* | *Estimate* | *CI* |
| (Intercept) | -2.772 | [-2.802; -2.743] | -2.766 | [-2.796; -2.736] | -2.860 | [-2.906; -2.814] | -2.854 | [-2.902; -2.808] |
| **Random effects** | *Estimate* | | *Estimate* | | *Estimate* | | *Estimate* | |
| species | 0.497 | | 0.500 | | 0.463 | | 0.468 | |

| | | | | |
|---|---|---|---|---|
| **Sampling size (N)** | | | | |
| source | 366 | 366 | 338 | 338 |
| species | 2160 | 2160 | 692 | 692 |
| total | 41879 | 41893 | 30128 | 30128 |

**Table S10: Environmental predictors of wood density variation (brms).** Shown are the results of four different mixed effects models (varying intercept/varying slope models) fitted with *brms*/STAN software (Bürkner, 2018; Carpenter et al., 2017) to explore environmental predictors of wood density variation within and across species. All models have the same structure, relying on three environmental predictors (annual means of temperature, water deficit, and wind speed) and three edaphic predictors (mean sand content, mean pH, mean cation exchange capacity). All six predictors are standardized (scaled) and split into species mean values (indicated by "species") and within-species deviations from the species means (indicated by "intrasp"). Species are allowed to vary both in their intercept and in their within-species effects (varying slopes). In addition, we include a crossed random effect for the measurement source/methodology, and allow the distributional parameter σ to vary across species. M16 and M17 are fitted to all species in the GWDD v.2 with wood density records from at least two distinct geographic locations (explicit coordinates), but differ in their environmental layers and scale of aggregation: M16 uses 1 km resolution data from CHELSA (Brun et al., 2022; Karger et al., 2017) and *soilgrids* predictions pre-aggregated at 1 km (Hengl et al., 2017). M17 uses ~4-5 km resolution data from TerraClimate (Abatzoglou et al., 2018) in conjunction with pre-aggregated 5 km *soilgrids* data. M18 and M19 use the same model structures, but restrict wood density records to species that display large within-species environmental or edaphic gradients, including only species for which the range of at least one environmental or edaphic predictor is in the top 10% of all species' ranges for that predictor. Point estimates are posterior means, intervals 95% credibility intervals. A visualization of effect sizes can be found in Figures S7-8.

| *lme4* | M16: Full dataset CHELSA 1 km | | M17: Full dataset TerraClimate 5 km | | M18: High quality subset CHELSA 1 km | | M19: High quality subset TerraClimate 5 km | |
|---|---|---|---|---|---|---|---|---|
| **Wood density (g cm$^{-3}$)** | *Estimate* | *CI* | *Estimate* | *CI* | *Estimate* | *CI* | *Estimate* | *CI* |
| (Intercept) | 0.555 | [0.545; 0.565] | 0.559 | [0.548; 0.569] | 0.540 | [0.526; 0.554] | 0.544 | [0.530; 0.558] |
| Temperature (intrasp) | 0.013 | [0.002; 0.024] | 0.001 | [-0.009; 0.012] | 0.014 | [0.002; 0.026] | 0.002 | [-0.010; 0.014] |
| Water deficit (intrasp) | 0.010 | [0.004; 0.015] | 0.014 | [0.007; 0.021] | 0.007 | [0.000; 0.014] | 0.012 | [0.004; 0.021] |
| Wind speed (intrasp) | -0.003 | [-0.007; 0.000] | -0.008 | [-0.013; -0.004] | -0.002 | [-0.007; 0.003] | -0.005 | [-0.011; 0.001] |
| Sand fraction. (intrasp) | 0.004 | [-0.002; 0.009] | -0.004 | [-0.010; 0.002] | 0.010 | [0.004; 0.016] | 0.002 | [-0.005; 0.009] |
| Soil pH (intrasp) | -0.005 | [-0.011; 0.001] | -0.007 | [-0.013; -0.001] | -0.001 | [-0.007; 0.005] | -0.005 | [-0.012; 0.001] |
| Cat. ex. cap. (intrasp) | 0.012 | [0.007; 0.017] | 0.003 | [-0.003; 0.009] | 0.006 | [0.001; 0.011] | -0.000 | [-0.007; 0.006] |
| Temperature (species) | 0.022 | [0.010; 0.035] | 0.013 | [0.000; 0.025] | 0.003 | [-0.014; 0.020] | -0.005 | [-0.023; 0.013] |
| Water deficit (species) | 0.038 | [0.027; 0.048] | 0.035 | [0.024; 0.046] | 0.044 | [0.026; 0.062] | 0.042 | [0.027; 0.058] |
| Wind speed (species) | -0.018 | [-0.028; -0.009] | -0.015 | [-0.023; -0.006] | -0.021 | [-0.039; -0.004] | -0.016 | [-0.031; -0.001] |
| Sand fraction. (species) | 0.005 | [-0.005; 0.016] | 0.002 | [-0.009; 0.013] | -0.004 | [-0.021; 0.014] | -0.007 | [-0.025; 0.011] |
| Soil pH (species) | -0.012 | [-0.024; -0.000] | -0.013 | [-0.026; 0.000] | -0.008 | [-0.027; 0.010] | -0.016 | [-0.036; 0.004] |
| Cat. ex. cap. (species) | 0.022 | [0.010; 0.033] | 0.002 | [-0.009; 0.014] | 0.015 | [-0.003; 0.032] | 0.001 | [-0.016; 0.018] |
| **Random effects** | *Estimate* | | *Estimate* | | *Estimate* | | *Estimate* | |
| species | 0.116 | | 0.116 | | 0.113 | | 0.113 | |
| source | 0.048 | | 0.048 | | 0.053 | | 0.053 | |
| Temperature (intrasp) | 0.067 | | 0.068 | | 0.063 | | 0.064 | |
| Water deficit (intrasp) | 0.052 | | 0.048 | | 0.044 | | 0.048 | |
| Wind speed (intrasp) | 0.020 | | 0.026 | | 0.028 | | 0.028 | |
| Sand fraction. (intrasp) | 0.047 | | 0.054 | | 0.036 | | 0.042 | |
| Soil pH (intrasp) | 0.051 | | 0.045 | | 0.034 | | 0.039 | |
| Cat. ex. cap. (intrasp) | 0.027 | | 0.038 | | 0.024 | | 0.037 | |
| residual | 0.063 | | 0.064 | | 0.055 | | 0.055 | |
| **Sampling size (N)** | | | | | | | | |
| source | 366 | | 366 | | 338 | | 338 | |
| species | 2160 | | 2160 | | 692 | | 692 | |
| total | 41879 | | 41893 | | 30128 | | 30128 | |

**Table S11: Environmental predictors of wood density variation (lme4).** Same as Table S10, but using reduced maximum likelihood modelling instead of a Bayesian approach (Bates et al., 2015) and not explicitly modelling the residual distribution. Intervals are 95% confidence intervals (Wald method).

| *brms/STAN* | **M20: Tropical species CHELSA 1 km** | | **M21: Tropical species TerraClimate 5 km** | | **M22: Extratropical species CHELSA 1 km** | | **M23: Extratropical species TerraClimate 5 km** | |
|---|---|---|---|---|---|---|---|---|
| **Wood density (g cm$^{-3}$)** | *Estimate* | *CI* | *Estimate* | *CI* | *Estimate* | *CI* | *Estimate* | *CI* |
| (Intercept) | 0.528 | [0.475; 0.582] | 0.517 | [0.467; 0.568] | 0.569 | [0.549; 0.588] | 0.561 | [0.540; 0.581] |
| Temperature (intrasp) | 0.013 | [-0.006; 0.032] | -0.000 | [-0.019; 0.018] | 0.016 | [0.002; 0.030] | 0.006 | [-0.009; 0.021] |
| Water deficit (intrasp) | 0.012 | [0.004; 0.021] | 0.021 | [0.008; 0.034] | 0.001 | [-0.007; 0.009] | 0.011 | [0.003; 0.020] |
| Wind speed (intrasp) | -0.004 | [-0.011; 0.003] | -0.021 | [-0.031; -0.011] | 0.002 | [-0.003; 0.007] | 0.004 | [-0.001; 0.010] |
| Sand fraction. (intrasp) | 0.004 | [-0.005; 0.014] | -0.009 | [-0.019; 0.001] | 0.004 | [0.001; 0.007] | 0.002 | [-0.001; 0.006] |
| Soil pH (intrasp) | -0.013 | [-0.021; -0.004] | -0.012 | [-0.023; -0.002] | 0.001 | [-0.004; 0.006] | -0.004 | [-0.009; 0.001] |
| Cat. ex. cap. (intrasp) | 0.019 | [0.010; 0.029] | -0.003 | [-0.014; 0.008] | 0.001 | [-0.004; 0.005] | -0.002 | [-0.008; 0.003] |
| Temperature (species) | 0.022 | [-0.030; 0.072] | -0.010 | [-0.056; 0.036] | 0.043 | [0.022; 0.064] | 0.042 | [0.020; 0.065] |
| Water deficit (species) | 0.059 | [0.038; 0.080] | 0.078 | [0.051; 0.104] | 0.033 | [0.009; 0.056] | 0.027 | [0.010; 0.045] |
| Wind speed (species) | -0.033 | [-0.056; -0.010] | -0.049 | [-0.068; -0.029] | 0.001 | [-0.018; 0.020] | 0.015 | [-0.004; 0.035] |
| Sand fraction. (species) | 0.013 | [-0.017; 0.042] | -0.009 | [-0.039; 0.022] | -0.007 | [-0.025; 0.011] | -0.013 | [-0.031; 0.005] |
| Soil pH (species) | -0.031 | [-0.055; -0.007] | -0.048 | [-0.079; -0.017] | -0.015 | [-0.041; 0.011] | -0.010 | [-0.033; 0.013] |
| Cat. ex. cap. (species) | 0.027 | [-0.001; 0.056] | -0.021 | [-0.051; 0.009] | 0.008 | [-0.009; 0.025] | 0.003 | [-0.014; 0.021] |
| **Random effects** | *Estimate* | | *Estimate* | | *Estimate* | | *Estimate* | |
| species | 0.122 | | 0.121 | | 0.088 | | 0.087 | |
| source | 0.048 | | 0.049 | | 0.054 | | 0.052 | |
| Temperature (intrasp) | 0.049 | | 0.049 | | 0.065 | | 0.075 | |
| Water deficit (intrasp) | 0.038 | | 0.044 | | 0.037 | | 0.034 | |
| Wind speed (intrasp) | 0.012 | | 0.029 | | 0.020 | | 0.023 | |
| Sand fraction. (intrasp) | 0.044 | | 0.051 | | 0.010 | | 0.011 | |
| Soil pH (intrasp) | 0.043 | | 0.043 | | 0.020 | | 0.018 | |
| Cat. ex. cap. (intrasp) | 0.034 | | 0.045 | | 0.016 | | 0.023 | |
| residual | 0.085 | | 0.085 | | 0.049 | | 0.049 | |
| **log(**$\sigma$**)** | *Estimate* | *CI* | *Estimate* | *CI* | *Estimate* | *CI* | *Estimate* | *CI* |
| (Intercept) | -2.603 | [-2.645; -2.561] | -2.599 | [-2.643; -2.557] | -3.130 | [-3.195; -3.068] | -3.129 | [-3.192; -3.065] |
| **Random effects** | *Estimate* | | *Estimate* | | *Estimate* | | *Estimate* | |
| species | 0.437 | | 0.442 | | 0.415 | | 0.416 | |

| | | | | |
|---|---|---|---|---|
| **Sampling size (N)** | | | | |
| source | 140 | 140 | 253 | 253 |
| species | 700 | 700 | 247 | 247 |
| total | 8783 | 8783 | 26437 | 26439 |

**Table S12: Environmental predictors of wood density variation, in and outside of tropics (brms).** Same as Table S10, but splitting the original data set into tropical species (>= 3 occurrences in the GWDD v.2 between 23.5 N and 23.5 S) and extratropical species (>= 3 occurrences outside of 23.5 N and 23.5 S). Point estimates are posterior means, intervals 95% credibility intervals. A visualization of effect sizes can be found in Figures S9-10.

| *lme4* | **M20: Tropical species CHELSA 1 km** | | **M21: Tropical species TerraClimate 5 km** | | **M22: Extratropical species CHELSA 1 km** | | **M23: Extratropical species TerraClimate 5 km** | |
|---|---|---|---|---|---|---|---|---|
| **Wood density (g cm$^{-3}$)** | *Estimate* | *CI* | *Estimate* | *CI* | *Estimate* | *CI* | *Estimate* | *CI* |
| (Intercept) | 0.520 | [0.466; 0.574] | 0.515 | [0.464; 0.567] | 0.568 | [0.549; 0.588] | 0.560 | [0.540; 0.580] |
| Temperature (intrasp) | 0.022 | [0.002; 0.042] | -0.003 | [-0.022; 0.016] | 0.017 | [0.003; 0.030] | 0.004 | [-0.012; 0.019] |
| Water deficit (intrasp) | 0.009 | [0.001; 0.018] | 0.020 | [0.007; 0.033] | 0.001 | [-0.007; 0.009] | 0.014 | [0.006; 0.022] |
| Wind speed (intrasp) | -0.001 | [-0.008; 0.006] | -0.017 | [-0.027; -0.007] | 0.002 | [-0.002; 0.007] | 0.005 | [-0.001; 0.010] |
| Sand fraction. (intrasp) | 0.005 | [-0.005; 0.014] | -0.009 | [-0.019; 0.001] | 0.004 | [0.000; 0.007] | 0.002 | [-0.001; 0.005] |
| Soil pH (intrasp) | -0.014 | [-0.023; -0.006] | -0.015 | [-0.026; -0.004] | 0.002 | [-0.003; 0.007] | -0.004 | [-0.008; 0.000] |
| Cat. ex. cap. (intrasp) | 0.021 | [0.012; 0.030] | -0.002 | [-0.013; 0.009] | 0.001 | [-0.003; 0.005] | -0.002 | [-0.007; 0.004] |
| Temperature (species) | 0.034 | [-0.018; 0.085] | -0.008 | [-0.055; 0.039] | 0.043 | [0.023; 0.063] | 0.042 | [0.020; 0.064] |
| Water deficit (species) | 0.059 | [0.037; 0.080] | 0.079 | [0.052; 0.106] | 0.035 | [0.011; 0.059] | 0.028 | [0.011; 0.045] |
| Wind speed (species) | -0.031 | [-0.054; -0.007] | -0.049 | [-0.068; -0.029] | 0.001 | [-0.018; 0.020] | 0.015 | [-0.004; 0.034] |
| Sand fraction. (species) | 0.015 | [-0.016; 0.045] | -0.007 | [-0.037; 0.024] | -0.006 | [-0.023; 0.011] | -0.012 | [-0.030; 0.005] |
| Soil pH (species) | -0.033 | [-0.057; -0.009] | -0.050 | [-0.081; -0.019] | -0.016 | [-0.042; 0.010] | -0.010 | [-0.033; 0.012] |
| Cat. ex. cap. (species) | 0.032 | [0.003; 0.062] | -0.018 | [-0.048; 0.012] | 0.011 | [-0.007; 0.029] | 0.004 | [-0.013; 0.022] |
| **Random effects** | *Estimate* | | *Estimate* | | *Estimate* | | *Estimate* | |
| species | 0.121 | | 0.119 | | 0.087 | | 0.086 | |
| source | 0.045 | | 0.047 | | 0.053 | | 0.051 | |
| Temperature (intrasp) | 0.030 | | 0.029 | | 0.059 | | 0.074 | |
| Water deficit (intrasp) | 0.037 | | 0.041 | | 0.036 | | 0.032 | |
| Wind speed (intrasp) | 0.013 | | 0.020 | | 0.019 | | 0.020 | |
| Sand fraction. (intrasp) | 0.043 | | 0.050 | | 0.012 | | 0.010 | |
| Soil pH (intrasp) | 0.037 | | 0.037 | | 0.018 | | 0.014 | |
| Cat. ex. cap. (intrasp) | 0.011 | | 0.031 | | 0.014 | | 0.023 | |
| residual | 0.092 | | 0.093 | | 0.050 | | 0.050 | |
| **Sampling size (N)** | | | | | | | | |
| source | 140 | | 140 | | 253 | | 253 | |
| species | 700 | | 700 | | 247 | | 247 | |
| total | 8783 | | 8783 | | 26437 | | 26439 | |

**Table S13: Environmental predictors of wood density variation, in and outside of tropics (lme4).** Same as Table S12, but using reduced maximum likelihood modelling instead of a Bayesian approach (Bates et al., 2015) and not explicitly modelling the residual distribution. Intervals are 95% confidence intervals (Wald method).

| brms/STAN | M20: Tropical species CHELSA 1 km | | M21: Tropical species TerraClimate 5 km | | M22: Extratropical species CHELSA 1 km | | M23: Extratropical species TerraClimate 5 km | |
|---|---|---|---|---|---|---|---|---|
| **Wood density (g cm$^{-3}$)** | *Estimate (original)* | *Estimate (rescaled)* | *Estimate (original)* | *Estimate (rescaled)* | *Estimate (original)* | *Estimate (rescaled)* | *Estimate (original)* | *Estimate (rescaled)* |
| Temperature (intrasp) | 0.014 | 0.003 | 0.001 | 0.000 | 0.017 | 0.005 | 0.006 | 0.002 |
| Water deficit (intrasp) | 0.012 | 0.008 | 0.021 | 0.010 | 0.001 | 0.000 | 0.012 | 0.005 |
| Wind speed (intrasp) | -0.003 | -0.002 | -0.020 | -0.009 | 0.002 | 0.001 | 0.004 | 0.002 |
| Sand fraction. (intrasp) | 0.005 | 0.002 | -0.008 | -0.003 | 0.004 | 0.003 | 0.002 | 0.002 |
| Soil pH (intrasp) | -0.012 | -0.006 | -0.012 | -0.006 | 0.001 | 0.001 | -0.004 | -0.002 |
| Cat. ex. cap. (intrasp) | 0.019 | 0.008 | -0.003 | -0.001 | 0.001 | 0.001 | -0.002 | -0.001 |
| Temperature (species) | 0.030 | 0.033 | -0.004 | -0.005 | 0.045 | 0.041 | 0.045 | 0.041 |
| Water deficit (species) | 0.057 | 0.049 | 0.078 | 0.055 | 0.032 | 0.024 | 0.026 | 0.023 |
| Wind speed (species) | -0.031 | -0.029 | -0.047 | -0.053 | 0.003 | 0.002 | 0.016 | 0.011 |
| Sand fraction. (species) | 0.017 | 0.009 | -0.005 | -0.003 | -0.006 | -0.005 | -0.012 | -0.009 |
| Soil pH (species) | -0.031 | -0.024 | -0.048 | -0.037 | -0.014 | -0.012 | -0.009 | -0.007 |
| Cat. ex. cap. (species) | 0.031 | 0.020 | -0.017 | -0.011 | 0.009 | 0.008 | 0.004 | 0.004 |

**Table S14: Rescaled environmental effect sizes, in and outside of tropics (brms).** Same as fixed effects shown in Table S12, but extended to include separately rescaled estimates of within-species and among-species effects. Rescaling was carried out by calculating average within-species and among-species standard deviations for each predictor and then multiplying effect sizes with these standard deviations. The rescaled effect sizes thus correspond to the effect sizes one might observe across a species actual range.

| | M24: Gymnosperms CHELSA 1 km (brms) | | M24: Gymnosperms CHELSA 1 km (lme4) | | M25: Gymnosperms TerraClimate 5 km (brms) | | M25: Gymnosperms TerraClimate 5 km (lme4) | |
|---|---|---|---|---|---|---|---|---|
| **Wood density (g cm$^{-3}$)** | *Estimate* | *CI* | *Estimate* | *CI* | *Estimate* | *CI* | *Estimate* | *CI* |
| (Intercept) | 0.476 | [0.452; 0.499] | 0.477 | [0.457; 0.497] | 0.471 | [0.447; 0.494] | 0.469 | [0.449; 0.489] |
| Temperature (intrasp) | 0.021 | [-0.008; 0.047] | 0.035 | [0.013; 0.057] | -0.001 | [-0.042; 0.036] | -0.011 | [-0.052; 0.030] |
| Water deficit (intrasp) | -0.008 | [-0.025; 0.009] | -0.011 | [-0.028; 0.005] | 0.014 | [-0.008; 0.036] | 0.022 | [0.005; 0.040] |
| Wind speed (intrasp) | 0.010 | [0.002; 0.021] | 0.013 | [0.004; 0.022] | 0.006 | [-0.005; 0.018] | 0.006 | [-0.004; 0.016] |
| Sand fraction. (intrasp) | 0.000 | [-0.007; 0.007] | -0.003 | [-0.011; 0.004] | 0.002 | [-0.006; 0.009] | 0.001 | [-0.007; 0.008] |
| Soil pH (intrasp) | -0.010 | [-0.022; 0.002] | -0.012 | [-0.021; -0.003] | -0.015 | [-0.027; -0.005] | -0.015 | [-0.022; -0.007] |
| Cat. ex. cap. (intrasp) | 0.005 | [-0.005; 0.015] | 0.003 | [-0.006; 0.012] | 0.002 | [-0.007; 0.012] | 0.003 | [-0.007; 0.012] |
| Temperature (species) | 0.004 | [-0.019; 0.029] | -0.002 | [-0.022; 0.019] | -0.006 | [-0.032; 0.021] | -0.000 | [-0.022; 0.021] |
| Water deficit (species) | 0.034 | [-0.001; 0.068] | 0.041 | [0.013; 0.068] | 0.041 | [0.008; 0.073] | 0.036 | [0.009; 0.064] |
| Wind speed (species) | -0.012 | [-0.039; 0.015] | -0.016 | [-0.038; 0.007] | -0.003 | [-0.026; 0.020] | 0.007 | [-0.012; 0.027] |
| Sand fraction. (species) | -0.002 | [-0.028; 0.023] | -0.015 | [-0.031; 0.001] | 0.005 | [-0.018; 0.028] | 0.004 | [-0.014; 0.023] |
| Soil pH (species) | -0.005 | [-0.035; 0.025] | -0.013 | [-0.034; 0.009] | -0.006 | [-0.037; 0.025] | -0.002 | [-0.026; 0.022] |
| Cat. ex. cap. (species) | -0.001 | [-0.027; 0.024] | -0.006 | [-0.025; 0.014] | -0.010 | [-0.035; 0.014] | -0.009 | [-0.027; 0.010] |
| **Random effects** | *Estimate* | | *Estimate* | | *Estimate* | | *Estimate* | |
| species | 0.052 | | 0.049 | | 0.050 | | 0.049 | |
| source | 0.053 | | 0.054 | | 0.052 | | 0.052 | |
| Temperature (intrasp) | 0.051 | | 0.046 | | 0.087 | | 0.106 | |
| Water deficit (intrasp) | 0.033 | | 0.036 | | 0.038 | | 0.043 | |
| Wind speed (intrasp) | 0.016 | | 0.023 | | 0.018 | | 0.022 | |
| Sand fraction. (intrasp) | 0.012 | | 0.015 | | 0.012 | | 0.015 | |
| Soil pH (intrasp) | 0.020 | | 0.021 | | 0.017 | | 0.012 | |
| Cat. ex. cap. (intrasp) | 0.015 | | 0.020 | | 0.014 | | 0.019 | |
| residual | 0.050 | | 0.050 | | 0.050 | | 0.051 | |
| **log(**σ) | *Estimate* | *CI* | | | *Estimate* | *CI* | | |
| (Intercept) | -3.041 | [-3.166; -2.920] | | | -3.031 | [-3.155; -2.905] | | |
| **Random effects** | *Estimate* | | | | *Estimate* | | | |
| species | 0.417 | | | | 0.424 | | | |

| | | | | |
|---|---|---|---|---|
| **Sampling size (N)** | | | | |
| source | 159 | 159 | 159 | 159 |
| species | 59 | 59 | 59 | 59 |
| total | 12089 | 12089 | 12089 | 12089 |

**Table S15: Environmental predictors of wood density variation in gymnosperms (brms + lme4).** Same as Tables S10-13, but with predictions only for gymnosperms. Shown are both models fits with the *brms* package and with *lme4*. Intervals are 95% credibility/confidence intervals (Wald method for ML estimates). Note how most effects are weak and overlap with 0. A visualization of effect sizes can be found in Figure S11.

| *Database* | *Method* | *samples for prediction* | *RMSE (g cm⁻³)* | *R2* | *$n_{species}$* |
|---|---|---|---|---|---|
| **GWDD v.2** | **Wood density means** | 0 / genus | 0.083 | 0.640 | 1557 |
| | | 1 | 0.084 | 0.730 | 1667 |
| | | 2 | 0.056 | 0.860 | 1667 |
| | **M26: Hierarchical model** | 0 / genus | 0.071 | 0.710 | 1557 |
| | | 1 | 0.053 | 0.840 | 1667 |
| | | 2 | 0.043 | 0.900 | 1667 |
| | **M27: Hierarchical model** branch & trunk | 0 / genus | 0.060 | 0.730 | 1557 |
| | | 1 | 0.046 | 0.840 | 1667 |
| | | 2 | 0.038 | 0.900 | 1667 |
| **GWDD v.1** | **Wood density means** | 0 / genus | 0.098 | 0.620 | 1447 |
| | | 1 | 0.095 | 0.740 | 1580 |
| | | 2 | 0.069 | 0.840 | 1312 |

**Table S16: Quality of wood density predictions at the species level.** Shown is to what extent the wood density of 1,667 species (all well-sampled in the GWDD v.2, with records from >= 5 sources) can be estimated when only a limited number of wood density measurements are available (0, 1 or 2). The baseline approach is to estimate species-level wood density from simple wood density means, either at genus level (0 samples) or averaging across the provided 1-2 samples. Shown are also two alternative hierarchical modelling approaches: M26, which is a simple random effects model, with a nested taxonomic structure plus extra random effects for study and measurement location (cf. Table S3), as well as M27, which extends M26 with a fixed effect for the trunkwood-branchwood distinction. Both models were refitted three times for each species, using both the local species-specific samples (0, 1, 2) and the remainder of the GWDD v.2. To reduce the computational burden, we used only the *lme4* package. Summary statistics of predictive power are the mean absolute error (MAE, g $cm^{-3}$), root mean square error (RMSE, g $cm^{-3}$) and $R^2$, calculated with reference to estimates based on a full set of measurements (>= 5 sources). A comparison with the GWDD v.1 is provided for completeness, again taking the GWDD v.2 values as reference. We note that in some cases, species were either not available for prediction (i.e., not recorded in GWDD v.1), or were the sole species in their genus, which also removed them from predictions at genus level.

| *Model* | *Extent* | ***RMSE (g cm$^{-3}$)*** *Number of local samples* 0 | 1 | 2 | 3 | ***R2*** *Number of local samples* 0 | 1 | 2 | 3 |
|---|---|---|---|---|---|---|---|---|---|
| **Species means** | Global | 0.089 | 0.088 | 0.087 | 0.086 | 0.690 | 0.700 | 0.710 | 0.710 |
| | Study | 0.093 | 0.091 | 0.087 | 0.086 | 0.660 | 0.680 | 0.710 | 0.710 |
| | Local | | 0.107 | 0.085 | 0.082 | | 0.590 | 0.720 | 0.740 |
| **M26: Hierarchical model** | Global | 0.088 | 0.086 | 0.085 | 0.084 | 0.700 | 0.710 | 0.720 | 0.730 |
| **M27: Hierarchical model** (branch & trunk) | Global | 0.087 | 0.086 | 0.084 | 0.083 | 0.710 | 0.710 | 0.730 | 0.730 |

**Table S17: Quality of wood density predictions at the individual plant level**. Shown is the predictive power of the same models as in Table S16, but now applied to predict individual-level wood density and tested at well-sampled sites from well-sampled studies (only species and studies with >= 3 locations per study and >= 4 measurements per site, $n_{species}$ = 318). For each approach, we show how well it is able to predict an individual wood density measurement when 0, 1, 2, or 3 measurements from the same locality and the same study (identical measurement methodology) are available. The baseline is provided by simple species means, calculated in three ways: 1/ using a combination of the 0, 1, 2, or 3 local samples and all measurements from elsewhere in the GWDD v.2 (“Global”), 2/ using a combination of the 0, 1, 2, or 3 local samples and all samples measured as part of the same study (“Study”), or 3/ using only the 0, 1, 2, or 3 local samples (“Local”). This is compared to two hierarchical modelling approaches (M26, M27) that model wood density with a nested taxonomic hierarchy as well as crossed random effects for study methodology and study site, thus implicitly adjusting predictions for methodological biases and local wood density shifts. Note that, assuming that within-species wood density variation is distributed with sd = 0.068 g cm$^{-3}$, we would expect an RMSE = 0.096 g cm$^{-3}$, i.e., sqrt(2 * $0.068^2$) when predicting a single tree’s wood density from another tree with no other knowledge about measurement location or the type of tissue sampled. The hierarchical models clearly outperform this expectation (RMSE = 0.086 g cm$^{-3}$).

#### C) Supplementary Figures

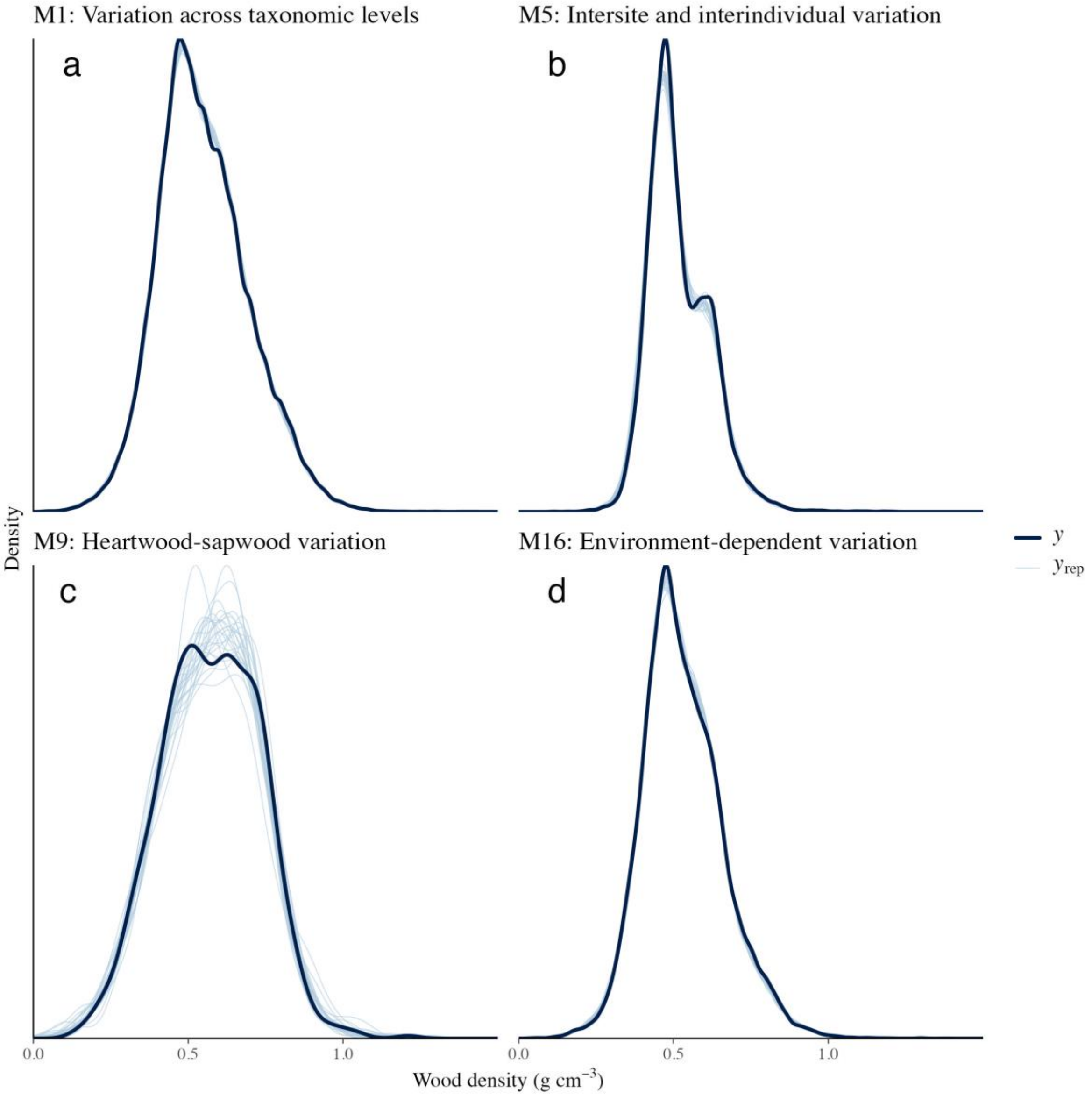

**Fig. S1: Sample posterior predictive checks.** Shown are posterior predictive checks for four of the models described in Table S3, using the default pp_check() function from the package *brms*. The black lines describe the distribution of wood density values for each of the GWDD v.2 subsets, the blue lines describe the modelled posterior density from 10 random posterior draws. All model fits successfully reproduce the data distribution, with no deviations in models M1 and M16 (panels a, d), a negligible underestimation of the mode in model M5 (panel c) and some uncertainty, though no bias, in model M9 (panel c).

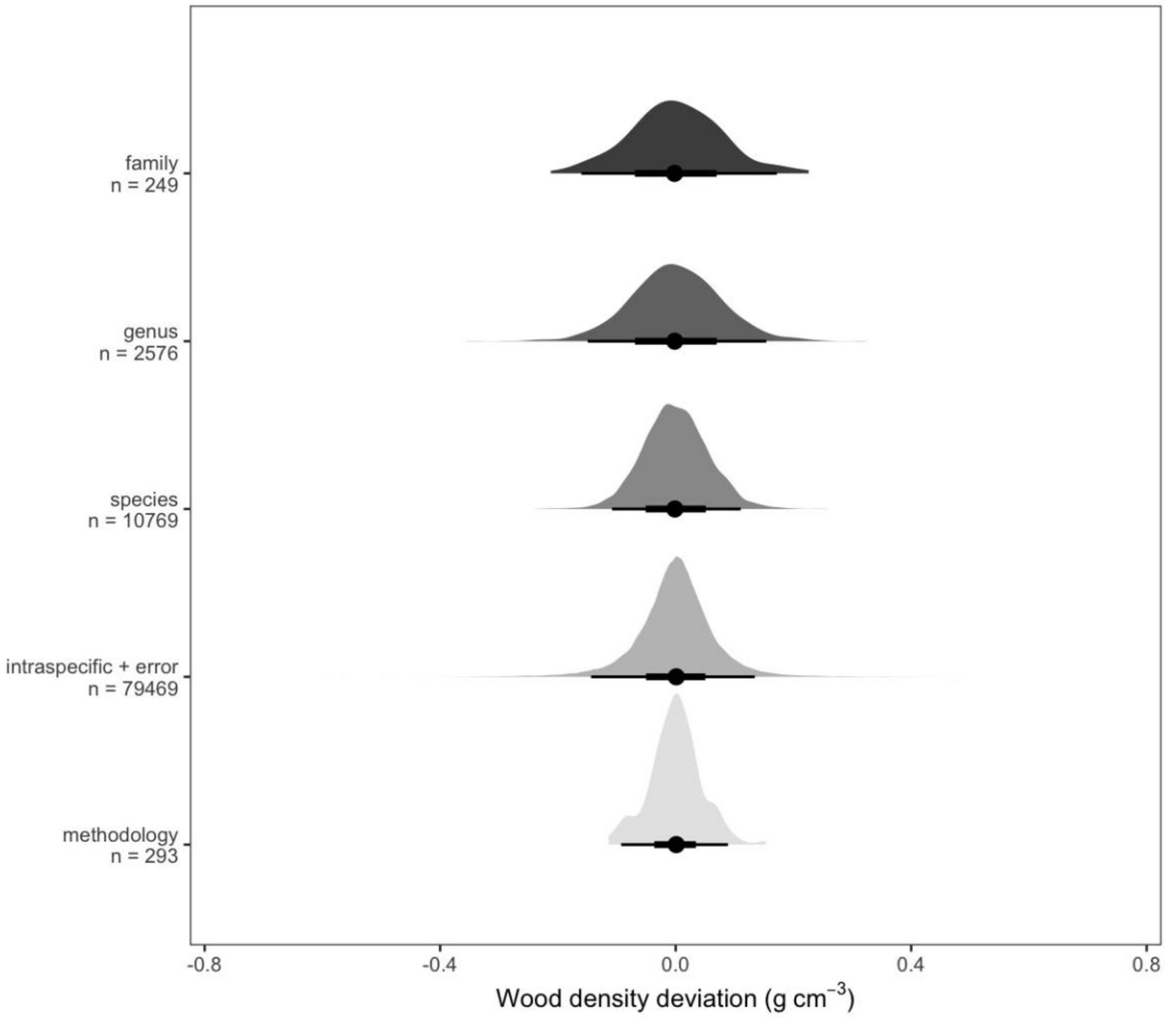

**Fig. S2: Variance components estimated from a Bayesian hierarchical model.** Shown are the different variance components as estimated from a Bayesian hierarchical model (M1 in Table S4). Shown are the nested random effects at family, genus and species level, as well as residual variation (intraspecific variation + error). We explicitly modelled sigma to assess the consistency of intraspecific variation, and also included a crossed random effect for methodology (i.e., the study where values were obtained from). The figure shows that variation at family and genus level is much larger than species or intraspecific variation, and that these are again larger than methodological effects. Note that the residuals (intraspecific variation + error) are overdispersed compared to a normal distribution, with a large number of outliers. Throughout, black dots indicate the median effect size and black intervals quantile ranges (66% and 95%).

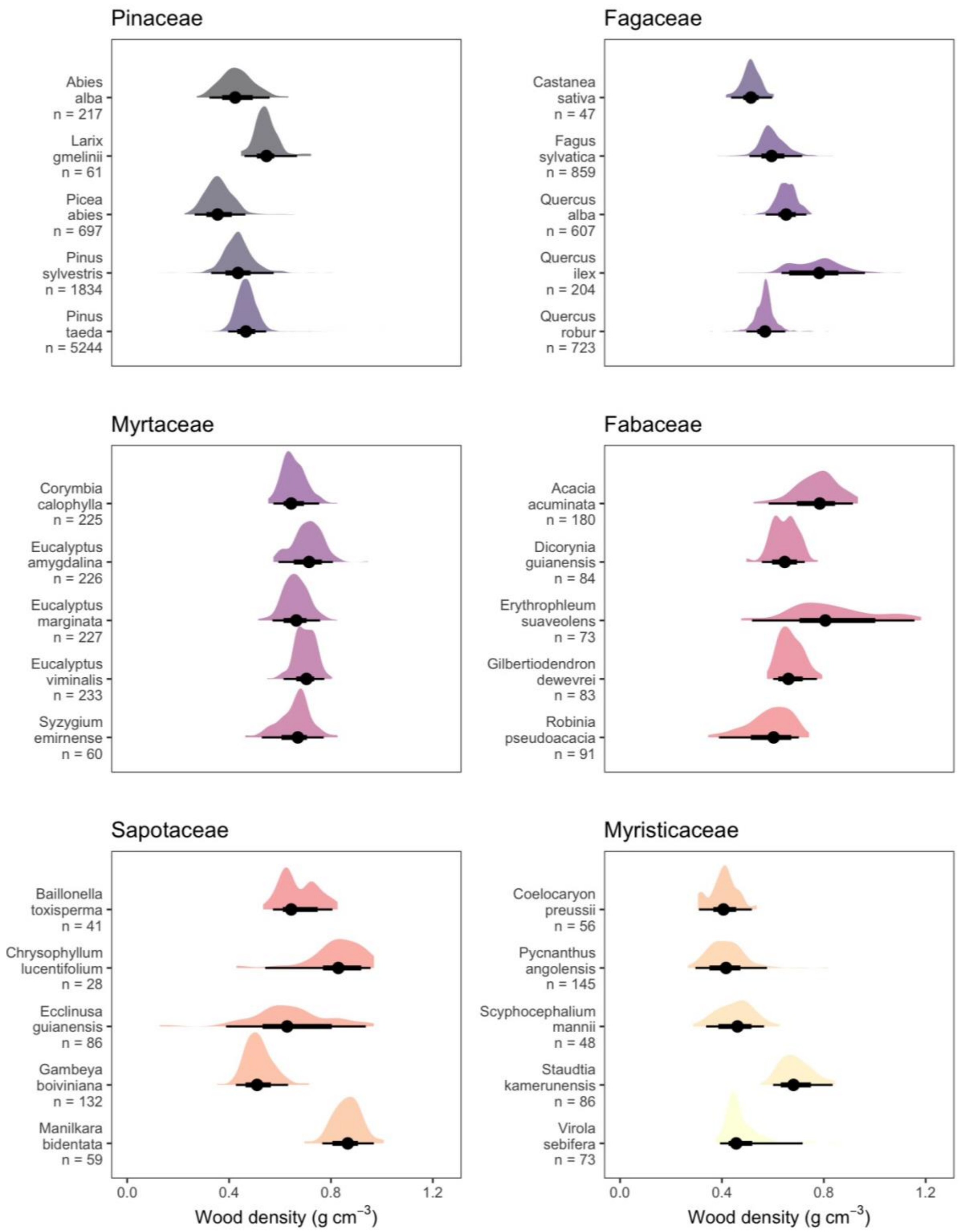

**Fig. S3: Intraspecific variation in selected species from six plant families.** Shown are estimated intraspecific wood density distributions for selected taxa, based on predictions from model M1 (Table S4). All values are based on wood density residuals, but have been corrected for methodological biases by subtracting study effects (cf. Fig. 1 inset and Table S4). Throughout, black dots indicate the median effect size and black intervals quantile ranges (66% and 95%).

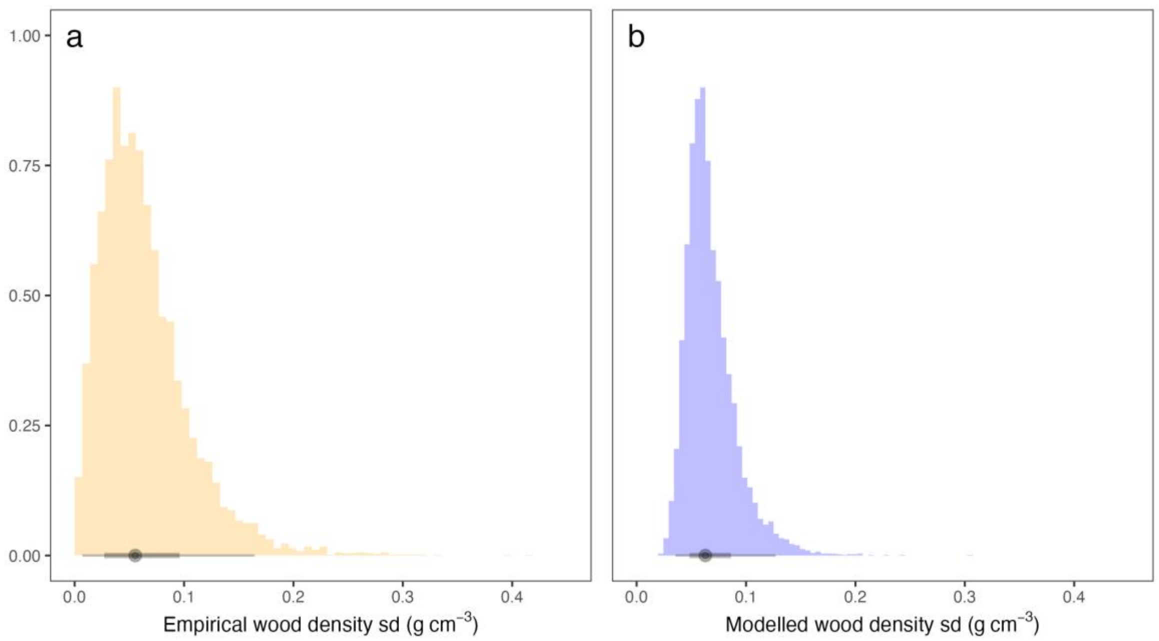

**Fig. S4: Consistency of the extent of intraspecific variation across taxa.** Shown is the distribution of empirical within-species standard deviations of wood density across all species with more than one plant-level record (a, $n_{species}$ = 6,361), as well as the distribution of inferred standard deviations (or sigma values) across the same species (b). The black dot indicates the median, the black intervals the corresponding quantile ranges (66% and 95%). Note that both follow approx. lognormal distributions, but the empirical standard deviations have a wider distribution. This is expected, as they include many species with very low sample sizes (e.g. 2-3 measurements), which the modelled distribution corrects for.

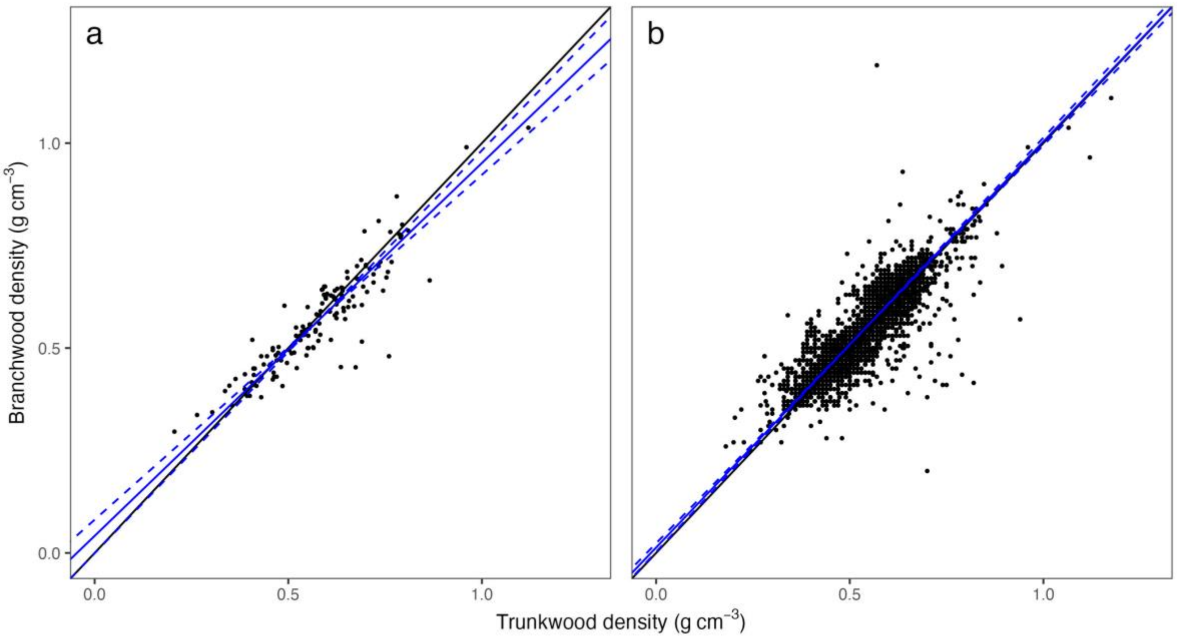

**Fig. S5: Wood density variation between trunkwood and branchwood, high quality regressions.** Shown is the same graph as in Fig. 3 (panels b and c), but for a subset of records where branch and trunk samples have been taken from the same individuals. Panel a) shows species mean values for both trunkwood and branchwood, panel b) shows each plant's mean trunkwood density and mean branchwood density. Blue lines are Major Axis regression lines, dashed lines the 95% CI. Note how the convergence observed in panel a) and in Fig. 3 disappears in panel b).

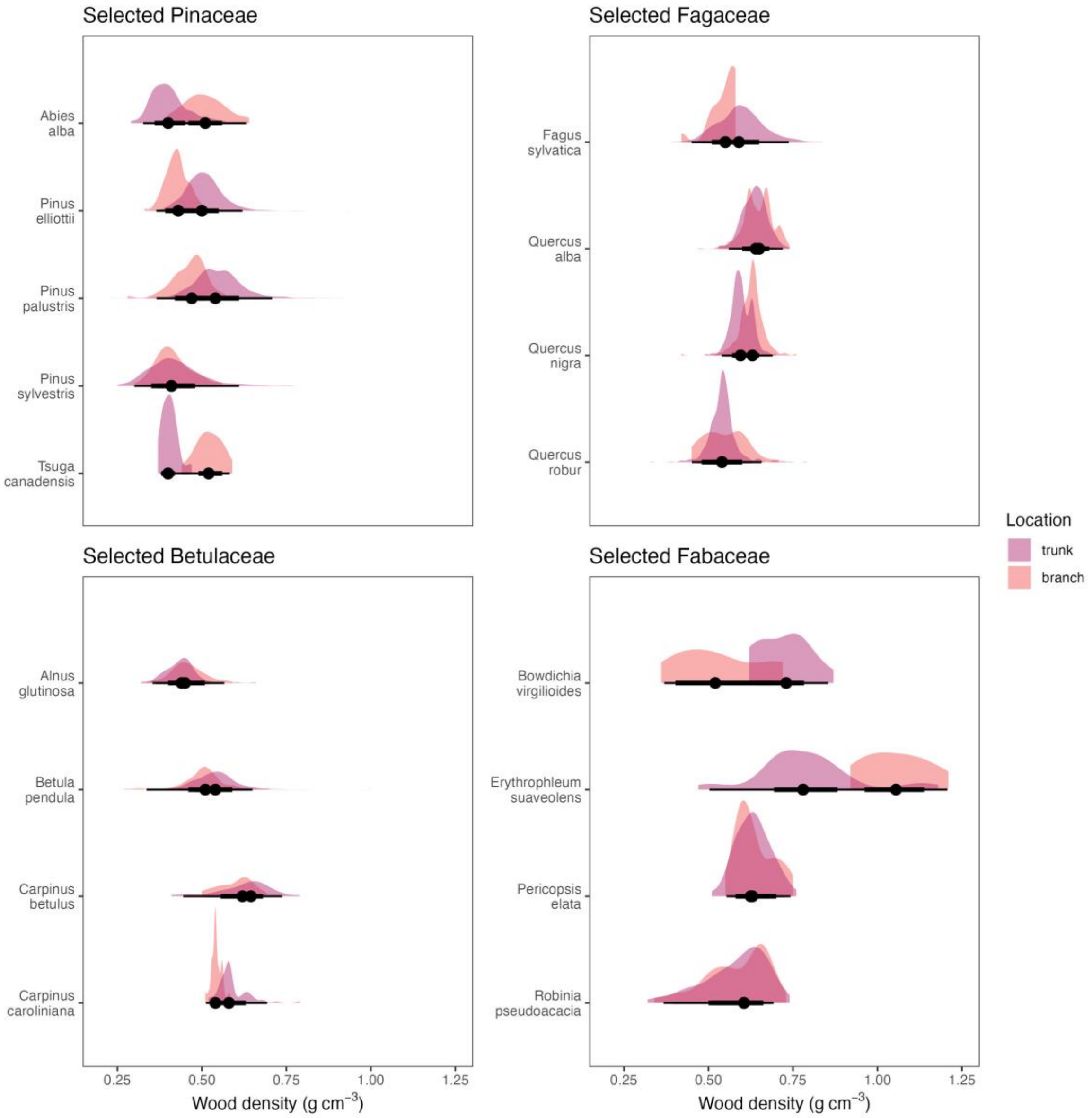

**Fig. S6: Wood density variation in trunkwood and in branchwood in selected species.** Shown are distributions of within-species variation in wood density for sample species from four families, split by within-tree location of samples (trunkwood or branchwood). Families and species were chosen to maximize sampling size. Black dots indicate the median effect size and black intervals quantile ranges (66% and 95%). Note how there is no clear clear convergence (i.e., branchwood distributions being closer to each other), nor a simple predictive pattern.

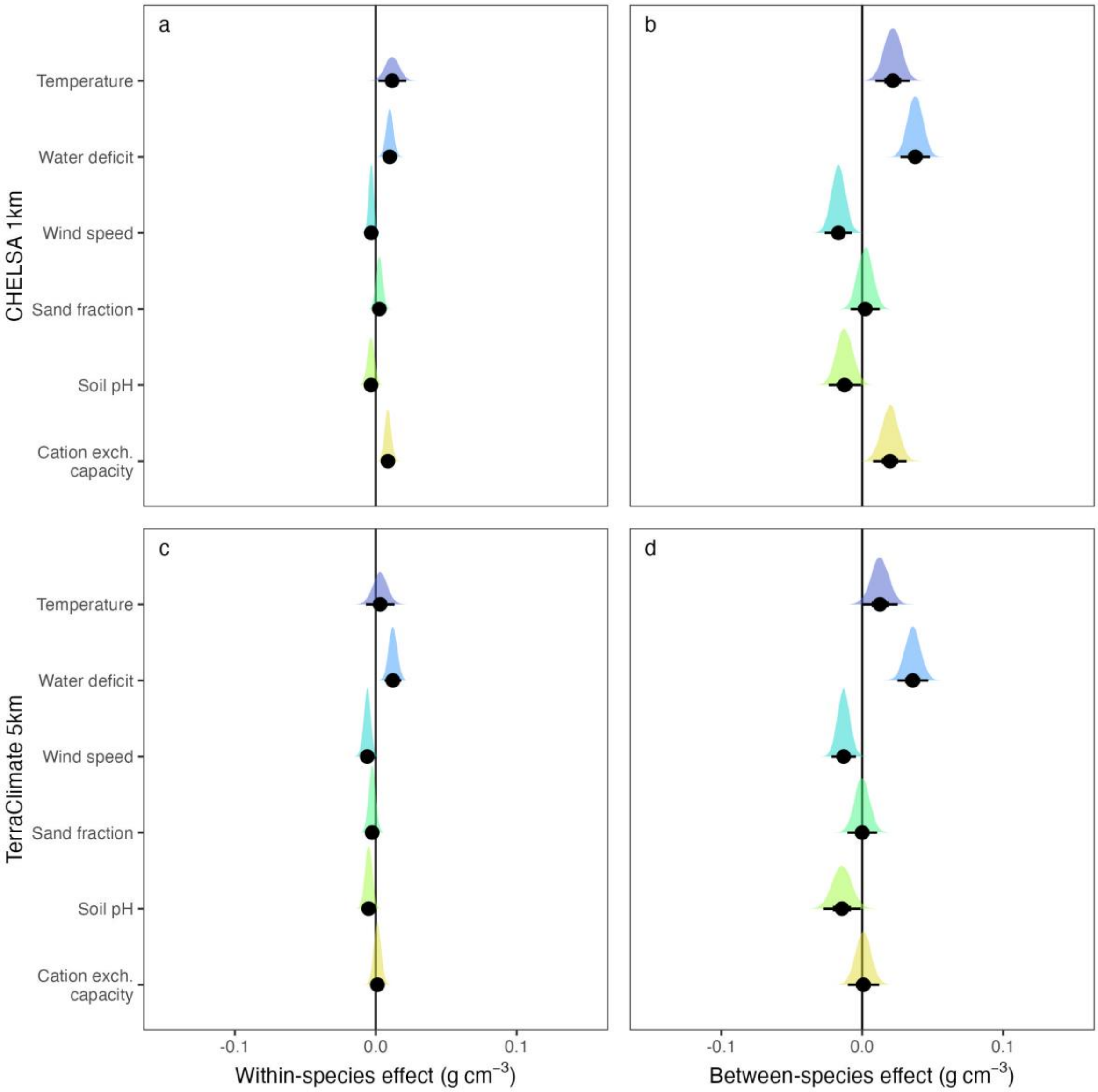

**Fig. S7: Environmental and edaphic effects on wood density.** Panels a) and b) are equivalent to panels a) and b) in Fig. 4 in the main text and show within-species and between-species effects of environmental variables on wood density, with slightly wider x axis for comparability with effect sizes in data subsets (cf. Fig. S9). Panels c) and d) are the equivalents of panels a) and b), but using the TerraClimate (Abatzoglou et al., 2018) climatology 1981-2010 instead of CHELSA (Brun et al., 2022; Karger et al., 2017). Variables are mean annual temperature ("Temperature", in °C), climatic water deficit ("Water deficit", in mm), and mean wind speed ("Wind speed", m s$^{-1}$). The *soilgrids* layers (Hengl et al., 2017) are the same, but have been extracted at 5 km to match the 4-5 km resolution of TerraClimate. The corresponding model results can be found in Table S10. Throughout, black dots indicate the median effect size and black intervals quantile ranges (66% and 95%).

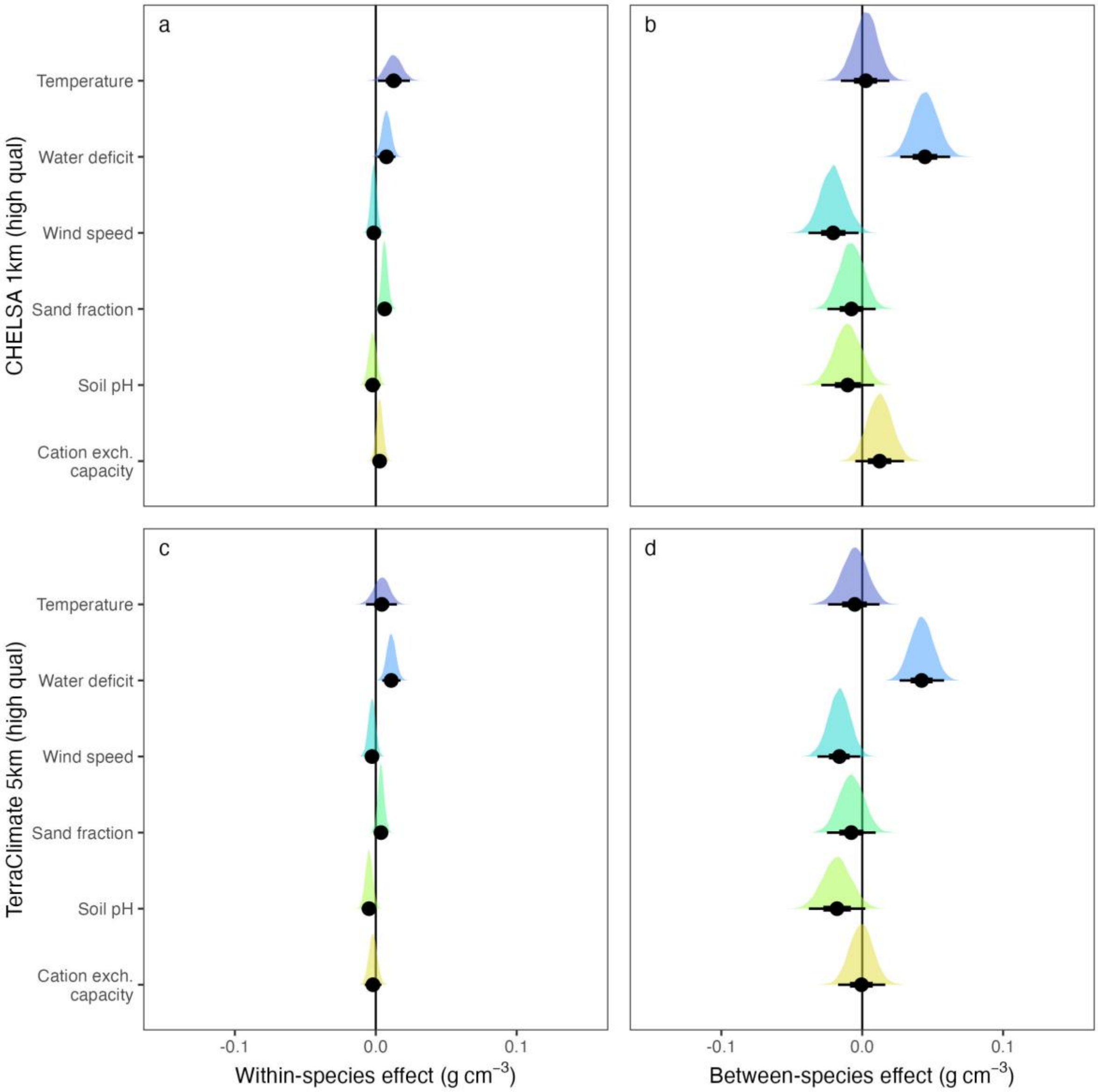

**Fig. S8: Environmental and edaphic effects on wood density, high-quality subset.** Same as Fig. S7, but for a higher-quality subset of the GWDD v.2 where each species varied strongly in at least environmental variable. This meant that a species was only included if the range of at least one of the environmental predictors was in the top 10% of ranges for that predictor among all other species. We note that this may slightly bias the data set towards the better-sampled higher latitude regions. The corresponding model results can be found in Table S10. Throughout, black dots indicate the median effect size and black intervals quantile ranges (66% and 95%).

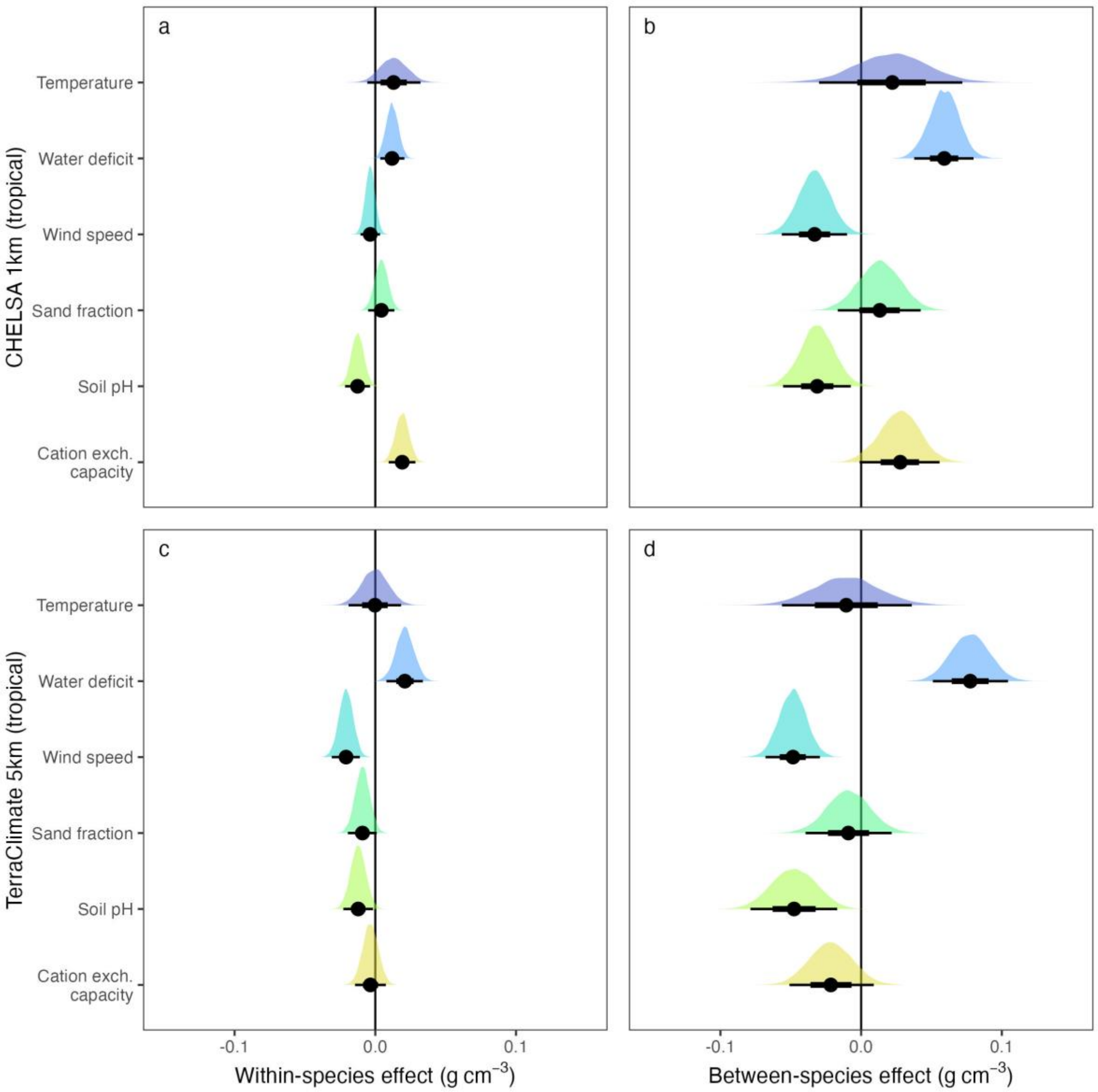

**Fig. S9: Environmental and edaphic effects on wood density, within tropics.** Same as Fig.S7-8, but for a subset of species with at least three locations in the tropics. Locations were defined as distinct 5 km grid cells. The corresponding model results can be found in Table S12. Throughout, black dots indicate the median effect size and black intervals quantile ranges (66% and 95%).

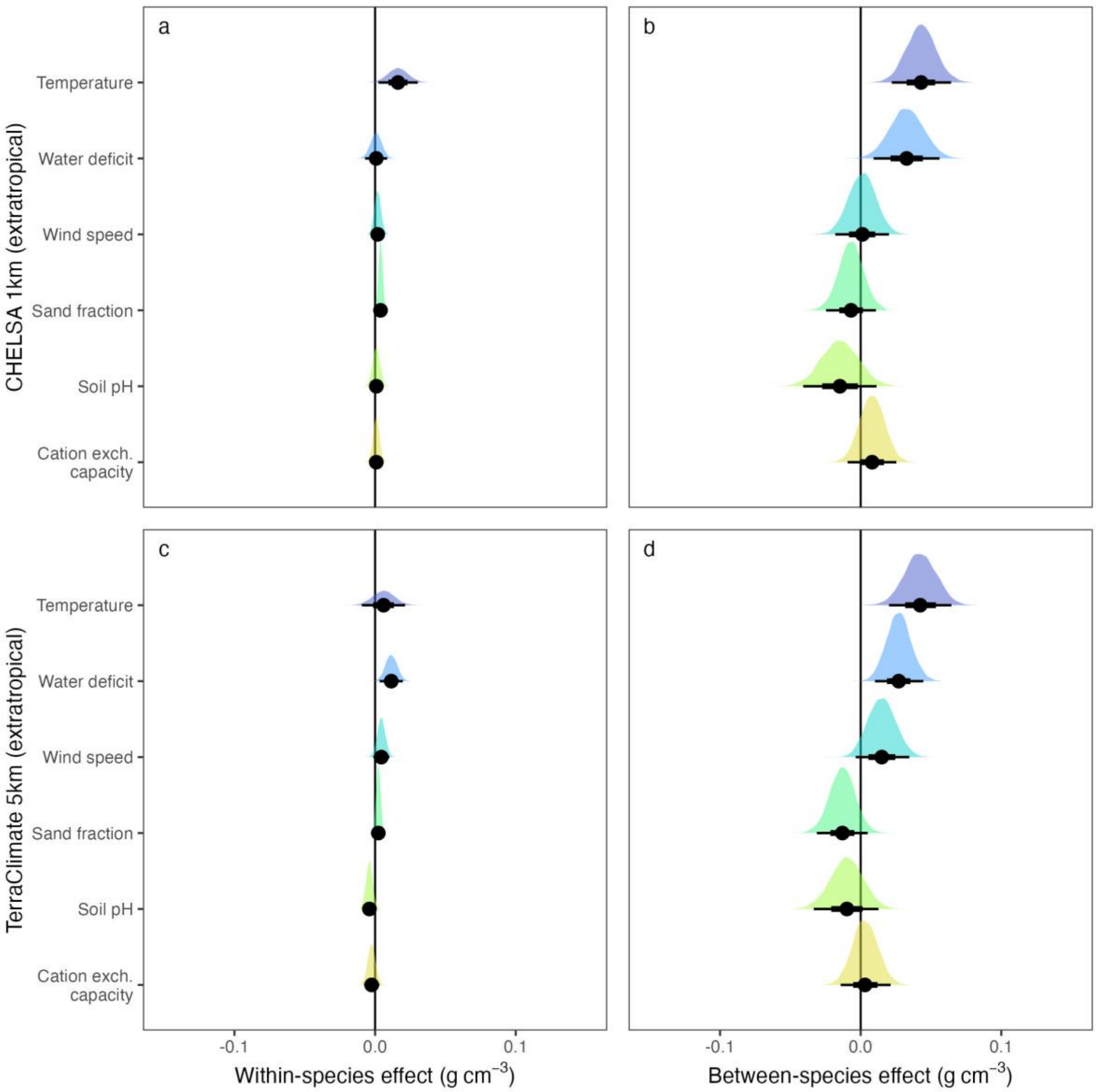

**Fig. S10: Environmental and edaphic effects on wood density, outside of tropics.** Same as Fig.S7-9, but for a subset of species with at least three locations outside of the tropics. Locations were defined as distinct 5 km grid cells. The corresponding model results can be found in Table S12. Throughout, black dots indicate the median effect size and black intervals quantile ranges (66% and 95%).

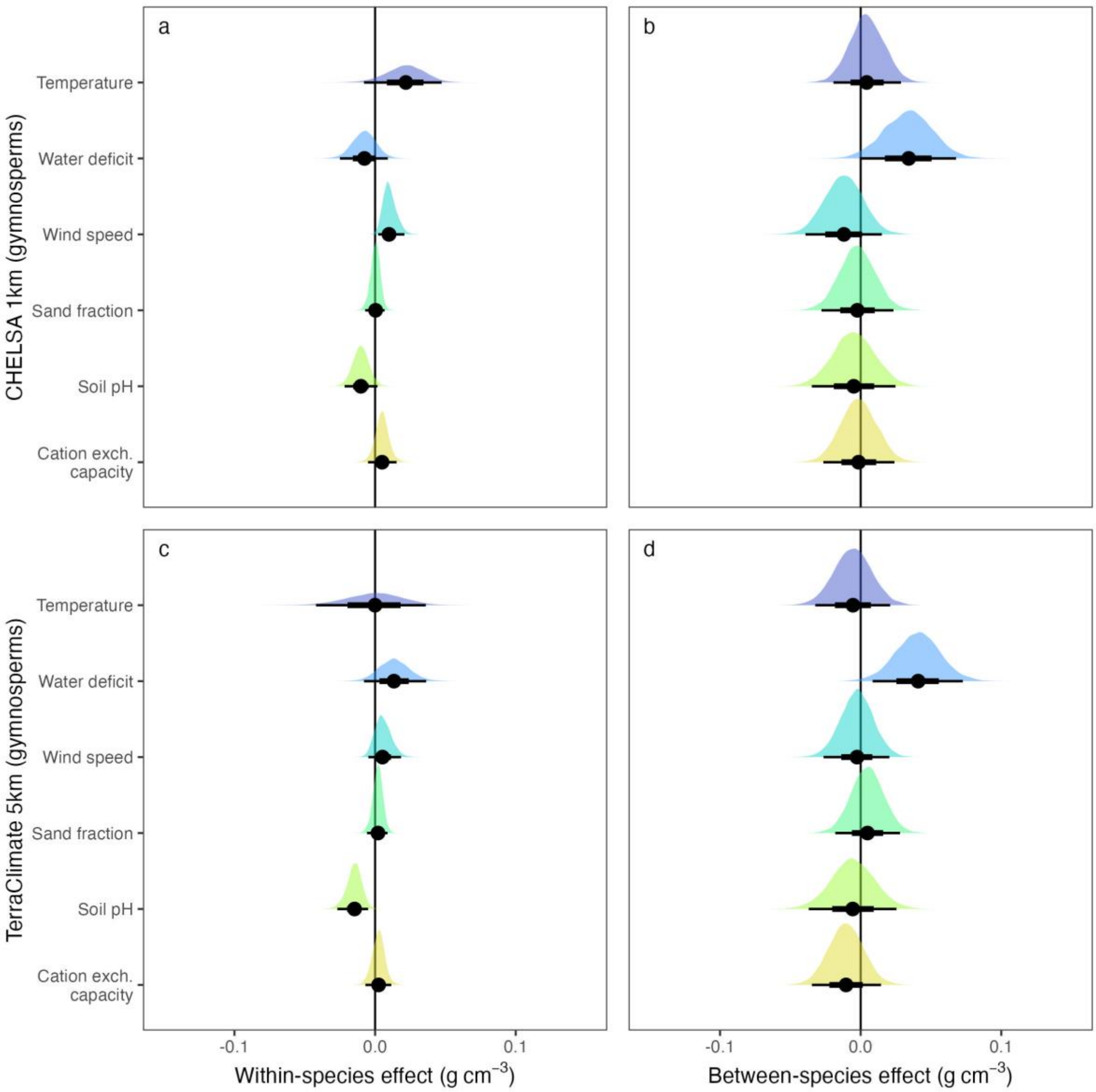

**Fig. S11: Environmental and edaphic effects on wood density in gymnosperms.** Same as Fig.S7-10, but for gymnosperms only. The corresponding model results can be found in Table S15. Throughout, black dots indicate the median effect size and black intervals quantile ranges (66% and 95%). Note how most effects are close to 0 or strongly overlap with 0.

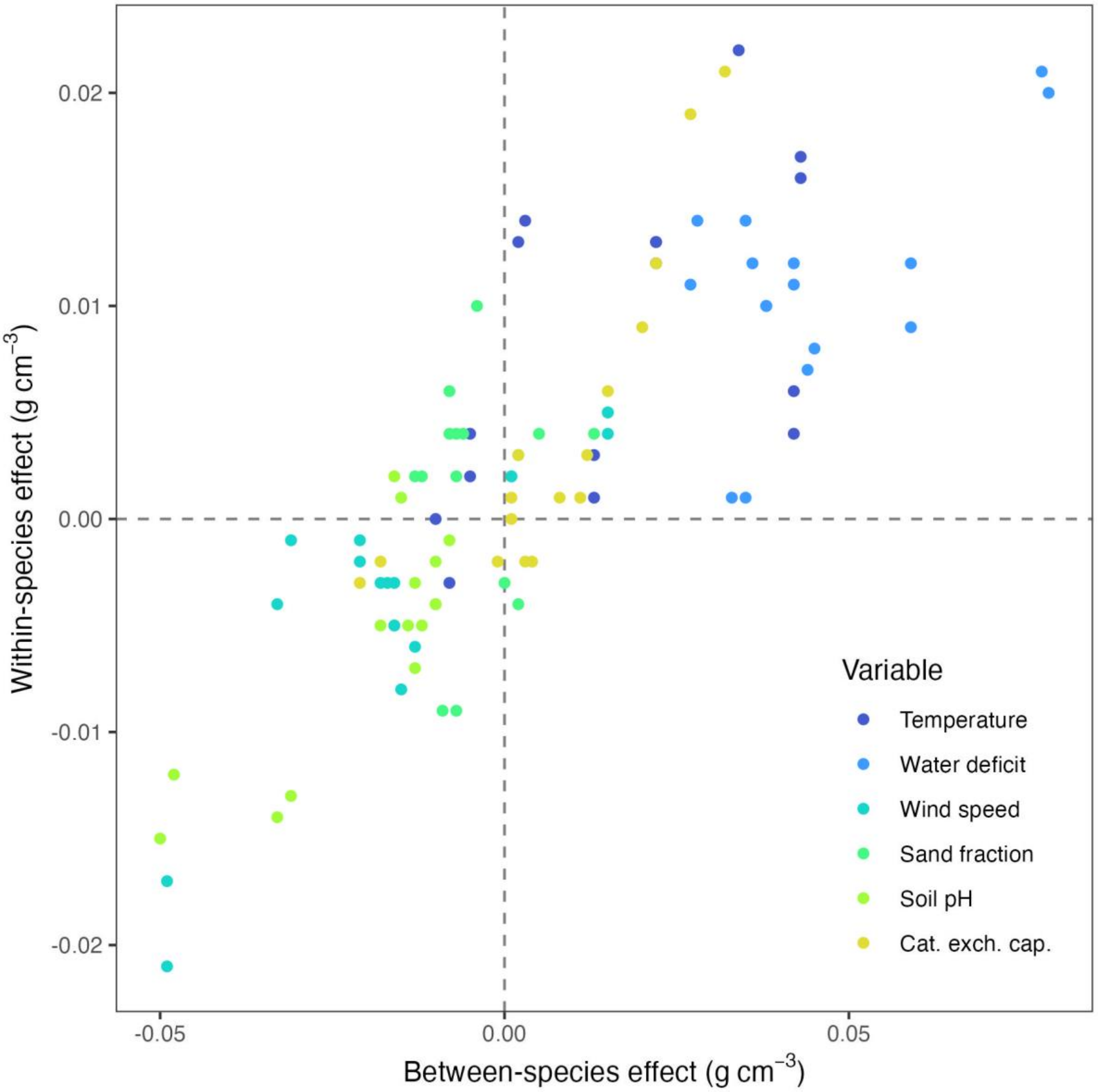

**Fig. S12: Correlation between within-species effects and between-species effects.** Shown are correlations between estimated within-species and between-species effects of environmental and edaphic predictors on wood density variation. Every point represents an estimated effect of one predictor from one of the 8 models we used (Table S3, Tables S10-13) and counting Bayesian and ML models separately (96 estimates overall). The overall correlation is r = 0.83. Note that this correlation between within-species and between-species effects is not due to correlation between environmental predictors within and across species, as these are decoupled by construction and also have strikingly different correlation structures (Fig. S1).

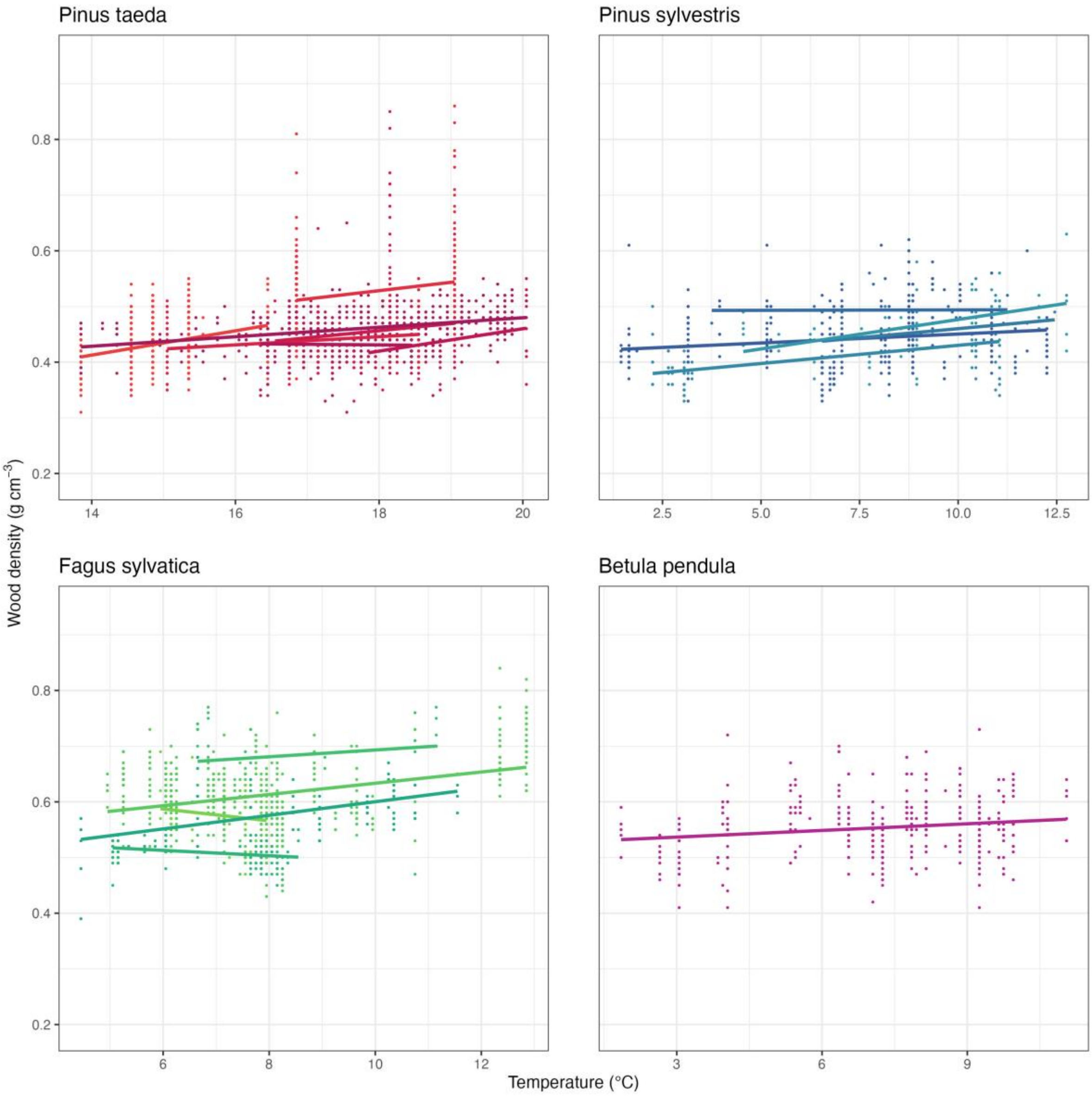

**Fig. S13: Temperature effects on within-species wood density variation, examples.** Shown are the within-species effects of temperature variation on wood density in four temperate species that cover large environmental gradients and are well-sampled (several studies or large sample sizes). Each line represents a different study and is fitted via simple OLS regression. Note how wood density seems to increase very slightly with temperature, but how this effect is overwhelmed by variation around the regression line and even differs between studies. Temperatures are based on the CHELSA data set as in Fig. 4.

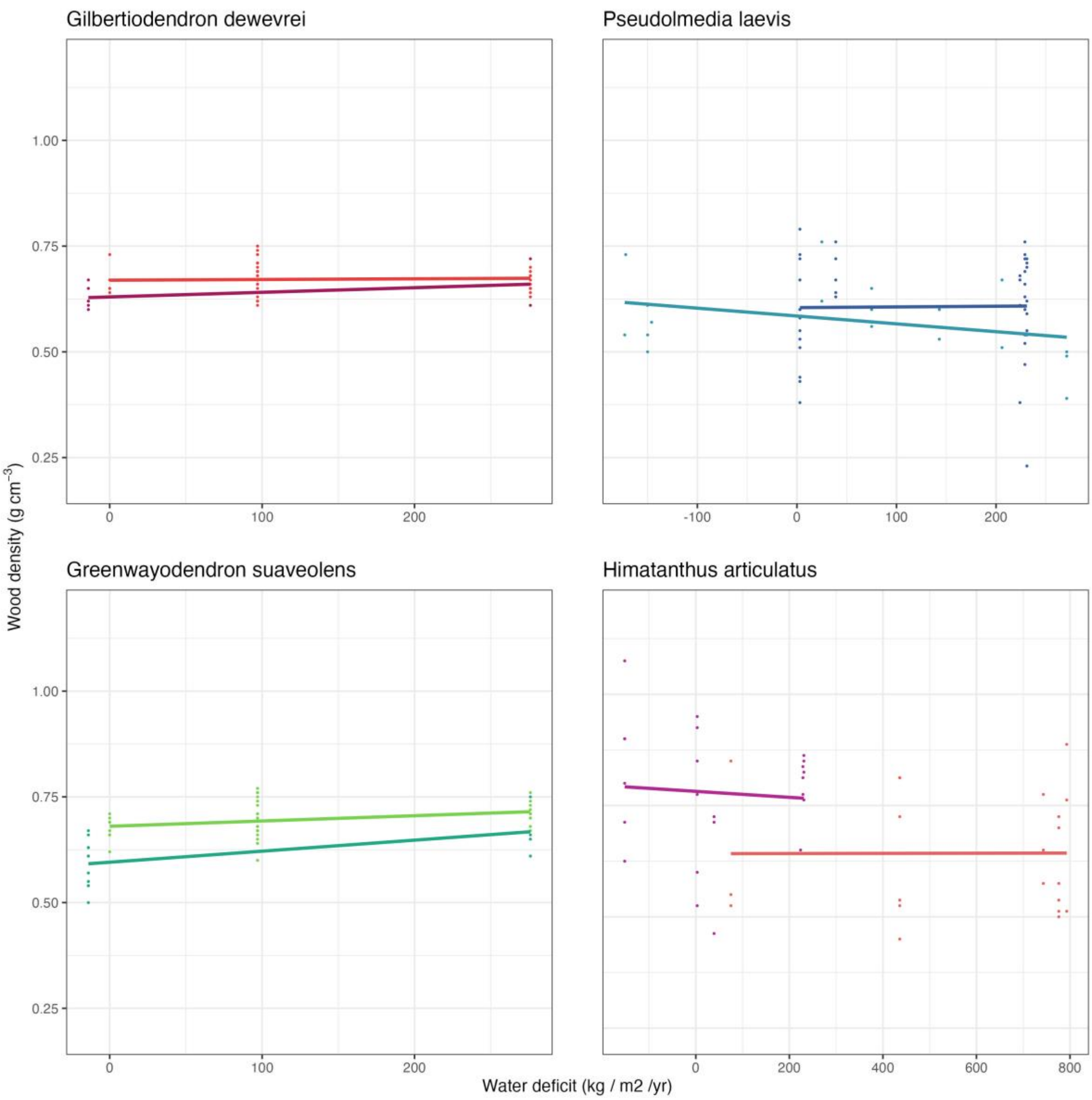

**Fig. S14: Water deficit effects on within-species wood density variation, examples.** Same as Fig. S11, but for water deficit. Shown are the within-species effects of water deficit on wood density in four tropical species that cover large environmental gradients and are well-sampled (two different studies for each). Each line represents a different study and is fitted via simple OLS regression. Note how there is no clear overall effect and how effect sizes are weak even across large gradients. Water deficits are based on the CHELSA data set as in Fig. 4.

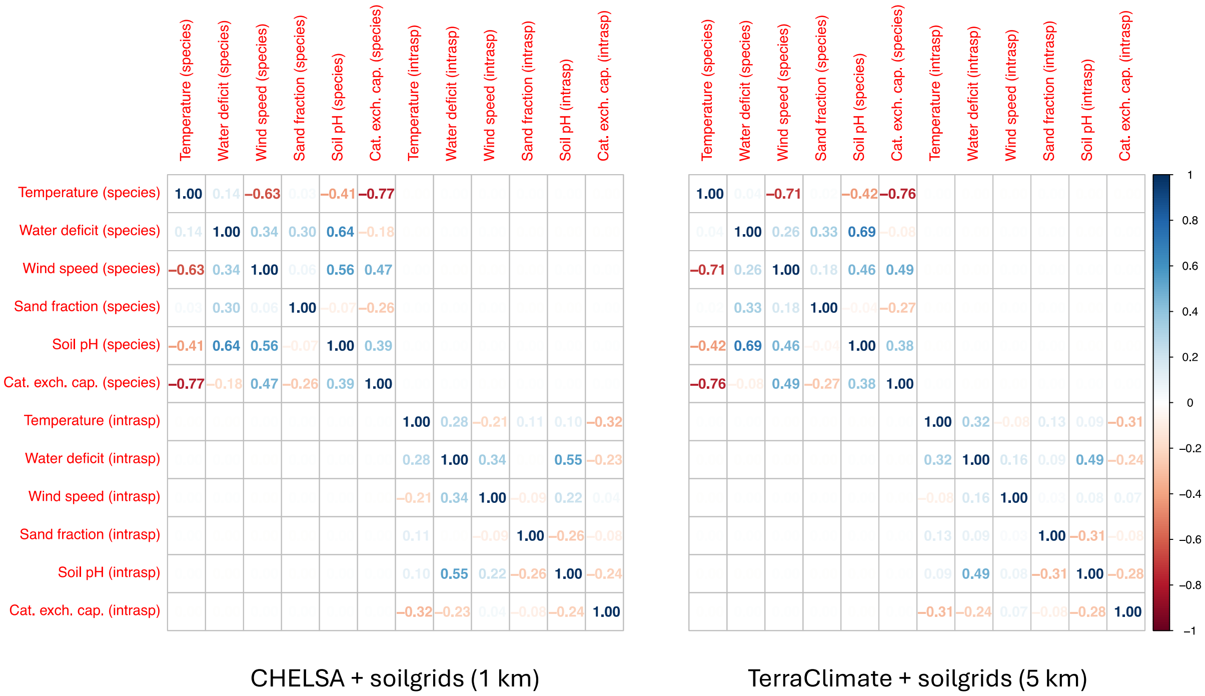

**Fig. S15: Correlation between environmental and edaphic predictors.** Shown are the correlations between environmental and edaphic predictors matched to the GWDD v.2. All predictors are separated into species mean values, summarizing the typical environment a species experiences ("species" in brackets), and deviations from the species means at the individual level, i.e., indicating within-species environmental variation ("intrasp" in brackets). Correlation matrices are shown for both sets of predictors used throughout the study, i.e. the CHELSA climatologies 1981-2010 (Brun et al., 2022; Karger et al., 2017) in conjunction with 1 km gridded *soilgrids* layers (Hengl et al., 2017), as well as TerraClimate climatologies 1981-2010 (Abatzoglou et al., 2018) in conjunction with 5 km gridded *soilgrids* layers. Note that, by construction, species means are fully decorrelated from within-species deviations from the species means ($r < 0.01$), hence large parts of the correlation matrices appear empty. Also note that, at species level, temperature is correlated in excess of $|r| = 0.7$ with one variable (cation exchange capacity) and has an absolute correlation of ~0.7 with one other variable (wind speed). Since we are predominately interested in within-species effects and not in exactly partitioning out species-level effects, we include all three variables in the model. A high absolute correlation, but below 0.7, also exists between soil pH and water deficit.
